# Increased serological response against human herpesvirus 6A is associated with risk for multiple sclerosis

**DOI:** 10.1101/737932

**Authors:** Elin Engdahl, Rasmus Gustafsson, Jesse Huang, Martin Biström, Izaura Lima Bomfim, Pernilla Stridh, Mohsen Khademi, Nicole Brenner, Julia Butt, Angelika Michel, Daniel Jons, Maria Hortlund, Lucia Alonso-Magdalena, Anna Karin Hedström, Louis Flamand, Masaru Ihira, Tetsushi Yoshikawa, Oluf Andersen, Jan A. Hillert, Lars Alfredsson, Tim Waterboer, Peter Sundström, Tomas Olsson, Ingrid Kockum, Anna Fogdell-Hahn

## Abstract

Human herpesvirus (HHV)-6A or HHV-6B involvement in multiple sclerosis (MS) etiology has remained controversial mainly due to the lack of serological methods that can distinguish the two viruses.

A novel multiplex serological assay measuring IgG reactivity against the immediate-early protein 1 from HHV-6A (IE1A) and HHV-6B (IE1B) was used in a MS cohort (8742 persons with MS and 7215 matched controls), and a pre-MS cohort (478 individuals and 476 matched controls) to investigate this further.

The IgG response against IE1A was positively associated with MS (OR = 1.55, p = 9×10^−22^), and increased risk of future MS (OR = 2.22, p = 2×10^−5^). An interaction was observed between IE1A and Epstein-Barr virus (EBV) antibody responses for MS risk (attributable proportion = 0.24, p = 6×10^−6^). In contrast, the IgG response against IE1B was negatively associated with MS (OR = 0.74, p = 6×10^−11^). The association did not differ between MS subtypes or vary with severity of disease. The genetic control of HHV-6A/B antibody responses were located to the Human Leukocyte Antigen (HLA) region and the strongest association for IE1A was the DRB1*13:01-DQA1*01:03-DQB1*06:03 haplotype while the main association for IE1B was DRB1*13:02-DQA1*01:02-DQB1*06:04.

In conclusion a role for HHV-6A in MS etiology is supported by an increased serological response against HHV-6A IE1 protein, an interaction with EBV, and an association to HLA genes.

## INTRODUCTION

Human herpesvirus 6A (HHV-6A) and HHV-6B are closely related *beta-herpesviruses* with distinct biological and immunological properties as well as differences in epidemiology and disease associations (1). HHV-6B is acquired early in life (2, 3), with the vast majority of children infected before the age of two. Primary HHV-6B infection results in roseola, a disease characterized by high fever, rashes and occasional febrile seizures (3–5). As with all herpesviruses, HHV-6A and HHV-6B can establish latency and reactivate later in life, which can lead to severe diseases such as encephalitis (reviewed in (6)). Less is known about any clinical manifestations of the primary infection of HHV-6A, but this virus has repeatedly been reported to be associated with multiple sclerosis (MS) (7–12). As previous studies have been limited in size or unable to separate the HHV-6A from B serologically, a more definite view on their respective roles in MS would benefit from a comprehensive population based case-control study on the diverging serological response against these two viruses.

MS is characterized by central nervous system inflammation and demyelination, with several different disease courses: relapsing remitting MS (RRMS), secondary progressive MS (SPMS), and primary progressive MS (PPMS). The etiology of the disease includes a genetic predisposition (13, 14). Lifestyle/environmental factors, like virus infections and smoking also play a role, and they often interact with MS risk genes (15). Among virus infections, the *gamma-herpesvirus* Epstein-Barr virus (EBV) has remained the strongest suspect in the MS etiology (16–21). Another *beta-herpesvirus*, cytomegalovirus (CMV), has through serological analysis been negatively associated with MS risk (22). We here explore the potential associations of HHV-6A and B in MS, and interaction with serological response to EBV and CMV, using serology applied to both a very large incident and prevalent MS case-control material, and importantly, also a pre-MS case-control cohort.

Seroconversion against HHV-6 usually occurs in early childhood (23–25) but as the two viruses have similar proteomes, it has been difficult to distinguish anti-HHV-6A from anti-HHV-6B antibody responses. This inability is a major concern when investigating virus-specific disease associations and a possible explanation for the contradictory associations between HHV-6 IgG response and MS (26–33). However, even though HHV-6A and HHV-6B are 90% homologous, there are parts of their genome with more divergence (34, 35). The immediate-early 1 (IE1) proteins (termed IE1A for HHV-6A and IE1B for HHV-6B), encoded by the open reading frame (ORF) U90-U89, are among the most divergent with only 62% homology (36, 37) and with differences in biological properties. IE1A, but not IE1B, can transactivate several heterologous promoters (37, 38) while IE1B, but not IE1A can silence IFN-α/β signaling (39). The ORF U11 coding for p100 in HHV-6A and 101K in HHV-6B also exhibit relatively high divergence with only 81% amino acid identity (40). These structural proteins are essential for viral growth and propagation (41), and 101K has been identified as the dominant antigen recognized by anti-HHV-6B IgG (42). With the aim to discriminate between IgG responses against these two viruses, we developed a novel bead-based multiplex serology assay measuring IgG antibodies against IE1A, IE1B, p100 and 101K, selecting the most divergent parts of these protein sequences. This assay was used to screen serum or plasma samples from persons with MS, persons that later develop MS, and controls for HHV-6A and HHV-6B protein-specific antibodies.

## RESULTS

### High IE1A antibody response is positively associated with MS and is a risk factor for developing MS in youth

We used a novel multiplex serology assay to investigate the specific IgG responses against the HHV-6A protein IE1A and the HHV-6B protein IE1B. To investigate if persons with MS and controls differed in IgG responses, logistic regression analyses were used to compare strong and weak responders, defined as the highest or lowest quartile of each measured response. This revealed that a high IE1A antibody response was positively associated with MS (OR = 1.55, p = 9×10^−22^) while a high IE1B antibody response was negatively associated with MS (OR = 0.74, p = 6×10^−11^) (**Table 1, Figures 1A** and **1B**).

**FIGURE 1.**
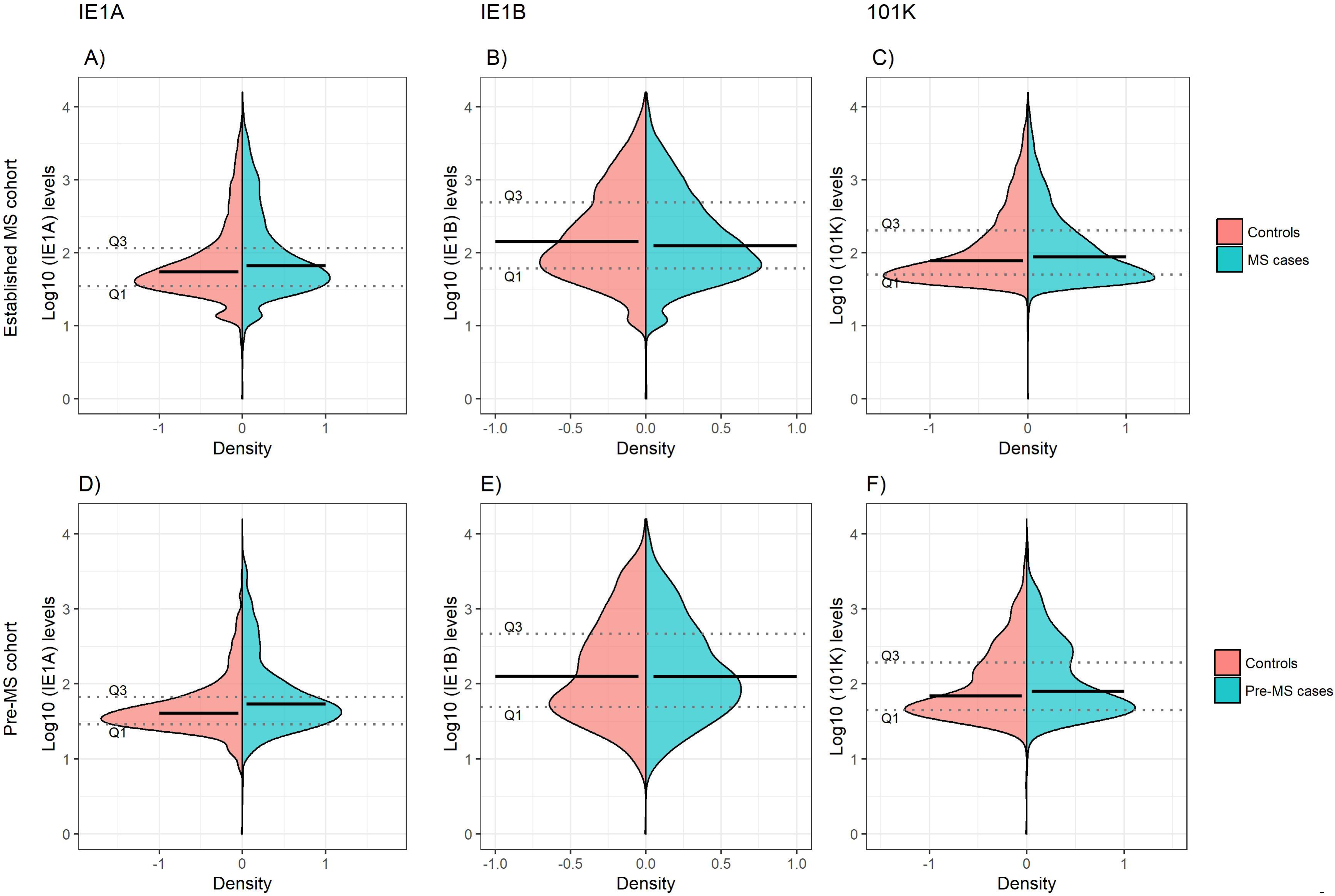
Antibody responses against HHV-6A and 6B proteins in MS cases and controls. Log10-transformed antibody levels measured as median fluorescence intensity (MFI) are visualized with bean plots for established MS cohort (n=8742 persons with MS (blue) and n=7215 controls (pink)) (A, B, C) and pre-MS cohort (n=478 persons with MS (blue) and n=476 controls (pink)) (D, E, F) for HHV-6A IE1A IgG (A and D); HHV-6B IE1B IgG (B and E); HHV-6B anti-101K IgG (C and F). The 1^st^ and 3^rd^ quartiles are indicated with dotted lines and solid lines indicate median.

**Table 1.**
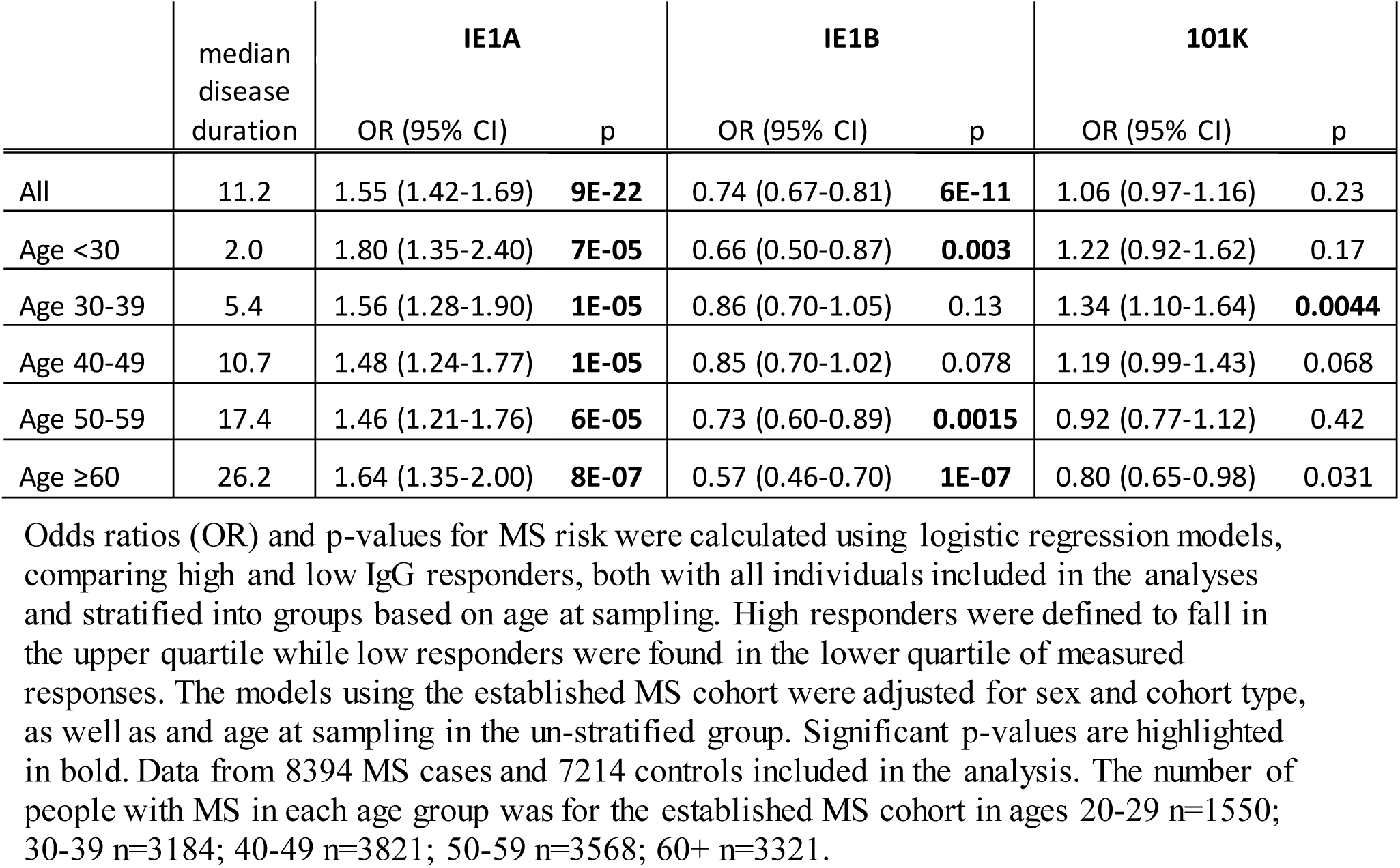
Association of IE1A, IE1B and 101K antibody response to MS in established MS cohort.

To investigate if these differences also were present before MS onset, serum samples drawn from persons with RRMS at a median of 8.3 years before symptom onset and from matched controls were analyzed. Strong IE1A responders had a higher risk of developing MS later in life, compared to low responders (OR = 2.22, p = 2×10^−5^) (**Table 2, Figure 1D**). No significant difference in IE1B IgG response was observed before MS onset (OR = 0.96, p = 0.8) (**Table 2** and **Figure 1E**).

The pre-MS cohort was divided into age groups and further analyzed (**Table 2**). In individuals younger than 20 years old, strong IE1A responders had a 3.38 (p = 0.004) times higher risk of developing MS later in life. This MS risk decreased with age, reaching an OR of 1.51 (p = 0.3) in the oldest age group (30-39 years). Also in the established MS cohort the highest OR was seen in the youngest age group (**Table 1**).

**Table 2.** Association of IE1A, IE1B and 101K antibody response to MS in pre-MS cohort HHV-6A in MS.

|  | median<br>disease<br>duration | IE1A |  | IE1B |  | 101K |  |
| --- | --- | --- | --- | --- | --- | --- | --- |
|  |  | OR (95% CI) | p | OR (95% CI) | p | OR (95% CI) | p |
| All | -8.3 | 2.22 (1.54-3.19) | **2E-05** | 0.96 (0.66-1.38) | 0.81 | 1.51 (1.06-2.17) | 0.024 |
| Age <20 | -9.8 | 3.38 (1.46-7.81) | **0.004** | 1.09 (0.46-2.57) | 0.84 | 1.36 (0.58-3.15) | 0.48 |
| Age 20-29 | -8.4 | 2.29 (1.43-3.67) | **0.001** | 0.88 (0.54-1.44) | 0.62 | 1.60 (1.00-2.54) | 0.048 |
| Age 30-39 | -5.7 | 1.51 (0.65-3.49) | 0.33 | 1.09 (0.51-2.35) | 0.83 | 1.42 (0.64-3.14) | 0.38 |

Odds ratios (OR) and p-values for MS risk were calculated using logistic regression models, comparing high and low IgG responders, both with all individuals included in the analyses and stratified into groups based on age at sampling. High responders were defined to fall in the upper quartile while low responders were found in the lower quartile of measured responses. The pre-MS cohort analyses are adjusted for sex, and age. Significant p-values are highlighted in bold. Data from 478 individuals who later developed MS and 476 matched controls included in the analysis.

### Interaction between high IE1A and EBV antibody responses on MS risk

As antibody responses against other herpesviruses such as EBV and CMV have been associated with MS (19, 22), the interplay between HHV-6A/-6B and these viruses in MS was analyzed. The median antibody levels of EBV and CMV index among controls were used as cutoffs for strong or weak responders. Interaction analyses revealed a significant additive interaction between IE1A and EBV responses on MS risk (attributable proportion due to interaction (AP = 0.24, p = 6×10^−6^, **Figure 2A**), meaning that 24% of the risk for developing MS in those with strong IE1A and strong EBV responses was due to interaction between these factors. No significant interaction between HHV-6A and CMV immune response was observed, neither was the HHV-6B IE1B or 101K responses interacting with EBV on MS risk (**Figure 2B**, SI appendix Figure S1A and B). An analysis of how the OR varied for the three HHV-6A and B antigens depending on EBV index was investigated using a sliding window approach and shows that IE1A mediated risk for MS is limited to individuals with EBV response higher than the median among controls (SI appendix Figure S2).

**FIGURE 2.**
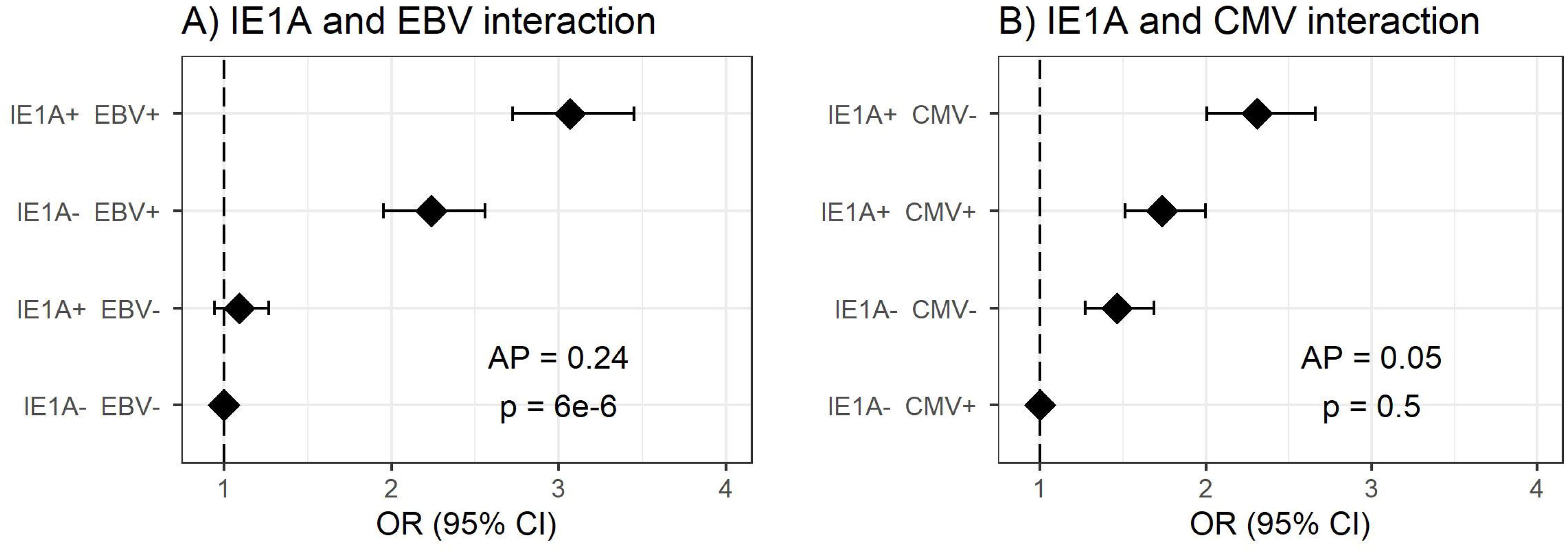
Interaction of antibody response against different herpesviruses in association to MS. Odds ratios (OR) and confidence intervals (CI) for IE1A and EBV (A) and IE1A and CMV (B), were obtained through logistic regression models adjusted for age, sex and cohort type analyzing the Established MS cohort (n=8742 persons with MS and n=7215 controls). OR were calculated in relation to the group with the lowest MS risk. Plus (+) indicates being a strong responder while minus (-) indicates being a weak responder. Strong IE1A response is defined as having an MFI value being in the upper quartile of measured response, while a low response is having an antibody measurement being in the lower quartile of measured response. Strong EBV / CMV response is defined as having a higher EBV / CMV index than the median among controls, while a weak response is having a lower index compared to the median among controls.

### Antibody levels against IE1A are higher in both established MS cases and pre-MS cases, compared to matched controls

Linear regression models were used to analyze the HHV-6A/6B IgG levels, and the results are in line with the association seen for high/low serological response. MS cases, both before (pre-MS) and after MS onset (established MS), had higher IgG levels against IE1A than controls (p = 6×10^−10^ and p = 9×10^−30^, respectively; **Figure 1**, Table S1 and S2 in Supplementary Material). In contrast, anti-IE1B IgG levels were lower in established MS cases compared to controls (p = 9×10^−14^), but this association was not observed before MS onset. When dividing the study cohorts into age groups, IE1A reactivity was consistently higher in MS cases compared to in controls (**Figure 3A** and **B**), while the pattern for the anti-IE1B reactivity was more inconsistent (Figure S3A-D in Supplementary Material). The associations of IE1A with MS were significant both with and without adjustment for EBV and CMV responses, indicating that the increased IE1A response in MS was not confounded by these two anti-viral responses (Table S1 and S2 in Supplementary Material). In addition to the IE proteins, antibodies against the structural protein 101K (HHV-6B) and p100 (HHV-6B) were measured. A high 101K serological response was not associated with MS or with later development of MS in the pre-MS cohort (**Table 1** and **2**). Results from the p100 analysis were excluded from further analyses due to the low reactivity against this antigen.

**FIGURE 3.**
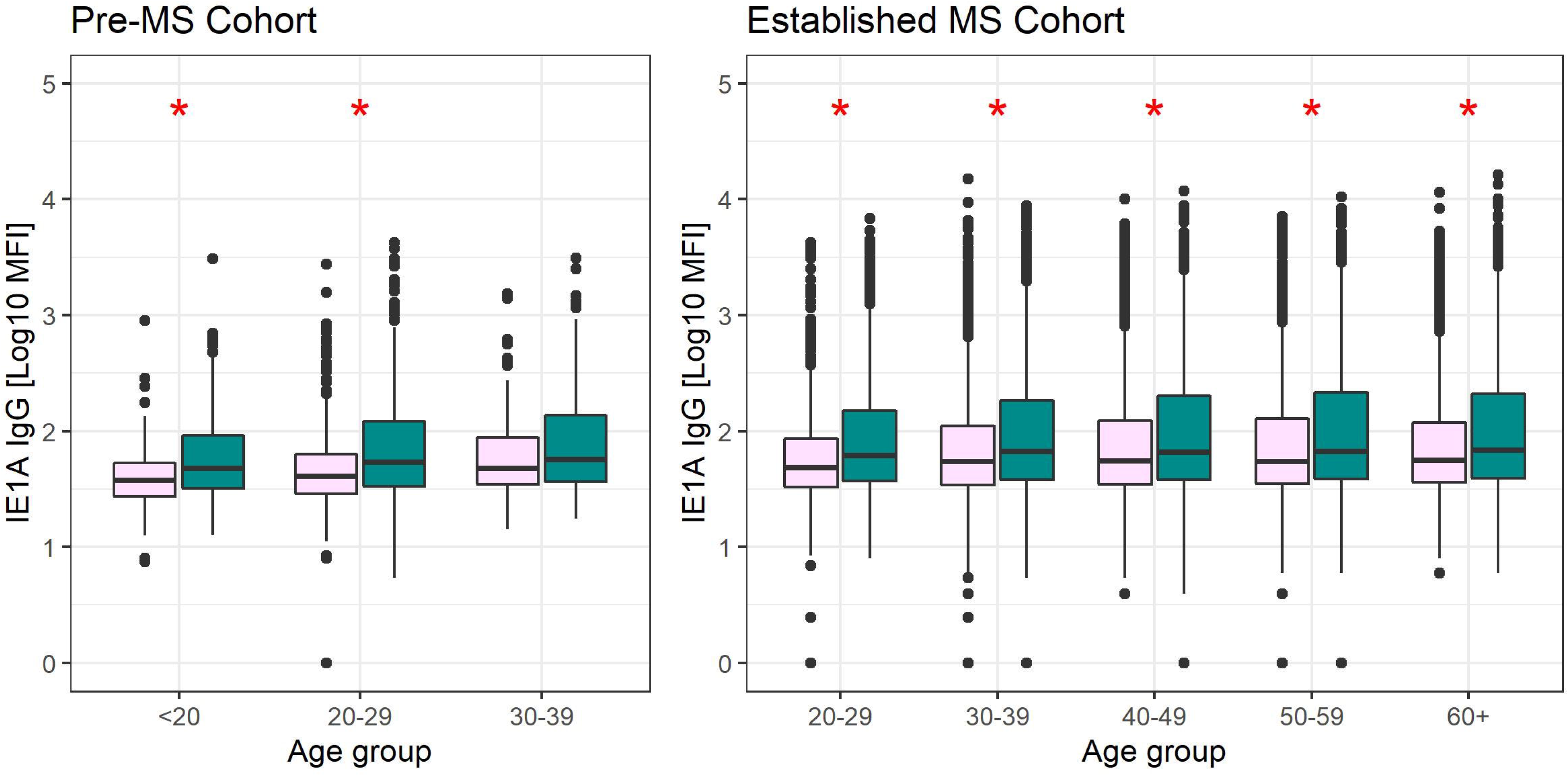
Median MFI response against HHV-6A IE1A protein in different age groups. Median of median fluorescence intensity (MFI) in different age groups for A) pre-MS cohort (n=478 persons who later developed MS (green) and n=476 controls (pink)) and B) established MS cohort (n=8394 persons with MS (green) and n=7214 controls (pink)). Statistics were calculated with linear regression. Significant (p<0.008) differences in IgG levels between MS cases and controls within each age group are indicated with *.

### High IE1A response is associated with relapsing and progressive MS, but not disease severity

The association of high IE1A responses was similar regardless of disease course (OR_RRMS_ = 1.62, p = 1×10^−20^; OR_SPMS_ = 1.49, p = 7×10^−7^; OR_PPMS_ = 1.53, p = 9×10^−4^). The same was true for high IE1B responders (OR_RRMS_ = 0.77, p = 2×10^−6^; OR_SPMS_ = 0.67, p = 2×10^−6^; OR_PPMS_ = 0.62, p = 7×10^−4^). A high 101K response was associated with RRMS (OR = 1.18, p = 1×10^−3^), but negatively associated with PPMS (OR = 0.66, p = 2×10^−3^).

HHV-6A and 6B serology was not associated with two MS severity scores, the Multiple Sclerosis Severity Score (MSSS) and the Age Related Multiple Sclerosis Severity Score (ARMSS) (43) (data not shown).

### The IgG responses against HHV-6B proteins vary with age and sex

The level of IgG responses against IE1B and 101K decreased with age (p = 4×10^−21^ and p = 3×10^−39^, respectively; Figure S3A-D in Supplementary Material). A sex difference could be observed, with women eliciting a significantly stronger antibody response against IE1B (p = 4×10^−4^) and 101K (p = 2×10^−24^). Antibody levels against IE1A did not differ significantly between the sexes nor with age.

### Smoking associates with increased IE1A IgG response in persons with MS

As smoking has been reported to be a risk factor for MS disease (44) and has been associated with higher HHV-6 IgG levels (32), the effect of smoking on HHV-6 protein-specific IgG responses was investigated. Persons with MS and with a history of regular smoking showed higher IE1A IgG levels compared to those who never smoked (p = 2×10^−5^). This was not observed in controls (p = 0.4). The responses against the other proteins were not affected by smoking, neither in MS cases nor in controls (data not shown).

### IgG responses against HHV-6 proteins are associated with different HLA haplotypes

To investigate the influence of genetic factors for the serological response against the HHV-6A and HHV-6B protein sequences, genome-wide association studies (GWAS) were performed for both IgG levels (**Figure 4**) and high/low response (Figure S4 in Supplementary Material). The primary genetic association was mapped to the Human Leukocyte Antigen (HLA) region (6p21). There was a clear difference in IE1A and IE1B response in regard to their association to SNPs located in the HLA region, where IE1A levels were associated (p < 5×10^−8^) with 191 SNPs mapping to the HLA region while IE1B IgG levels were significantly associated with only two SNPs in this region.

**FIGURE 4.**
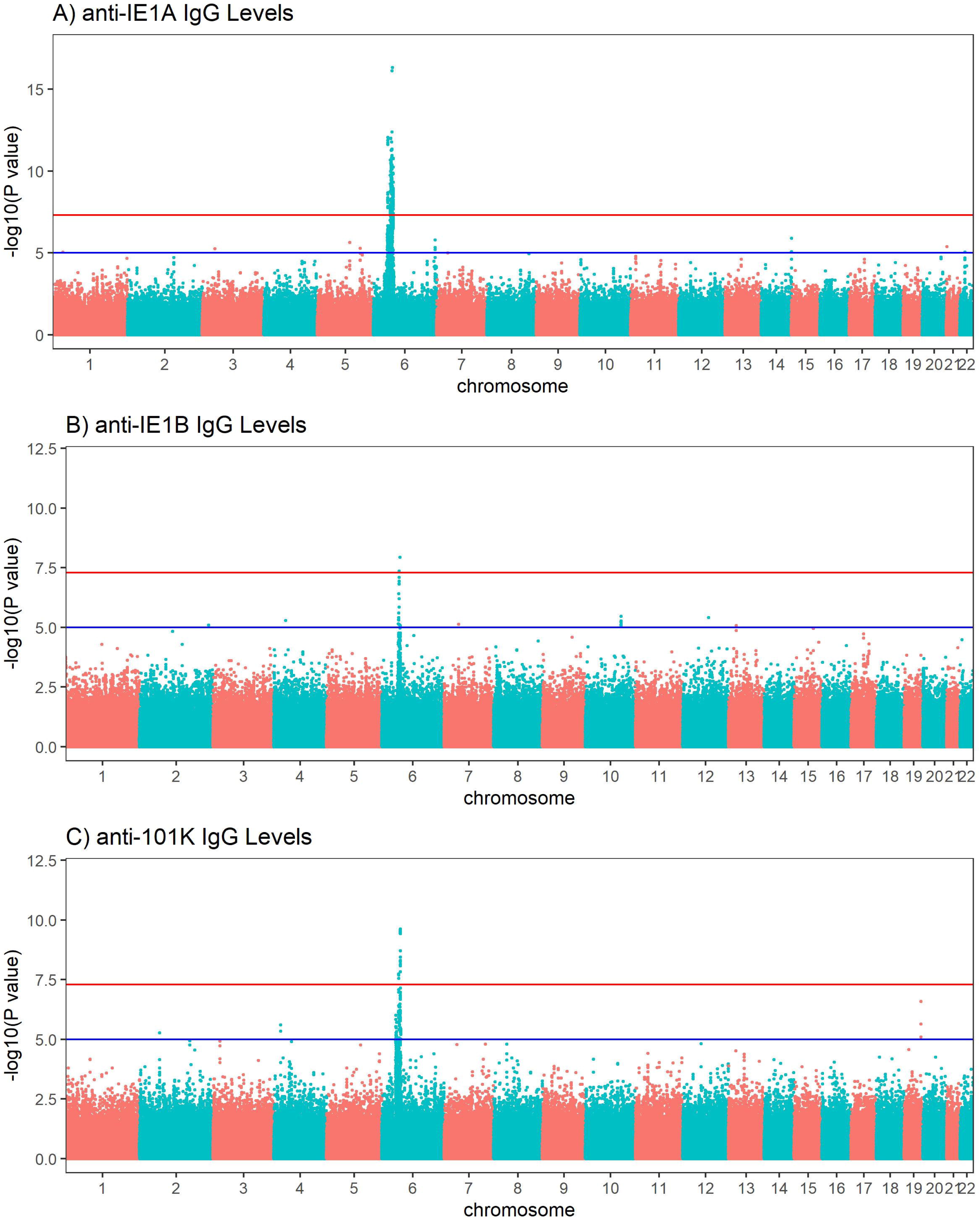
Manhattan plots visualizing associations between SNPs and anti-HHV-6A/6B protein IgG response levels. GWAS data (n=6,396 MS cases and n=5,530 controls from the established MS cohort) obtained through linear regression models showing associations between SNPs and IgG response (Log10 levels) against (A) IE1A, (B) IE1B and (C) 101K. Red lines indicate GWAS significance level of 5×10^−8^ (-log10 = 7.3 on the y-axis) and blue lines indicate suggestive association (p=10^−5^). Analysis was carried out jointly in MS cases and controls and adjusted with age, sex, cohort type and case status.

Deciphering of the associations within the HLA region showed that IgG responses against the three different protein sequences were associated with different HLA haplotypes (**Table 3**, Tables S3 and S4 in Supplementary Material). There were some differences between cases and controls, but for most haplotypes the association was stronger when cases and controls were analyzed together (**Table 3**, Tables S3 and S4 in Supplementary Material). Several HLA alleles exhibited significant association with IE1A response at the cut-point level for GWAS (p < 5×10^−8^). Higher IE1A IgG levels were associated with carrying the DRB1*13:01-DQA1*01:03-DQB1*06:03 haplotype in both MS cases and controls also after adjustments for HLA. The DPA1*02:01-DPB1*01:01 and the DRB1*04:01-DQA1*03-DQB1*03:02 haplotypes were associated with a lower IE1A response in both MS cases and all subjects. These associations where also still significant after adjustment for HLA (**Table 3**). The main HLA association for IE1B was DRB1*13:02-DQA1*01:02-DQB1*06:04, although none of the HLA associations reached the genome wide significance of p < 5×10^−8^ (Table S3 in Supplementary Material). Anti-101K response showed associations with several SNPs in the HLA region, but for the individual HLA haplotypes detected, none of them reached genome wide significance of p < 5×10^−8^ (Table S4 in Supplementary Material).

**Table 3.**
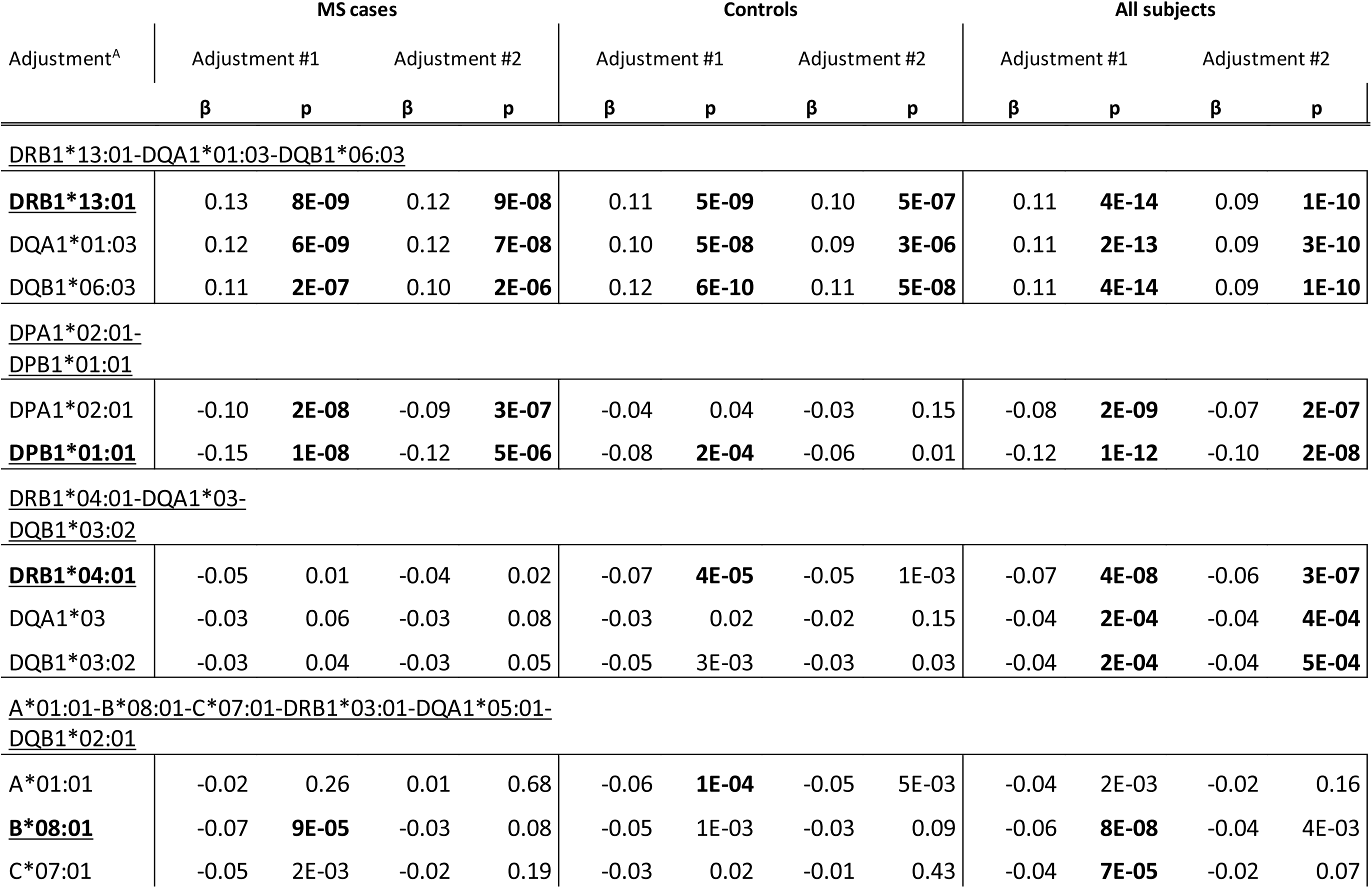
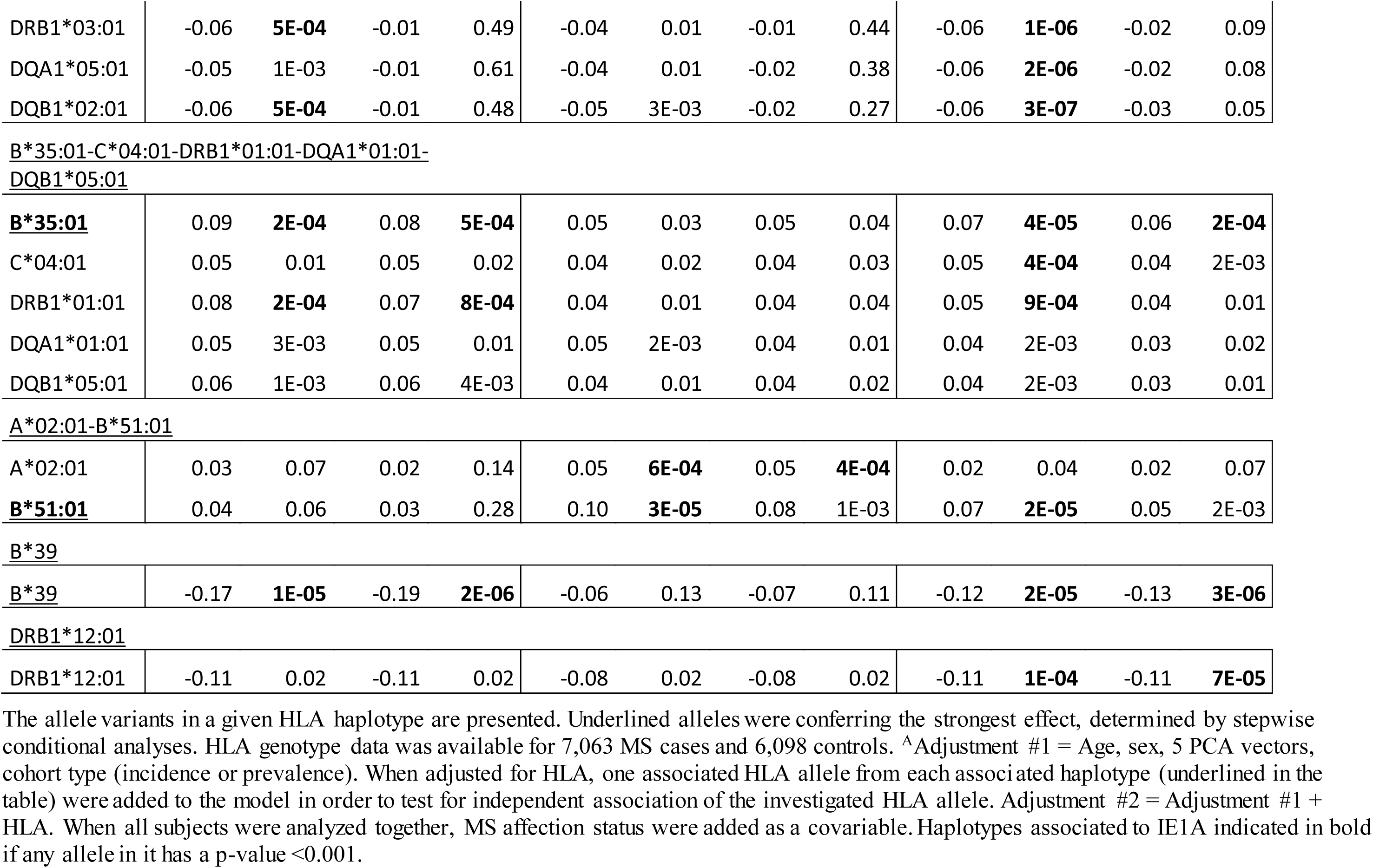
Association between IgG levels and HLA haplotypes and IE1A.

| Adjustment<sup>A</sup> | MS cases |  |  |  | Controls |  |  |  | All subjects |  |  |  |
| --- | --- | --- | --- | --- | --- | --- | --- | --- | --- | --- | --- | --- |
|  | Adjustment #1 |  | Adjustment #2 |  | Adjustment #1 |  | Adjustment #2 |  | Adjustment #1 |  | Adjustment #2 |  |
| | $\beta$ | p | $\beta$ | p | $\beta$ | p | $\beta$ | p | $\beta$ | p | $\beta$ | p |
| <u>DRB1*13:01-DQA1*01:03-DQB1*06:03</u> |  |  |  |  |  |  |  |  |  |  |  |  |
| **<u>DRB1\*13:01</u>** | 0.13 | **8E-09** | 0.12 | **9E-08** | 0.11 | **5E-09** | 0.10 | **5E-07** | 0.11 | **4E-14** | 0.09 | **1E-10** |
| DQA1*01:03 | 0.12 | **6E-09** | 0.12 | **7E-08** | 0.10 | **5E-08** | 0.09 | **3E-06** | 0.11 | **2E-13** | 0.09 | **3E-10** |
| DQB1*06:03 | 0.11 | **2E-07** | 0.10 | **2E-06** | 0.12 | **6E-10** | 0.11 | **5E-08** | 0.11 | **4E-14** | 0.09 | **1E-10** |
| <u>DPA1*02:01-DPB1*01:01</u> |  |  |  |  |  |  |  |  |  |  |  |  |
| DPA1*02:01 | -0.10 | **2E-08** | -0.09 | **3E-07** | -0.04 | 0.04 | -0.03 | 0.15 | -0.08 | **2E-09** | -0.07 | **2E-07** |
| **<u>DPB1\*01:01</u>** | -0.15 | **1E-08** | -0.12 | **5E-06** | -0.08 | **2E-04** | -0.06 | 0.01 | -0.12 | **1E-12** | -0.10 | **2E-08** |
| <u>DRB1*04:01-DQA1*03-DQB1*03:02</u> |  |  |  |  |  |  |  |  |  |  |  |  |
| **<u>DRB1\*04:01</u>** | -0.05 | 0.01 | -0.04 | 0.02 | -0.07 | **4E-05** | -0.05 | 1E-03 | -0.07 | **4E-08** | -0.06 | **3E-07** |
| DQA1*03 | -0.03 | 0.06 | -0.03 | 0.08 | -0.03 | 0.02 | -0.02 | 0.15 | -0.04 | **2E-04** | -0.04 | **4E-04** |
| DQB1*03:02 | -0.03 | 0.04 | -0.03 | 0.05 | -0.05 | 3E-03 | -0.03 | 0.03 | -0.04 | **2E-04** | -0.04 | **5E-04** |
| <u>A*01:01-B*08:01-C*07:01-DRB1*03:01-DQA1*05:01-DQB1*02:01</u> |  |  |  |  |  |  |  |  |  |  |  |  |
| A*01:01 | -0.02 | 0.26 | 0.01 | 0.68 | -0.06 | **1E-04** | -0.05 | 5E-03 | -0.04 | 2E-03 | -0.02 | 0.16 |
| **<u>B\*08:01</u>** | -0.07 | **9E-05** | -0.03 | 0.08 | -0.05 | 1E-03 | -0.03 | 0.09 | -0.06 | **8E-08** | -0.04 | 4E-03 |
| C*07:01 | -0.05 | 2E-03 | -0.02 | 0.19 | -0.03 | 0.02 | -0.01 | 0.43 | -0.04 | **7E-05** | -0.02 | 0.07 |
| DRB1*03:01 | -0.06 | **5E-04** | -0.01 | 0.49 | -0.04 | 0.01 | -0.01 | 0.44 | -0.06 | **1E-06** | -0.02 | 0.09 |
| DQA1*05:01 | -0.05 | 1E-03 | -0.01 | 0.61 | -0.04 | 0.01 | -0.02 | 0.38 | -0.06 | **2E-06** | -0.02 | 0.08 |
| DQB1*02:01 | -0.06 | **5E-04** | -0.01 | 0.48 | -0.05 | 3E-03 | -0.02 | 0.27 | -0.06 | **3E-07** | -0.03 | 0.05 |
| **<u>B\*35:01</u>** | 0.09 | **2E-04** | 0.08 | **5E-04** | 0.05 | 0.03 | 0.05 | 0.04 | 0.07 | **4E-05** | 0.06 | **2E-04** |
| C*04:01 | 0.05 | 0.01 | 0.05 | 0.02 | 0.04 | 0.02 | 0.04 | 0.03 | 0.05 | **4E-04** | 0.04 | 2E-03 |
| DRB1*01:01 | 0.08 | **2E-04** | 0.07 | **8E-04** | 0.04 | 0.01 | 0.04 | 0.04 | 0.05 | **9E-04** | 0.04 | 0.01 |
| DQA1*01:01 | 0.05 | 3E-03 | 0.05 | 0.01 | 0.05 | 2E-03 | 0.04 | 0.01 | 0.04 | 2E-03 | 0.03 | 0.02 |
| DQB1*05:01 | 0.06 | 1E-03 | 0.06 | 4E-03 | 0.04 | 0.01 | 0.04 | 0.02 | 0.04 | 2E-03 | 0.03 | 0.01 |
| A\*02:01-B\*51:01 |  |  |  |  |  |  |  |  |  |  |  |  |
| A*02:01 | 0.03 | 0.07 | 0.02 | 0.14 | 0.05 | **6E-04** | 0.05 | **4E-04** | 0.02 | 0.04 | 0.02 | 0.07 |
| **<u>B\*51:01</u>** | 0.04 | 0.06 | 0.03 | 0.28 | 0.10 | **3E-05** | 0.08 | 1E-03 | 0.07 | **2E-05** | 0.05 | 2E-03 |
| B\*39 |  |  |  |  |  |  |  |  |  |  |  |  |
| **<u>B\*39</u>** | -0.17 | **1E-05** | -0.19 | **2E-06** | -0.06 | 0.13 | -0.07 | 0.11 | -0.12 | **2E-05** | -0.13 | **3E-06** |
| DRB1\*12:01 |  |  |  |  |  |  |  |  |  |  |  |  |
| DRB1*12:01 | -0.11 | 0.02 | -0.11 | 0.02 | -0.08 | 0.02 | -0.08 | 0.02 | -0.11 | **1E-04** | -0.11 | **7E-05** |

The allele variants in a given HLA haplotype are presented. Underlined alleles were conferring the strongest effect, determined by stepwise conditional analyses. HLA genotype data was available for 7,063 MS cases and 6,098 controls. <sup>A</sup>Adjustment #1 = Age, sex, 5 PCA vectors, cohort type (incidence or prevalence). When adjusted for HLA, one associated HLA allele from each associated haplotype (underlined in the table) were added to the model in order to test for independent association of the investigated HLA allele. Adjustment #2 = Adjustment #1 + HLA. When all subjects were analyzed together, MS affection status were added as a covariable. Haplotypes associated to IE1A indicated in bold if any allele in it has a p-value <0.001.

Re-analyses were performed adjusting for the most associated HLA allele in each HLA haplotype reported in **Table 3**, and Tables S3 and S4 in Supplementary Material. The number of significant SNPs (p < 5×10^−8^) decreased from 191 to 4 in the IE1A GWAS, and no SNP in the HLA locus remained significantly associated with the antibody responses against IE1B and 101K (Figure S5 and Tables S7 and S8 in Supplementary Material).

Although several HLA haplotypes were found to influence anti-HHV-6A/6B protein specific IgG responses, the association between serological responses and MS disease remained to a large extent unaltered when adjusting for carriage of associated HLA alleles (Tables S3 and S4 in Supplementary Material).

### IgG response to HHV-6A and B proteins interacts with MS risk HLA alleles

We investigated the interaction between IE1A, IE1B and 101K IgG responses and the major MS risk HLA alleles DRB1*15:01 and A*02:01 in conferring risk to MS. High IgG response to IE1A interacted with both DRB1*15:01 and absence of A*02:01 (AP=0.31, p=2×10^−8^ and AP=0.21, p=2×10^−4^ respectively) while IE1B only interacted with DRB1*15:01 (AP=0.19, p=1×10^−3^, **Figure 5**). The interaction between IE1A and DRB1*15:01/A*02:01 on MS risk was only observed in persons with high EBV levels, while the IE1B-DRB1*15:01 interaction was only significant in persons with low EBV levels (data not shown).

**FIGURE 5.**
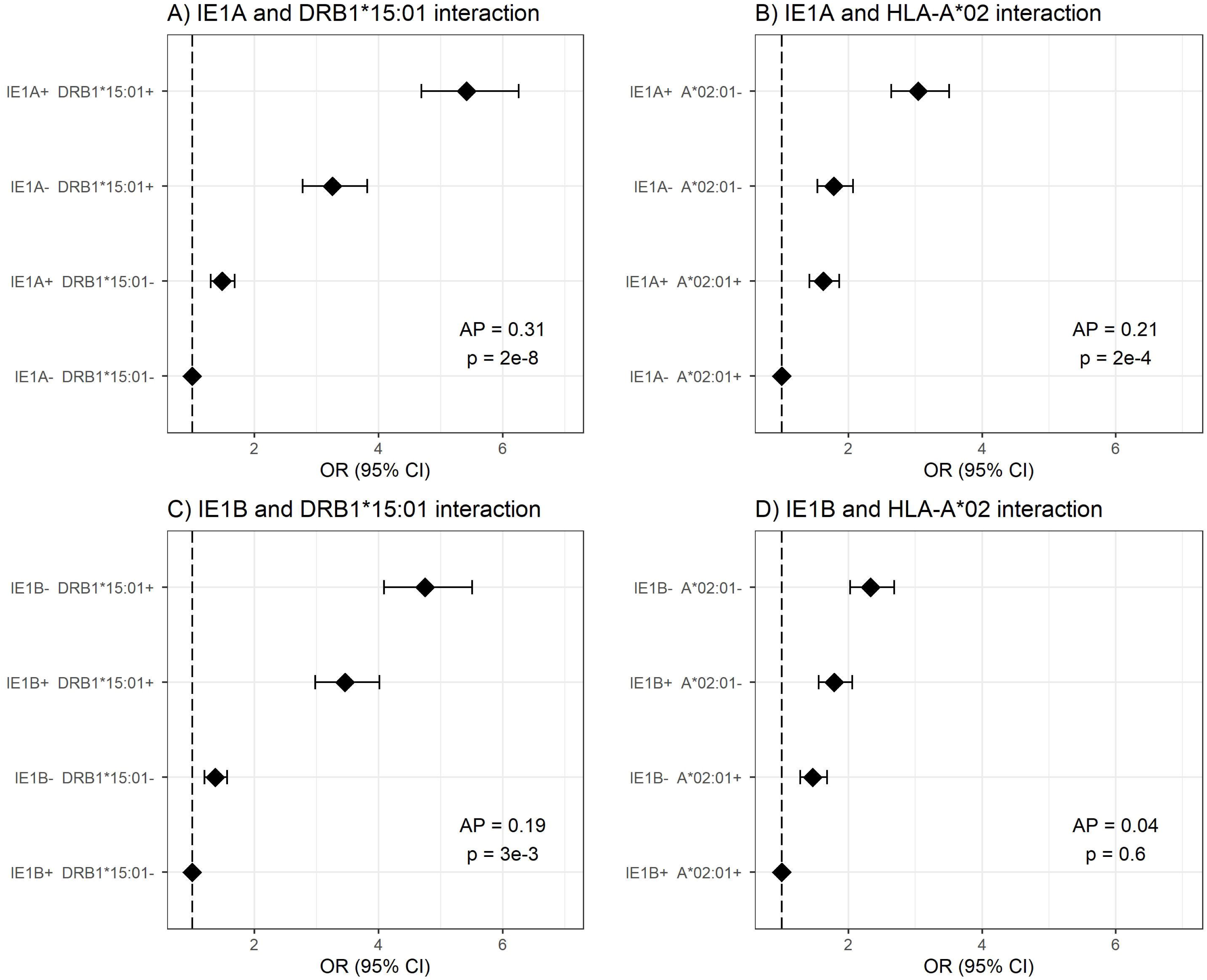
Interaction analysis between IE1A and IE1B IgG response and main MS risk HLA alleles. Odds ratios (OR) and confidence intervals (CI) for A) IE1A and DRB1*15:01, B) IE1A and A*02:01, C) IE1B and DRB1*15:01, and D) IE1B and A*02 were obtained through logistic regression models adjusted for age, sex and cohort type analyzing the Established MS cohort (n=7,063 MS cases and n=6,098 controls). OR were calculated in relation to the group with the lowest MS risk. Plus (+) indicates being a strong responder while minus (-) indicates being a weak responder defined as having an MFI value in the upper quartile of measured response. AP=attributable proportion due to interaction, p is p-value for interaction. No adjustment for EBV was done in these figures.

## DISCUSSION

We show in a large national case-control cohort that persons with MS have higher IgG reactivity against the IE1 protein sequence from HHV-6A compared to controls, while an association in the opposite direction was observed for reactivity against the corresponding IE1 protein sequence from HHV-6B. Importantly, the positive association for IE1A was observed also in samples drawn before MS onset, indicating that it is not simply the state of chronic inflammatory disease that induces the higher level of anti-viral antibodies, but that differences in serological status precede clinical symptom onset. Since having a strong IE1A response during adolescence (< 20 years) conferred the highest risk of developing MS on average 10 years later in life (**Table 2**), our data argues against a reversed causation. We presume that the increased humoral immune reactivity to the selected viral proteins reflects a more intense primary infection and/or reactivation, resulting in higher viral load and therefore an increased anti-viral response. Assuming that HHV-6A infection does play a role in disease onset, this data suggests that acquisition of HHV-6A at a younger age might play an important role in triggering MS.

The positive association seen for HHV-6A (IE1A), but not for HHV-6B (IE1B), with MS disease is in line with some previous studies (8, 10–12) and is interesting considering the differences between the viruses. Both HHV-6A and 6B have the ability to remain latent in the brain (45), but only HHV-6A has been shown to infect and form latent infection in oligodendrocytes (46), the myelin-producing cell and the presumed target of the autoimmune reaction in MS. Speculatively, reactivation of the virus from oligodendrocytes could direct the immune system towards these target cells, suggesting a mechanism that would explain a selective association of HHV-6A with MS. Furthermore, human oligodendrocyte progenitor cells expressing the HHV-6A latency-associated viral protein U94A do not migrate accurately (47), which may yield insufficient myelin repair in the brain and hence could provide another potential link between HHV-6A infection and MS disease. An additional possibility for a potential causative role for HHV-6A in myelin tissue destruction is supported by *in vitro* data showing that supernatants from HHV-6A, but not from HHV-6B, infected cell cultures induce caspase-independent cell death (e.g. necroptosis) in oligodendrocytes (48). This virus-specific pattern is in line with the data in the present study where increased IE1A, but not IE1B, IgG levels are seen in MS plasma. Thus, an association to HHV-6A gives plausible explanations for both the myelin degradation and impaired re-myelination in MS.

Regarding interactions with already established risk factors for MS, we can report an additive interaction between strong IgG responses to IE1A and EBV (**Figure 2A**), not seen for IE1B (Figure S1A in Supplementary Material). This suggest that increased immune response to both viruses are involved in MS and would be consistent with studies reporting that HHV-6A, but not HHV-6B, infection of cells carrying the EBV genome can activate EBV replication (49–51). As HHV-6A infection can activate LMP-1 and EBV nuclear antigen (EBNA)-2 protein expression (50), two proteins important for EBV immortalization of B cells, one can speculate that the increased IE1A IgG levels seen in the present study may be a result of increased infection and transformation of EBV infected B cells. However, that would lead to a general increase of antibody of all specificities including the response against IE1B, which we did not find. An interaction between HHV-6A and EBV in MS has been suggested elsewhere (52), where the author hypothesizes that HHV-6A activates latent EBV in B-cells resident in MS lesions and that both viruses, together, are fundamental for the etio-pathogenic processes of MS. We could in addition observe interaction between the main MS HLA risk alleles and IE1A IgG response (**Figure 5**) indicating that these MS risk factors are acting jointly in increasing risk of MS, at least in a group of patients. Similar interaction with HLA has previously been reported for immune response to EBV (19).

The specificity of the serological response we report here is interesting. As antigens, IE1A proteins are located in the cell nucleus and two relevant questions to consider are how the B-cell activates a response against these antigens and what functions these antibodies might have. Increased IgG responses against intra-nuclear herpesvirus proteins in MS have been reported previously for the HHV-6A and 6B protein p41, and the EBV protein EBNA-1 (53, 54). For B cells to be directed against nuclear or intracellular antigens the cell needs to be disrupted, for example through necroptosis, a form of immunogenic programmed cell death where cell swelling results in rupture of the cell membrane and release of intracellular components into the surrounding tissue (reviewed in (55, 56)). In line with this hypothesis, necroptosis markers have been observed in MS lesions (57) and the intrathecally produced antibodies characteristic of persons with MS often are directed against ubiquitous intracellular proteins (58). The role of these anti-nuclear antigen B-cell responses are less clear. Since the antibodies directed against nuclear antigens probably do not have any neutralizing effect on the viruses, they would not protect against infection, reactivation or dissemination. An anti-IEA response might be seen as a marker of both increased infection and increased tissue destruction and their function might be to clear cell debris. The relevance of these antibodies, protective or detrimental, in infections and regarding the association with autoimmune disease is yet to be determined.

Due to the high similarity and potential antibody cross-reactivity between HHV-6A and HHV-6B, methods discriminating between their serological responses have been difficult to develop. The IE1A and IE1B sequences used in our assay align to some extent (Figure S9 and Table S6 in Supplementary Material) and the possibility of cross-reactivity should not be neglected. However, the lack of correlation between the IE1A and IE1B serological measurements (Figure S6 in Supplementary Material) in combination with their associations with MS in opposite directions (**Table 1** and **2**), suggests that the method indeed has the potential to discriminate between HHV-6A and HHV-6B. Validating the method using serum from children with primary infection further supported this notion, where seroconversion after primary HHV-6B infection was seen for IE1B and 101K only (Table S5 in Supplementary Material).

Serological responses against all three antigens investigated in this study were mainly influenced by genetic factors in the HLA region. HLA associations with serological responses have been seen before (59) and is expected, since long lasting and IgG isotype switched B-cell response is T-cell dependent and facilitated through interaction between HLA and the T-cell receptor. The associated HLA haplotypes were relatively similar in MS and controls, suggesting that the influence of HLA had more to do with control of the viral infection than MS disease. Overall, the known MS-associated HLA haplotype DRB1*15:01 (60) was not associated with serological levels and the associations to MS were still significant after correction for the major MS associated HLA alleles, which would indicate that the serological response could not be explained solely by previous known genetic risk factors for MS. An interesting exception was the association of IE1A and presence of HLA-A*02:01 that was seen in controls, but not for persons with MS. Moreover, we found an interaction between both the DRB1*15:01 and absence of HLA-A*02 (the extended HLA haplotype confirming the highest risk for MS) with IE1A only in persons with high EBV levels, and an interaction of DRB1*15:01 with IE1B only in persons with low EBV levels. Thus, the interaction between HHV-6A and MS associated HLA alleles seems to have effect on the risk for MS only in persons with high anti-EBV response.

The IE1A antigen had the strongest HLA association in comparison with the other antigens investigated, but this might only reflect the ability of the HLA systems to respond differently to different infections on a population level. Furthermore, individuals respond differently to antigens from the same virus (Figure S6B in Supplementary Material). Difference in protein structure, location, phase of expression, and function for the IE1 or 101K proteins (37) possibly makes the immune system encounter them under divergent conditions.

In the present study we could confirm some of our previously data using the HHV-6B lysate based commercial ELISA, namely lower serological response against HHV-6B lysate (32) and the 101K protein in males and in HLA-A*02 carriers. Female sex has been associated with increased acquisition of HHV-6B in children (5), and it is possible that the HHV-6B lysate IgG (32) and 101K IgG responses reflect this difference in primary HHV-6B infection. The association with HLA-A*02, found in our previous study (32) and suggestively confirmed for anti-101K IgG response in the present study, indicates a role for CD8+ T cells in the immune response against HHV-6B. In line with this notion, 101K peptides have been shown to be presented by HLA-A*02:01 on HHV-6B infected cells and these cells are recognized and killed by CD8+ T cells (59). One can hypothesize that if infected cells are removed, the systemic viral burden may decrease, thus possibly explaining the lower levels of anti-HHV-6A/6B (32) and anti-101K IgG levels in individuals with the HLA-A*02 allele.

In conclusion, we provide strong serological data supporting a role for HHV-6A in MS etiology, though causality, as with all forms of association studies remain to be proven.

## MATERIALS AND METHODS

### Study subjects

Two different patient cohorts were used, one with samples taken during MS disease (Established MS, **Table 4**) and one where samples had been collected before MS onset (pre-MS, **Table 5**). Some individuals (348 of the RRMS patients and 1 control) included in the Established MS cohort were also included in the pre-MS cohort. These individuals were excluded from the Established MS cohort when association between serological response and MS disease was analyzed in both cohorts, but were included in other analyses.

**Table 4.** Demographic data of the MS cohort.

|  | MS cases | Controls | All subjects |
| --- | --- | --- | --- |
| Number of subjects | 8742 | 7215 | 15957 |
| % Females | 72% | 75% | 74% |
| Age at sampling [mean ± SD] | 47.1 ± 14.0 | 48.2 ± 13.4 | 47.6 ± 13.8 |
| Age category |  |  |  |
| <20 years | 112 [1%] | 53 [1%] | 165 [1%] |
| 20-29 years | 972 [11%] | 611 [8%] | 1583 [10%] |
| 30-39 years | 1831 [21%] | 1471 [20%] | 3302 [21%] |
| 40-49 years | 2115 [24%] | 1832 [25%] | 3948 [25%] |
| 50-59 years | 1918 [22%] | 1707 [24%] | 3625 [23%] |
| >60 years | 1794 [21%] | 1541 [21%] | 3335 [21%] |
| Median (IQR) age at MS onset<sup>A</sup> | 32 (15) | - | - |
| Median (IQR) years between symptom onset and serum collection<sup>B</sup> | 10.9 (17.3) | - | - |
| Disease course at sampling |  |  |  |
| RRMS | 5586 [64%] | - | - |
| SPMS | 1804 [21%] | - | - |
| PPMS | 549 [6%] | - | - |
| Missing data / Other | 803 [9%] | - | - |
| Ever smokers<sup>C</sup> | 3336 [51%] | 2461 [41%] | 5797 [46%] |

<sup>A</sup>Median age when the first MS symptom was reported to have occurred, i.e. not the same as age at MS diagnosis, data available for 8,505 persons with MS.
<sup>B</sup>Median disease duration. in years, calculated as the time from age at onset to age at sampling.
<sup>C</sup>Past and/or present regular smoking habits, smoking data was obtained for 12,530 individuals.

**Table 5.** Demographic data of the Pre-MS cohort.

|  | **MS Cases** | **Controls** | **All Subjects** |
| --- | --- | --- | --- |
| Number of subjects | 478 | 476 | 954 |
| % Female | 83% | 83% | 83% |
| Age at sampling [mean ± SD] | 24.7 ± 6.4 | 24.6 ± 6.4 | 24.7 ± 6.4 |
| Age category |  |  |  |
| < 20 years | 93 [20%] | 91 [19%] | 184 [19%] |
| 20-29 years | 274 [57%] | 279 [59%] | 553 [58%] |
| 30-39 years | 111 [23%] | 106 [22%] | 217 [23%] |
| Median (IQR) age at MS onset | 34 (12) | - | - |
| Median (IQR) years between symptom onset and serum collection | - 8.3 (10) | - | - |
| Disease course at MS onset |  |  |  |
| RRMS | 478 [100%] | - | - |

### Established MS cohort

The established MS cohort included persons with MS and matched controls from the Epidemiological Investigation of Multiple Sclerosis (EIMS (44), n = 5674), Genes and Environment in Multiple Sclerosis (GEMS (61), n = 8903), Immunomodulation and Multiple Sclerosis Epidemiology study (IMSE (62), n = 1079) and Stockholm Prospective Assessment of Multiple Sclerosis (SPASM / STOPMS (63), n = 301). EIMS and STOPMS have an incidence design, i.e. newly diagnosed persons with MS were invited to join these studies, whereas GEMS and IMSE have a prevalence design, i.e. already diagnosed persons with MS were invited to join these studies. To each person with MS included in GEMS and EIMS, several non-MS individuals, randomly selected from the National population register, matched for age at diagnosis, sex and residency, were invited to participate as control subject. In total, 8742 persons with MS and 7215 controls were included (**Table 4**). Blood samples were either drawn and shipped over night in room temperature prior to plasma isolation or frozen within a few hours after sampling. Plasma samples were stored at –80 ^°^C until analysis. Information regarding smoking habits were obtained for 6534 persons with MS and 5996 controls through self-reported questionnaires (64). Data regarding disease characteristics was obtained mainly through the national Swedish MS registry (65) and in some instances from medical records. All MS cases fulfilled the McDonald Criteria (66, 67) and all study participants provided written, informed consent. Some (n=497) individuals had given more than one sample and for these only the earliest collected sample was included. This study was conducted in line with the aims of the EIMS, IMSE, GEMS and SPASM/STOPMS studies, all which were approved by the Regional Ethical Review Board in Stockholm and performed according to the ethical standards of the Declaration of Helsinki.

Disease severity, Multiple Sclerosis Severity Scores (MSSS) (68) and Age Related Multiple Sclerosis Severity scores (ARMSS) (69), was calculated from expanded disability status scale (EDSS) measurements reported to the Swedish MS registry by the treating neurologist.

### Pre-MS cohort

The pre-MS cohort is a prospective case-control study on biobank samples drawn before symptom onset (**Table 5**, Figure S8 in Supplementary Material). As of February 2012, the Swedish MS registry (65) containing 11,196 MS cases was cross-linked with three Swedish microbiological biobanks which contain the remainders of sera after clinical microbiological analyses performed at the University Hospitals of Skåne and Gothenburg, and the Public Health Agency of Sweden. The serum samples had been stored at –20 ^°^C in these biobanks until analysis.

From individuals who later developed RRMS, only samples drawn before the age of 40 were included (n=478). Individuals who did not develop MS served as controls (n=476). Controls were matched for biobank, sex, date of blood sampling and date of birth, in order of decreasing priority. The un-even number of cases and controls was due to dropout of study participants after the matching process, leaving 474 matched sets of cases and controls. Between cases and controls the mean difference in age at sampling and date of serum collection was 65 days and 6 days, respectively. For six individuals there were no matching case or control but these individuals were still included in the analyses. In cases where registry data were incomplete, notably time of onset, the registered data were checked and corrected at one of the contributing center. The study was approved by the Regional Ethical Review Board in Umeå and performed according to the ethical standards of the Declaration of Helsinki.

### Measurement of IgG antibodies

For detection of IgG antibodies against the different viral proteins, a multiplex serological assay using beads coated with recombinant glutathione s-transferase (GST) fusions proteins was used. The assay procedure has been described in detail elsewhere (70). In short, antigens were expressed as GST fusion proteins using modified pGEX vectors in *E. coli*. Four different HHV-6 protein sequences were expressed: HHV-6A and -6B specific regions of the IE1 protein, IE1A and IE1B respectively, and a divergent region of the structural protein 101K (HHV-6B) and p100 (HHV-6A) (Figure S9 and Table S6 in Supplementary Material). The antigen expressing bacteria were lysed, the lysate cleared of insoluble components and thereafter *in situ* purified on specific polystyrene beads set (SeroMap, Luminex Corporation) coupled to glutathione-casein (GC). Plasma/serum from the study subjects were diluted 1:1000 and pre-incubated with GST lysate (71) to remove antibodies specific for GST or wildtype bacterial proteins present in the lysate. Beads coated with different antigens were mixed and incubated with pre-incubated plasma/serum in filter bottom 96-well plates. After washing, a biotinylated goat-anti human IgG secondary antibody (Dianova) was added and, after additional washing steps, detected by streptavidin-R-phycoerythrin (Moss). Median fluorescence intensity (MFI) was measured with a Luminex 200 analyzer. On every plate, four plate controls were tested to assess assay variation, as described in detail elsewhere (72, 73), yielding antigen-specific coefficients of variation of 13.2 % – 18.7 % for the EBV and CMV antigens. The plate controls did not react with any of the HHV-6A or B antigens.

In addition to the HHV-6 antigens, antibody responses against other viral proteins were measured with this multiplex assay (as described above)(74). The IgG responses against four CMV antigens (pp150, pp52, pp28, pp65) and two EBV antigens (one EBNA1 peptide sequence, aa 385-420 (75), and VCA p18) were used to calculate CMV and EBV indexes. These indexes reflect an overall response against several epitopes of the viruses on a continuous scale. The indexes for one individual were calculated as the sum of the fractions of MFI determined by the measured virus protein over the median MFI among controls of that specific protein. In addition, three HHV-6A proteins (major capsid protein (MCP), U94A and p100) were first included as antigens but were excluded due to low reactivity.

### Batch control

The samples were run in different batches over different days. To compensate for the batch-to-batch variation, inter-batch controls (two plates with 180 samples) were analyzed within each run and used to correct for the variations. Standard linear model, or non-linear models (e.g. logarithmic, exponential) where appropriate, were used to adjust the batch variation and only batch-corrected MFI values are presented in this study.

### Validation of assay

To investigate the specificity of the Luminex assay, samples from 10 children with *exanthema subitum* (ES; HHV-6B primary infection) were investigated. For all individuals, one sample was available from the acute ES phase and one from the convalescence phase. The median time between these samples was 7.5 days. If considering 7 days as the minimum time to mount an antibody response, only 5 individuals were retained in the analysis as individuals that should be HHV-6B positive. Comparing the samples collected during the acute ES phase with the samples collected during the convalescence phase of these five HHV-6B infected children, a ≥10 times increase in MFI value was observed for 4 individuals (80%) for the 101K antigen, and for 2 individuals (40%) for IE1B. The antibody responses against the two HHV-6A protein sequences (IE1A and p100) were not increased in the samples from the convalescence phase. Although based on few individuals, these results suggest that the assay can detect seroconversions upon primary HHV-6B infection and that negligible cross-reactivity occurs between 101K and p100, or between IE1A and IE1B.

Measured reactivities against IE1A, IE1B, 101K and p100 for the ES samples are presented in Table S3 in Supplementary Material. The ES control samples were analyzed separately from the MS study samples using a secondary antibody directed against IgA+IgM+IgG (Dianova) instead of the anti-IgG antibody described above, and the results from 1:100 dilutions were used instead of 1:1000 as for the MS samples. Thus, a direct comparison of the MFI values between the MS and ES samples is not possible, instead the relative MFI shifts between the paired samples are more informative.

To validate if the differential seroresponse between acute and convalescent ES (n=5) constitute a true seroconversion, the difference in measures were compared to the general variability of antibody response from 39 paired (established MS) samples collected < 2 months apart, but with a minimum time of 7 days between the two samples. The response against 101K was significantly elevated in the convalescent sample from the ES cohort compared to the second sample of the adult MS cohort (p=0.0003, Mann Whitney U test). The relative changes of the other antigens were not significantly different.

Together, these results indicate that the assay detect HHV-6B IgG responses correctly, but whether an HHV-6A infection elicit only IE1A and/or p100 IgG responses could not be determined.

### Statistics

Statistical analyses were performed in the software R, version 3.4, on data from the established MS cohort, and with SPSS, version 23, on data from the pre-MS cohort. All graphs were constructed in R version 3.4.

To control for potential confounders, the antibody responses were analyzed using regression models adjusted for age and sex. The established MS cohort was also adjusted for study design (incidence or prevalence design) in all regression analyses. As the p100 IgG responses were mainly in the technical noise area of the assay, the anti-p100 response was therefore excluded from the analyses.

Associations between MS and antibody status were investigated using logistic regression models comparing the MS frequency in strong and weak responders. Individuals with MFI values in the 4^th^ quartile among controls were regarded as strong responders and individuals with MFI values in 1^st^ quartile among controls were regarded as weak responders. To get a clear separation between strong and weak responders, the in-between responders (individuals with MFI values in between 25^th^ and 75^th^ percentile) were excluded in this analysis. Strong and weak responders may reflect seropositive and seronegative individuals, but true serostatus cannot be determined due to lack of validated control samples. The quartiles were determined separately for the two cohorts and are indicated in **Figure 1**.

To assess if additive interaction occurred between antibody responses to different viruses and between MS HLA risk alleles and HHV-6 immune response in and MS risk, and between MS HLA risk alleles and antibody response, interaction analyses were performed in the established MS cohort. This was done by calculating the proportion attributable to interaction (AP) using logistic regression analysis, where the odds ratios were calculated in relation to the group with the lowest MS risk (76).

Antibody levels were analyzed using linear regression models. The IgG responses were heavily left-skewed. To obtain a more normal distribution, the antibody levels (MFI values) were transformed using a log base 10 transformation prior to statistical analysis. When investigating how MS disease influenced the antibody levels, the models were conducted with unadjusted or adjusted data for potential confounders in order to see if the association between MS and antibody levels remained (Tables S1 and S2 in Supplementary Material). In this analysis, both study cohorts were adjusted for EBV and CMV index and the established MS cohort was also adjusted for carriage of HLA alleles (MS risk alleles HLA-A*02:01, HLA-DRB1*15:01 and the alleles associated with each antibody response indicated in **Table 3** and Table S3 and S4 in Supplementary Material).

Correlations between antibody measurements were investigated with Spearman correlation tests (Figures S5 and S6 in Supplementary Material).

The main hypothesis tested in this study is that HHV-6A/6B IgG responses are associated with MS disease. To test this hypothesis, three antibody specificities were measured in two different study cohorts, corresponding to 6 independent tests thus using the Bonferroni correction on an alpha level of 0.05 yields a threshold of 0.008 for significance. When other hypotheses were answered, the same alpha value of 0.008 was regarded as significant.

### GWAS

Genotypes for ∼720,000 single nucleotide polymorphisms (SNPs) were determined using an Illumina OmniExpress BeadChip for 6,396 MS cases and 5,530 controls from the established MS cohort. SNPs with a minor allele frequency of < 2%, with a call-rate of less than 98% or those which were not in Hardy-Weinberg equilibrium among controls (p < 0.0001) were removed from analysis. Individuals with > 2% failed genotype calls, with increased heterozygosity (> mean + 2 SD), related individuals (increased identity by descent, IBD), or individuals where the recorded sex differed from the genotype result were removed from analysis. Population outliers identified using the SmartPCA program were removed. A principal component analysis was conducted using Eigensoft (77) and five PCA components were used to control for population stratification. Linear regression models investigating the association between SNPs and Log10-transformed IE1A, IE1B and 101K antibody levels were analyzed both separately for MS cases and controls, and for all subjects together, using PLINK v1.9 (78) adjusted for 5 PCA vectors, age, sex, cohort type (and MS disease when all subjects were analyzed together). Logistic regression models, adjusted as the linear regression models, were performed to investigate the association between SNPs and strong or weak response (comparing the 3^rd^ and 1^st^ quartile respectively) against IE1A, IE1B, and 101K.

Associations for SNPs with p < 5×10^−8^ were regarded as significant, but all SNPs with suggestive association (p < 10^−5^) are reported in Tables S4 and S5 in Supplementary Material. Genome build GRCh37 was used.

### HLA-imputation and Associations with HLA haplotypes

HLA allele variants for MHC class I and II were imputed by the software HLA*IMP:02 (79) for 7,063 MS cases and 6,098 controls using genotypes from the MS Replication Chip (80). This chip densely covered in the MHC region. Associations between HLA alleles and IgG responses against each HHV-6 antigen were determined by linear regression models using R version 3.3.1. Associated alleles were combined into haplotypes using previously reported common haplotypes in the Caucasian population (81). Analyses were stratified by MS affection status, but were also conducted on persons with MS patients together with controls. Models were adjusted for age, sex, cohort type (incidence or prevalence), 6 PCA vectors, and for MS disease status when all subjects were analyzed together. To determine the allele with the strongest effect in each associated haplotype, stepwise conditional analyses were performed. Secondary analyses were made where associated HLA alleles were added as co-variables to test for independent associations.

## ACKNOWLEDGEMENTS

The authors thank Dr. Steven Jacobson, Dr. Elisabetta Caselli and Dr. Dario Di Luca for providing serum for method validation. The authors also thank Ingileif Jonsdottir and Kári Stefánsson for genotyping done by deCODE, and Alexander T Dilthey at Wellcome Trust Centre for Human Genetics, University of Oxford, for contribution in HLA imputation. Swedish Research Council (grant no 2015-02419), Swedish Brain Foundation, KAW Foundation, Margareta af Ugglas Foundation. J Huang, P Stridh and I Kockum were supported partly by funding from MultipleMS Horizon 2020 grant number 733161. J Huang was also partly supported by EndMS doctoral studentship EGID: 3045 from the Multiple Sclerosis society of Canada.

## AUTHOR CONTRIBUTIONS STATEMENT

EE contributed to design and interpretation of the work, acquisition, statistical analyses, design of tables and figures for the establish MS cohort. RG identified the antigens used. JH contributed to genetic analyses and choice of statistical methods. MB was responsible for the pre-MS cohort.

ILB, PS, MK were responsible for the MS case control cohort database and biobank. ILB did HLA imputation and selection of viral antigens. PS managed the GWAs genotyping. AKH was responsible for the EIMS study and HLA interactions.

NB, JB and AM set up and validated the multiplex serological assay. DJ, MH and LAM contributed to the pre-MS cohort.

LF helpful in designed the protein antigens.

MI and TY were contributing to the validation. OA contributed to the pre-MS and MS cohort. JAH and LA initiated of register and biobank.

TW was responsible for the multiplex serological assay.

PS initiated the pre-MS cohort and supervised interpretation.

TO initiated the serological analyses, initiated and governed the MS case control register, genetic analyses and biobank.

IK initiated the serological analyses and governed the MS case control register, genetic analyses and biobank.

AFH supervised the initiation, execution and finalization of the HHV-6A/B serology study.

## CONFLICT OF INTEREST STATEMENT

Jan Hillert has received honoraria for serving on advisory boards for Biogen, Sanofi-Genzyme and Novartis and speaker’s fees from Biogen, Novartis, Merck-Serono, Bayer-Schering, Teva and Sanofi-Genzyme. He has served as P.I. for projects, or received unrestricted research support from, BiogenIdec, Merck, Novartis and Sanofi-Genzyme. Lucia Alonso Magdalena has received speaking honoraria from Merck-Serono and served at the advisory board for Merck-Serono and Biogen. Lars Alfredsson received speaker honoraria from Biogen Idec and TEVA. Tomas Olsson has received lecture and/or Advisory board honoraria, and unrestricted MS research grants from Biogen, Novartis, Sanofi, Merck-Serono and Roche. Anna Fogdell-Hahn has received speaker’s fees from Pfizer, Biogen, Merck-Serono, and Sanofi-Genzyme. She has served as P.I. for projects, or received unrestricted research support from, Biogen Idec and Pfizer. All the other co-authors declare no conflict of interest.

## REFERENCES

1. Ablashi D, Agut H, Alvarez-Lafuente R, Clark DA, Dewhurst S, DiLuca D, et al. Classification of HHV-6A and HHV-6B as distinct viruses. Arch Virol (2014) 159(5):863–70. doi: 10.1007/s00705-013-1902-5. PubMed PMID: 24193951; PubMed Central PMCID: PMCPMC4750402.

2. Hall CB, Caserta MT, Schnabel KC, Long C, Epstein LG, Insel RA, et al. Persistence of human herpesvirus 6 according to site and variant: possible greater neurotropism of variant A. Clin Infect Dis (1998) 26(1):132–7. PubMed PMID: 9455521.

3. Dewhurst S, McIntyre K, Schnabel K, Hall CB. Human herpesvirus 6 (HHV-6) variant B accounts for the majority of symptomatic primary HHV-6 infections in a population of U.S. infants. J Clin Microbiol (1993) 31(2):416–8. PubMed PMID: 8381815; PubMed Central PMCID: PMCPMC262777.

4. Hall CB, Long CE, Schnabel KC, Caserta MT, McIntyre KM, Costanzo MA, et al. Human herpesvirus-6 infection in children. A prospective study of complications and reactivation. N Engl J Med (1994) 331(7):432–8. doi: 10.1056/NEJM199408183310703. PubMed PMID: 8035839.

5. Zerr DM, Meier AS, Selke SS, Frenkel LM, Huang ML, Wald A, et al. A population-based study of primary human herpesvirus 6 infection. N Engl J Med (2005) 352(8):768–76. doi: 10.1056/NEJMoa042207. PubMed PMID: 15728809.

6. Hill JA, Zerr DM. Roseoloviruses in transplant recipients: clinical consequences and prospects for treatment and prevention trials. Curr Opin Virol (2014) 9:53–60. doi: 10.1016/j.coviro.2014.09.006. PubMed PMID: 25285614; PubMed Central PMCID: PMCPMC4570620.

7. Akhyani N, Berti R, Brennan MB, Soldan SS, Eaton JM, McFarland HF, et al. Tissue distribution and variant characterization of human herpesvirus (HHV)-6: increased prevalence of HHV-6A in patients with multiple sclerosis. J Infect Dis (2000) 182(5):1321–5. doi: 10.1086/315893. PubMed PMID: 11023456.

8. Alvarez-Lafuente R, De las Heras V, Bartolome M, Picazo JJ, Arroyo R. Relapsing-remitting multiple sclerosis and human herpesvirus 6 active infection. Arch Neurol (2004) 61(10):1523–7. doi: 10.1001/archneur.61.10.1523. PubMed PMID: 15477505.

9. Alvarez-Lafuente R, Garcia-Montojo M, De las Heras V, Bartolome M, Arroyo R. Clinical parameters and HHV-6 active replication in relapsing-remitting multiple sclerosis patients. J Clin Virol (2006) 37 Suppl 1:S24–6. doi: 10.1016/S1386-6532(06)70007-5. PubMed PMID: 17276363.

10. Rotola A, Merlotti I, Caniatti L, Caselli E, Granieri E, Tola MR, et al. Human herpesvirus 6 infects the central nervous system of multiple sclerosis patients in the early stages of the disease. Mult Scler (2004) 10(4):348–54. PubMed PMID: 15327028.

11. Soldan SS, Leist TP, Juhng KN, McFarland HF, Jacobson S. Increased lymphoproliferative response to human herpesvirus type 6A variant in multiple sclerosis patients. Ann Neurol (2000) 47(3):306–13. PubMed PMID: 10716249.

12. Virtanen JO, Farkkila M, Multanen J, Uotila L, Jaaskelainen AJ, Vaheri A, et al. Evidence for human herpesvirus 6 variant A antibodies in multiple sclerosis: diagnostic and therapeutic implications. J Neurovirol (2007) 13(4):347–52. doi: 10.1080/13550280701381332. PubMed PMID: 17849318.

13. International Multiple Sclerosis Genetics C, Wellcome Trust Case Control C, Sawcer S, Hellenthal G, Pirinen M, Spencer CC, et al. Genetic risk and a primary role for cell-mediated immune mechanisms in multiple sclerosis. Nature (2011) 476(7359):214–9. Epub 2011/08/13. doi: 10.1038/nature10251. PubMed PMID: 21833088; PubMed Central PMCID: PMC3182531.

14. International Multiple Sclerosis Genetics C, Beecham AH, Patsopoulos NA, Xifara DK, Davis MF, Kemppinen A, et al. Analysis of immune-related loci identifies 48 new susceptibility variants for multiple sclerosis. Nat Genet (2013) 45(11):1353–60. doi: 10.1038/ng.2770. PubMed PMID: 24076602; PubMed Central PMCID: PMCPMC3832895.

15. Olsson T, Barcellos LF, Alfredsson L. Interactions between genetic, lifestyle and environmental risk factors for multiple sclerosis. Nat Rev Neurol (2017) 13(1):25–36. doi: 10.1038/nrneurol.2016.187. PubMed PMID: 27934854.

16. Angelini DF, Serafini B, Piras E, Severa M, Coccia EM, Rosicarelli B, et al. Increased CD8+ T cell response to Epstein-Barr virus lytic antigens in the active phase of multiple sclerosis. PLoS Pathog (2013) 9(4):e1003220. doi: 10.1371/journal.ppat.1003220. PubMed PMID: 23592979; PubMed Central PMCID: PMCPMC3623710.

17. Bjornevik K, Riise T, Bostrom I, Casetta I, Cortese M, Granieri E, et al. Negative interaction between smoking and EBV in the risk of multiple sclerosis: The EnvIMS study. Mult Scler (2017) 23(7):1018–24. doi: 10.1177/1352458516671028. PubMed PMID: 27663872.

18. Serafini B, Rosicarelli B, Franciotta D, Magliozzi R, Reynolds R, Cinque P, et al. Dysregulated Epstein-Barr virus infection in the multiple sclerosis brain. J Exp Med (2007) 204(12):2899–912. doi: 10.1084/jem.20071030. PubMed PMID: 17984305; PubMed Central PMCID: PMCPMC2118531.

19. Sundqvist E, Sundstrom P, Linden M, Hedstrom AK, Aloisi F, Hillert J, et al. Epstein-Barr virus and multiple sclerosis: interaction with HLA. Genes Immun (2012) 13(1):14–20. doi: 10.1038/gene.2011.42. PubMed PMID: 21776012.

20. Sundstrom P, Juto P, Wadell G, Hallmans G, Svenningsson A, Nystrom L, et al. An altered immune response to Epstein-Barr virus in multiple sclerosis: a prospective study. Neurology (2004) 62(12):2277–82. PubMed PMID: 15210894.

21. Ascherio A, Munger KL, Lunemann JD. The initiation and prevention of multiple sclerosis. Nat Rev Neurol (2012) 8(11):602–12. doi: 10.1038/nrneurol.2012.198. PubMed PMID: 23045241; PubMed Central PMCID: PMCPMC4467212.

22. Sundqvist E, Bergstrom T, Daialhosein H, Nystrom M, Sundstrom P, Hillert J, et al. Cytomegalovirus seropositivity is negatively associated with multiple sclerosis. Mult Scler (2014) 20(2):165–73. doi: 10.1177/1352458513494489. PubMed PMID: 23999606.

23. Okuno T, Takahashi K, Balachandra K, Shiraki K, Yamanishi K, Takahashi M, et al. Seroepidemiology of human herpesvirus 6 infection in normal children and adults. J Clin Microbiol (1989) 27(4):651–3. PubMed PMID: 2542358; PubMed Central PMCID: PMCPMC267390.

24. Hall CB, Caserta MT, Schnabel KC, McDermott MP, Lofthus GK, Carnahan JA, et al. Characteristics and acquisition of human herpesvirus (HHV) 7 infections in relation to infection with HHV-6. J Infect Dis (2006) 193(8):1063–9. doi: 10.1086/503434. PubMed PMID: 16544246.

25. Ward KN, Gray JJ, Fotheringham MW, Sheldon MJ. IgG antibodies to human herpesvirus-6 in young children: changes in avidity of antibody correlate with time after infection. J Med Virol (1993) 39(2):131–8. PubMed PMID: 8387569.

26. Behzad-Behbahani A, Mikaeili MH, Entezam M, Mojiri A, Pour GY, Arasteh MM, et al. Human herpesvirus-6 viral load and antibody titer in serum samples of patients with multiple sclerosis. J Microbiol Immunol Infect (2011) 44(4):247–51. doi: 10.1016/j.jmii.2010.08.002. PubMed PMID: 21524958.

27. Ortega-Madueno I, Garcia-Montojo M, Dominguez-Mozo MI, Garcia-Martinez A, Arias-Leal AM, Casanova I, et al. Anti-human herpesvirus 6A/B IgG correlates with relapses and progression in multiple sclerosis. PLoS One (2014) 9(8):e104836. doi: 10.1371/journal.pone.0104836. PubMed PMID: 25110949; PubMed Central PMCID: PMCPMC4128748.

28. Sola P, Merelli E, Marasca R, Poggi M, Luppi M, Montorsi M, et al. Human herpesvirus 6 and multiple sclerosis: survey of anti-HHV-6 antibodies by immunofluorescence analysis and of viral sequences by polymerase chain reaction. J Neurol Neurosurg Psychiatry (1993) 56(8):917–9. PubMed PMID: 8394408; PubMed Central PMCID: PMCPMC1015152.

29. Khaki M, Ghazavi A, Ghasami K, Rafiei M, Payani MA, Ghaznavi-Rad E, et al. Evaluation of viral antibodies in Iranian multiple sclerosis patients. Neurosciences (Riyadh) (2011) 16(3):224–8. PubMed PMID: 21677611.

30. Derfuss T, Hohlfeld R, Meinl E. Intrathecal antibody (IgG) production against human herpesvirus type 6 occurs in about 20% of multiple sclerosis patients and might be linked to a polyspecific B-cell response. J Neurol (2005) 252(8):968–71. doi: 10.1007/s00415-005-0794-z. PubMed PMID: 15772735.

31. Enbom M, Wang FZ, Fredrikson S, Martin C, Dahl H, Linde A. Similar humoral and cellular immunological reactivities to human herpesvirus 6 in patients with multiple sclerosis and controls. Clin Diagn Lab Immunol (1999) 6(4):545–9. PubMed PMID: 10391860; PubMed Central PMCID: PMCPMC95725.

32. Engdahl E, Gustafsson R, Ramanujam R, Sundqvist E, Olsson T, Hillert J, et al. HLA-A(*)02, gender and tobacco smoking, but not multiple sclerosis, affects the IgG antibody response against human herpesvirus 6. Hum Immunol (2014) 75(6):524–30. doi: 10.1016/j.humimm.2014.03.001. PubMed PMID: 24662416.

33. Villoslada P, Juste C, Tintore M, Llorenc V, Codina G, Pozo-Rosich P, et al. The immune response against herpesvirus is more prominent in the early stages of MS. Neurology (2003) 60(12):1944–8. Epub 2003/06/25. PubMed PMID: 12821737.

34. Dominguez G, Dambaugh TR, Stamey FR, Dewhurst S, Inoue N, Pellett PE. Human herpesvirus 6B genome sequence: coding content and comparison with human herpesvirus 6A. J Virol (1999) 73(10):8040–52. PubMed PMID: 10482553; PubMed Central PMCID: PMCPMC112820.

35. Isegawa Y, Mukai T, Nakano K, Kagawa M, Chen J, Mori Y, et al. Comparison of the complete DNA sequences of human herpesvirus 6 variants A and B. J Virol (1999) 73(10):8053–63. PubMed PMID: 10482554; PubMed Central PMCID: PMCPMC112821.

36. Stanton R, Wilkinson GW, Fox JD. Analysis of human herpesvirus-6 IE1 sequence variation in clinical samples. J Med Virol (2003) 71(4):578–84. doi: 10.1002/jmv.10508. PubMed PMID: 14556272.

37. Gravel A, Gosselin J, Flamand L. Human Herpesvirus 6 immediate-early 1 protein is a sumoylated nuclear phosphoprotein colocalizing with promyelocytic leukemia protein-associated nuclear bodies. J Biol Chem (2002) 277(22):19679–87. doi: 10.1074/jbc.M200836200. PubMed PMID: 11901159.

38. Martin ME, Nicholas J, Thomson BJ, Newman C, Honess RW. Identification of a transactivating function mapping to the putative immediate-early locus of human herpesvirus 6. J Virol (1991) 65(10):5381–90. PubMed PMID: 1654446; PubMed Central PMCID: PMCPMC249019.

39. Jaworska J, Gravel A, Flamand L. Divergent susceptibilities of human herpesvirus 6 variants to type I interferons. Proc Natl Acad Sci U S A (2010) 107(18):8369–74. doi: 10.1073/pnas.0909951107. PubMed PMID: 20404187; PubMed Central PMCID: PMCPMC2889514.

40. Pellett PE, Sanchez-Martinez D, Dominguez G, Black JB, Anton E, Greenamoyer C, et al. A strongly immunoreactive virion protein of human herpesvirus 6 variant B strain Z29: identification and characterization of the gene and mapping of a variant-specific monoclonal antibody reactive epitope. Virology (1993) 195(2):521–31. doi: 10.1006/viro.1993.1403. PubMed PMID: 7687803.

41. Mahmoud NF, Kawabata A, Tang H, Wakata A, Wang B, Serada S, et al. Human herpesvirus 6 U11 protein is critical for virus infection. Virology (2016) 489:151–7. doi: 10.1016/j.virol.2015.12.011. PubMed PMID: 26761397.

42. Yamamoto M, Black JB, Stewart JA, Lopez C, Pellett PE. Identification of a nucleocapsid protein as a specific serological marker of human herpesvirus 6 infection. J Clin Microbiol (1990) 28(9):1957–62. PubMed PMID: 2172295; PubMed Central PMCID: PMCPMC268086.

43. Manouchehrinia A, Westerlind H, Kingwell E, Zhu F, Carruthers R, Ramanujam R, et al. Age Related Multiple Sclerosis Severity Score: Disability ranked by age. Mult Scler (2017) 23(14):1938–46. doi: 10.1177/1352458517690618. PubMed PMID: 28155580; PubMed Central PMCID: PMCPMC5700773.

44. Hedstrom AK, Baarnhielm M, Olsson T, Alfredsson L. Tobacco smoking, but not Swedish snuff use, increases the risk of multiple sclerosis. Neurology (2009) 73(9):696–701. PubMed PMID: 19720976.

45. Cuomo L, Trivedi P, Cardillo MR, Gagliardi FM, Vecchione A, Caruso R, et al. Human herpesvirus 6 infection in neoplastic and normal brain tissue. J Med Virol (2001) 63(1):45–51. PubMed PMID: 11130886.

46. Ahlqvist J, Fotheringham J, Akhyani N, Yao K, Fogdell-Hahn A, Jacobson S. Differential tropism of human herpesvirus 6 (HHV-6) variants and induction of latency by HHV-6A in oligodendrocytes. J Neurovirol (2005) 11(4):384–94. doi: 10.1080/13550280591002379. PubMed PMID: 16162481.

47. Campbell A, Hogestyn JM, Folts CJ, Lopez B, Proschel C, Mock D, et al. Expression of the Human Herpesvirus 6A Latency-Associated Transcript U94A Disrupts Human Oligodendrocyte Progenitor Migration. Scientific reports (2017) 7(1):3978. doi: 10.1038/s41598-017-04432-y. PubMed PMID: 28638124; PubMed Central PMCID: PMCPMC5479784.

48. Kong H, Baerbig Q, Duncan L, Shepel N, Mayne M. Human herpesvirus type 6 indirectly enhances oligodendrocyte cell death. J Neurovirol (2003) 9(5):539–50. doi: 10.1080/13550280390241241. PubMed PMID: 13129768.

49. Cuomo L, Angeloni A, Zompetta C, Cirone M, Calogero A, Frati L, et al. Human herpesvirus 6 variant A, but not variant B, infects EBV-positive B lymphoid cells, activating the latent EBV genome through a BZLF-1-dependent mechanism. AIDS Res Hum Retroviruses (1995) 11(10):1241–5. doi: 10.1089/aid.1995.11.1241. PubMed PMID: 8573381.

50. Cuomo L, Trivedi P, de Grazia U, Calogero A, D’Onofrio M, Yang W, et al. Upregulation of Epstein-Barr virus-encoded latent membrane protein by human herpesvirus 6 superinfection of EBV-carrying Burkitt lymphoma cells. J Med Virol (1998) 55(3):219–26. PubMed PMID: 9624610.

51. Flamand L, Stefanescu I, Ablashi DV, Menezes J. Activation of the Epstein-Barr virus replicative cycle by human herpesvirus 6. J Virol (1993) 67(11):6768–77. PubMed PMID: 8411380; PubMed Central PMCID: PMCPMC238118.

52. Fierz W. Multiple sclerosis: an example of pathogenic viral interaction? Virol J (2017) 14(1):42. doi: 10.1186/s12985-017-0719-3. PubMed PMID: 28241767; PubMed Central PMCID: PMCPMC5330019.

53. Ablashi DV, Lapps W, Kaplan M, Whitman JE, Richert JR, Pearson GR. Human Herpesvirus-6 (HHV-6) infection in multiple sclerosis: a preliminary report. Mult Scler (1998) 4(6):490–6. doi: 10.1177/135245859800400606. PubMed PMID: 9987758.

54. Santiago O, Gutierrez J, Sorlozano A, de Dios Luna J, Villegas E, Fernandez O. Relation between Epstein-Barr virus and multiple sclerosis: analytic study of scientific production. Eur J Clin Microbiol Infect Dis (2010) 29(7):857–66. doi: 10.1007/s10096-010-0940-0. PubMed PMID: 20428908.

55. Dondelinger Y, Hulpiau P, Saeys Y, Bertrand MJ, Vandenabeele P. An evolutionary perspective on the necroptotic pathway. Trends Cell Biol (2016) 26(10):721–32. doi: 10.1016/j.tcb.2016.06.004. PubMed PMID: 27368376.

56. Kunzelmann K. Ion channels in regulated cell death. Cell Mol Life Sci (2016) 73(11-12):2387–403. doi: 10.1007/s00018-016-2208-z. PubMed PMID: 27091155.

57. Ofengeim D, Ito Y, Najafov A, Zhang Y, Shan B, DeWitt JP, et al. Activation of necroptosis in multiple sclerosis. Cell Rep (2015) 10(11):1836–49. doi: 10.1016/j.celrep.2015.02.051. PubMed PMID: 25801023; PubMed Central PMCID: PMCPMC4494996.

58. Brandle SM, Obermeier B, Senel M, Bruder J, Mentele R, Khademi M, et al. Distinct oligoclonal band antibodies in multiple sclerosis recognize ubiquitous self-proteins. Proc Natl Acad Sci U S A (2016) 113(28):7864–9. doi: 10.1073/pnas.1522730113. PubMed PMID: 27325759; PubMed Central PMCID: PMCPMC4948369.

59. Hammer C, Begemann M, McLaren PJ, Bartha I, Michel A, Klose B, et al. Amino Acid Variation in HLA Class II Proteins Is a Major Determinant of Humoral Response to Common Viruses. Am J Hum Genet (2015) 97(5):738–43. doi: 10.1016/j.ajhg.2015.09.008. PubMed PMID: 26456283; PubMed Central PMCID: PMCPMC4667104.

60. Jersild C, Fog T, Hansen GS, Thomsen M, Svejgaard A, Dupont B. Histocompatibility determinants in multiple sclerosis, with special reference to clinical course. Lancet (1973) 2(7840):1221–5. PubMed PMID: 4128558.

61. Hedstrom AK, Hillert J, Olsson T, Alfredsson L. Smoking and multiple sclerosis susceptibility. Eur J Epidemiol (2013) 28(11):867–74. doi: 10.1007/s10654-013-9853-4. PubMed PMID: 24146047; PubMed Central PMCID: PMCPMC3898140.

62. Holmen C, Piehl F, Hillert J, Fogdell-Hahn A, Lundkvist M, Karlberg E, et al. A Swedish national post-marketing surveillance study of natalizumab treatment in multiple sclerosis. Mult Scler (2011) 17(6):708–19. doi: 10.1177/1352458510394701. PubMed PMID: 21228027.

63. Khademi M, Kockum I, Andersson ML, Iacobaeus E, Brundin L, Sellebjerg F, et al. Cerebrospinal fluid CXCL13 in multiple sclerosis: a suggestive prognostic marker for the disease course. Mult Scler (2011) 17(3):335–43. doi: 10.1177/1352458510389102. PubMed PMID: 21135023.

64. Hedstrom AK, Lima Bomfim I, Barcellos L, Gianfrancesco M, Schaefer C, Kockum I, et al. Interaction between adolescent obesity and HLA risk genes in the etiology of multiple sclerosis. Neurology (2014) 82(10):865–72. doi: 10.1212/WNL.0000000000000203. PubMed PMID: 24500647; PubMed Central PMCID: PMCPMC3959752.

65. Hillert J, Stawiarz L. The Swedish MS registry - clinical support tool and scientific resource. Acta Neurol Scand (2015) 132(199):11–9. doi: 10.1111/ane.12425. PubMed PMID: 26046553; PubMed Central PMCID: PMCPMC4657484.

66. McDonald WI, Compston A, Edan G, Goodkin D, Hartung HP, Lublin FD, et al. Recommended diagnostic criteria for multiple sclerosis: guidelines from the International Panel on the diagnosis of multiple sclerosis. Ann Neurol (2001) 50(1):121–7. PubMed PMID: 11456302.

67. Polman CH, Reingold SC, Edan G, Filippi M, Hartung HP, Kappos L, et al. Diagnostic criteria for multiple sclerosis: 2005 revisions to the “McDonald Criteria”. Ann Neurol (2005) 58(6):840–6. doi: 10.1002/ana.20703. PubMed PMID: 16283615.

68. Roxburgh RH, Seaman SR, Masterman T, Hensiek AE, Sawcer SJ, Vukusic S, et al. Multiple Sclerosis Severity Score: using disability and disease duration to rate disease severity. Neurology (2005) 64(7):1144–51. doi: 10.1212/01.WNL.0000156155.19270.F8. PubMed PMID: 15824338.

69. Manouchehrinia A, Westerlind H, Kingwell E, Zhu F, Carruthers R, Ramanujam R, et al. Age Related Multiple Sclerosis Severity Score: Disability ranked by age. Mult Scler (2017):1352458517690618. doi: 10.1177/1352458517690618. PubMed PMID: 28155580.

70. Waterboer T, Sehr P, Michael KM, Franceschi S, Nieland JD, Joos TO, et al. Multiplex human papillomavirus serology based on in situ-purified glutathione s-transferase fusion proteins. Clin Chem (2005) 51(10):1845–53. doi: 10.1373/clinchem.2005.052381. PubMed PMID: 16099939.

71. Waterboer T, Sehr P, Pawlita M. Suppression of non-specific binding in serological Luminex assays. J Immunol Methods (2006) 309(1-2):200–4. doi: 10.1016/j.jim.2005.11.008. PubMed PMID: 16406059.

72. Kreimer AR, Johansson M, Yanik EL, Katki HA, Check DP, Lang Kuhs KA, et al. Kinetics of the Human Papillomavirus Type 16 E6 Antibody Response Prior to Oropharyngeal Cancer. J Natl Cancer Inst (2017) 109(8). doi: 10.1093/jnci/djx005. PubMed PMID: 28376197.

73. Michael KM, Waterboer T, Sehr P, Rother A, Reidel U, Boeing H, et al. Seroprevalence of 34 human papillomavirus types in the German general population. PLoS Pathog (2008) 4(6):e1000091. doi: 10.1371/journal.ppat.1000091. PubMed PMID: 18566657; PubMed Central PMCID: PMCPMC2408730.

74. Brenner N, Mentzer AJ, Butt J, Michel A, Prager K, Brozy J, et al. Validation of Multiplex Serology detecting human herpesviruses 1-5. PLoS One (2018) 13(12):e0209379. doi: 10.1371/journal.pone.0209379. PubMed PMID: 30589867; PubMed Central PMCID: PMCPMC6307738.

75. Salzer J, Nystrom M, Hallmans G, Stenlund H, Wadell G, Sundstrom P. Epstein-Barr virus antibodies and vitamin D in prospective multiple sclerosis biobank samples. Mult Scler (2013) 19(12):1587–91. doi: 10.1177/1352458513483888. PubMed PMID: 23549431.

76. Andersson T, Alfredsson L, Kallberg H, Zdravkovic S, Ahlbom A. Calculating measures of biological interaction. Eur J Epidemiol (2005) 20(7):575–9. PubMed PMID: 16119429.

77. Price AL, Patterson NJ, Plenge RM, Weinblatt ME, Shadick NA, Reich D. Principal components analysis corrects for stratification in genome-wide association studies. Nat Genet (2006) 38(8):904–9. doi: 10.1038/ng1847. PubMed PMID: 16862161.

78. Chang CC, Chow CC, Tellier LC, Vattikuti S, Purcell SM, Lee JJ. Second-generation PLINK: rising to the challenge of larger and richer datasets. Gigascience (2015) 4:7. doi: 10.1186/s13742-015-0047-8. PubMed PMID: 25722852; PubMed Central PMCID: PMCPMC4342193.

79. Dilthey AT, Moutsianas L, Leslie S, McVean G. HLA*IMP--an integrated framework for imputing classical HLA alleles from SNP genotypes. Bioinformatics (2011) 27(7):968–72. doi: 10.1093/bioinformatics/btr061. PubMed PMID: 21300701; PubMed Central PMCID: PMCPMC3065693.

80. Consortium IMSG. The Multiple Sclerosis Genomic Map: Role of peripheral immune cells and resident microglia in susceptibility(2017.

81. Gonzalez-Galarza FF, Takeshita LY, Santos EJ, Kempson F, Maia MH, da Silva AL, et al. Allele frequency net 2015 update: new features for HLA epitopes, KIR and disease and HLA adverse drug reaction associations. Nucleic Acids Res (2015) 43(Database issue):D784–8. doi: 10.1093/nar/gku1166. PubMed PMID: 25414323; PubMed Central PMCID: PMCPMC4383964.

