## Supplemental figures and tables for "Increased serological response against human herpesvirus 6A is associated with risk for multiple sclerosis"

#### 1 Supplementary Figures and Tables for

**<u>Supplementary Figures and Tables contains:</u>**

**Figure S1.** Interaction of antibody response against different herpesviruses in association to MS

**Figure S2.** Moving OR analysis visualizing MS risk

**Figure S3.** Median MFI response against HHV-6B proteins IE1B and 101K at different ages.

**Figure S4.** Manhattan plots IgG serostatus

**Figure S5.** Manhattan plots of the association results when associated HLA alleles are added as covariates

**Figure S6.** Correlation between different antibody responses for HHV-6A and B antigens

**Figure S7.** Correlation between anti-HHV-6A/6B, CMV and EBV serology

**Figure S8.** Flow chart depicting identification and selection of patients for inclusion in the Pre-MS cohort

**Figure S9.** Alignment of amino acid sequences used in the multiplex serological assay.

**Table S1.** Difference in IgG response against investigated HHV-6A/6B proteins between MS cases and controls in the pre-MS cohort

**Table S2.** Difference in IgG response against investigated HHV-6A/6B proteins between MS cases and controls in the established MS cohort

**Table S3.** Association between IgG levels and HLA haplotypes and IE1B

**Table S4.** Association between IgG levels and HLA haplotypes and 101K

**Table S5.** Anti-IE1A, IE1B, 101K and p100 IgG response in children with primary HHV-6B infection

**Table S6.** Alignment of amino acid sequences used in the multiplex serological assay.

**Table S7**. Antibody responses analyzed for association with non-HLA SNPs serolevels

**Table S8**. Antibody responses analyzed for association with non-HLA SNPs serostatus

**Supplementary Figure S1**

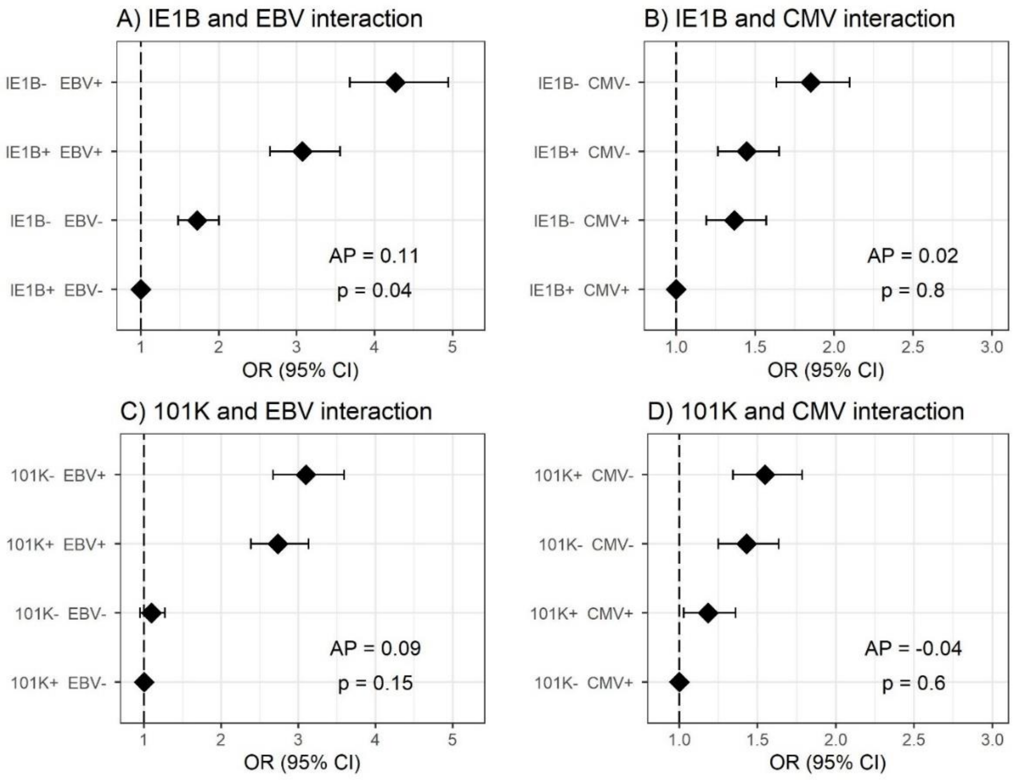


**Supplementary Figure S2**. **Interaction of antibody response against different herpesviruses in association to MS.** Odds ratios (OR) and confidence intervals (CI) for A) IE1B and EBV, B) IE1B and CMV, C) 101K and EBV, and D) 101K and CMV were obtained through logistic regression models adjusted for age, sex and cohort type analyzing the Established MS cohort (n=8742 persons with MS and n=7215 controls)**.** OR were calculated in relation to the group with the lowest MS risk. Plus (+) indicates being a strong responder while minus ( - ) indicates being a weak responder. Strong IE1B / 101K response is defined as having an MFI value being in the upper quartile of measured response, while a low response is having an antibody measurement being in the lower quartile of measured response. Strong EBV / CMV response is defined as having a higher EBV / CMV index than the median among controls, while a weak response is having a lower index compared to the median among controls.

**Supplementary Figure S2**

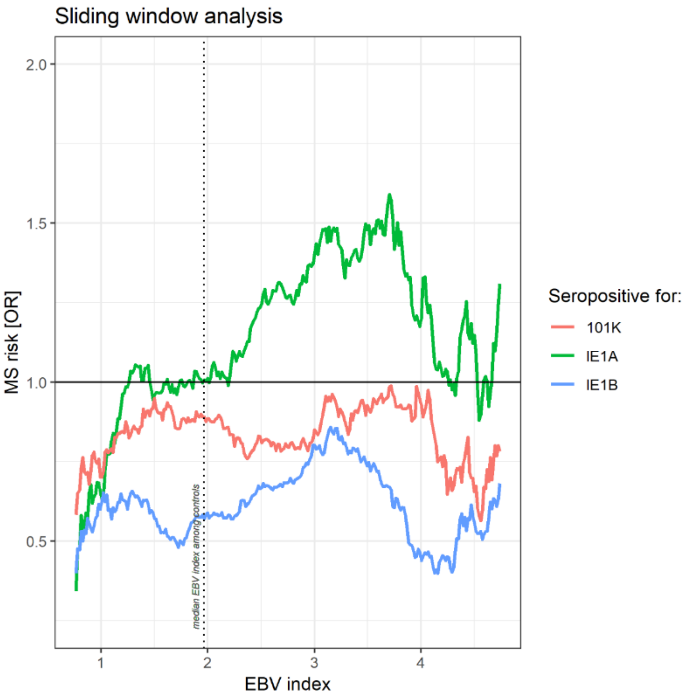


**Supplementary Figure S2. Moving OR analysis visualizing MS risk for high IE1A, IE1B and 101K responders, depending on EBV IgG response.** HHV-6A/-6B associated MS risk calculated using logistic regression analyses adjusted for age, sex and cohort type in a sliding window approach, i.e. stratified for EBV response in the established MS cohort (8742 persons with MS and 7215 controls). Strong responders are defined as those present in the highest quartile of each measured response, and in the regression models strong responders were compared to weak responders, here defined as those present in the lowest quartile of each measured response. Each analysis window contained a span of 0.75 units EBV index and overlapping analyses were performed as the window moved 0.015 units to its next analysis. Windows containing < 100 MS cases or < 100 controls were excluded which left 266 analysis windows, i.e. 266 plotted data points for each antigen response in the graph.

**Supplementary Figure S3**

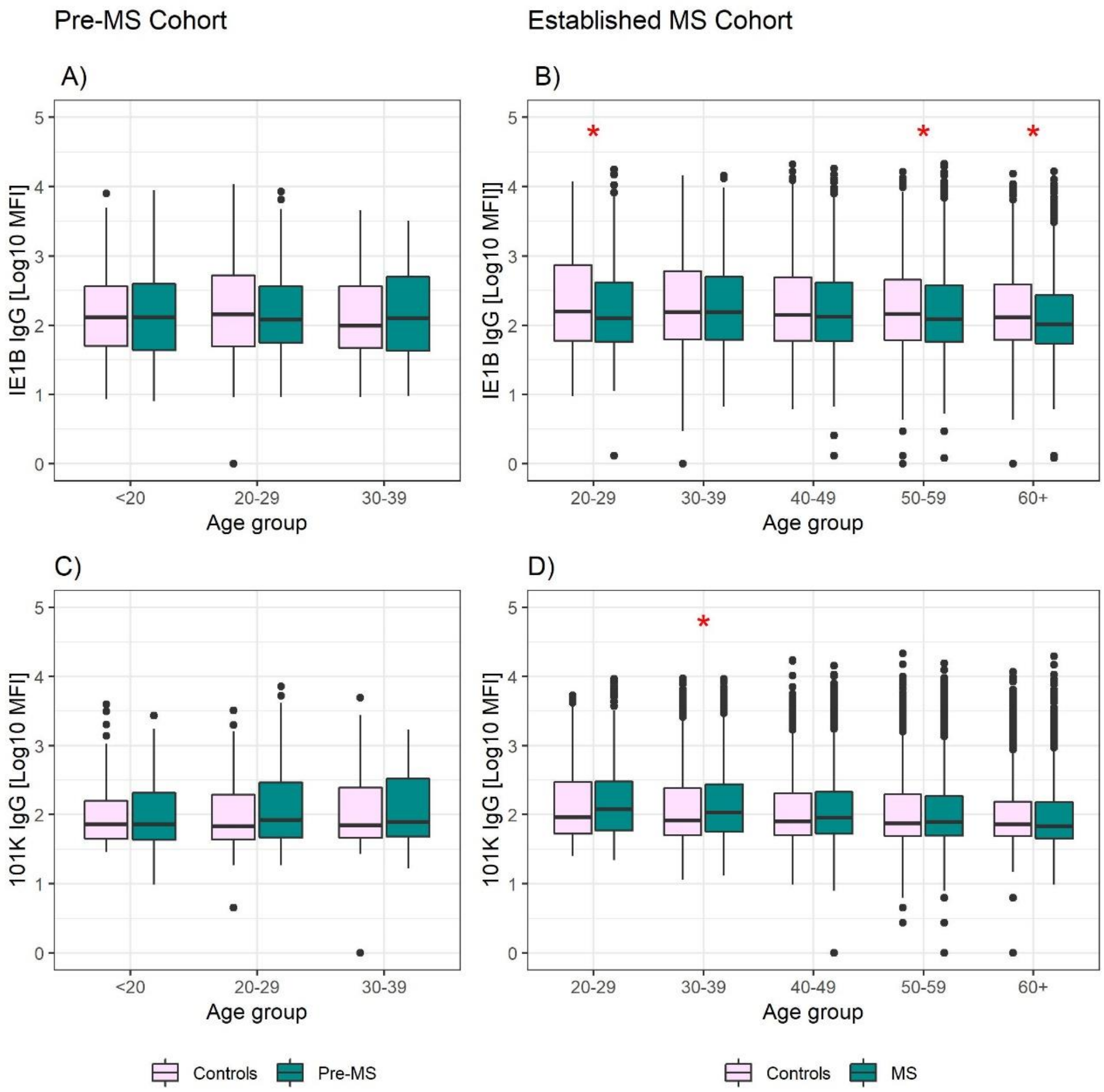


**Supplementary Figure S3. Median MFI response against HHV-6B proteins IE1B and 101K in different age groups.** Median of median fluorescence intensity (MFI) in different age groups for pre-MS cohort (n=478 persons) who later developed MS (green) and n=476 controls (pink); A, C) and established MS cohort (n=8394 persons with MS (green) and n=7214 controls (pink); B and D) for HHV-6B IE1B IgG (A and B); HHV-6B 101K IgG (C and D). Statistics were calculated with linear regression. Significant ($p<0.008$) differences in IgG levels between MS cases and controls within each age group are indicated with *.

**Supplementary Figure S4**

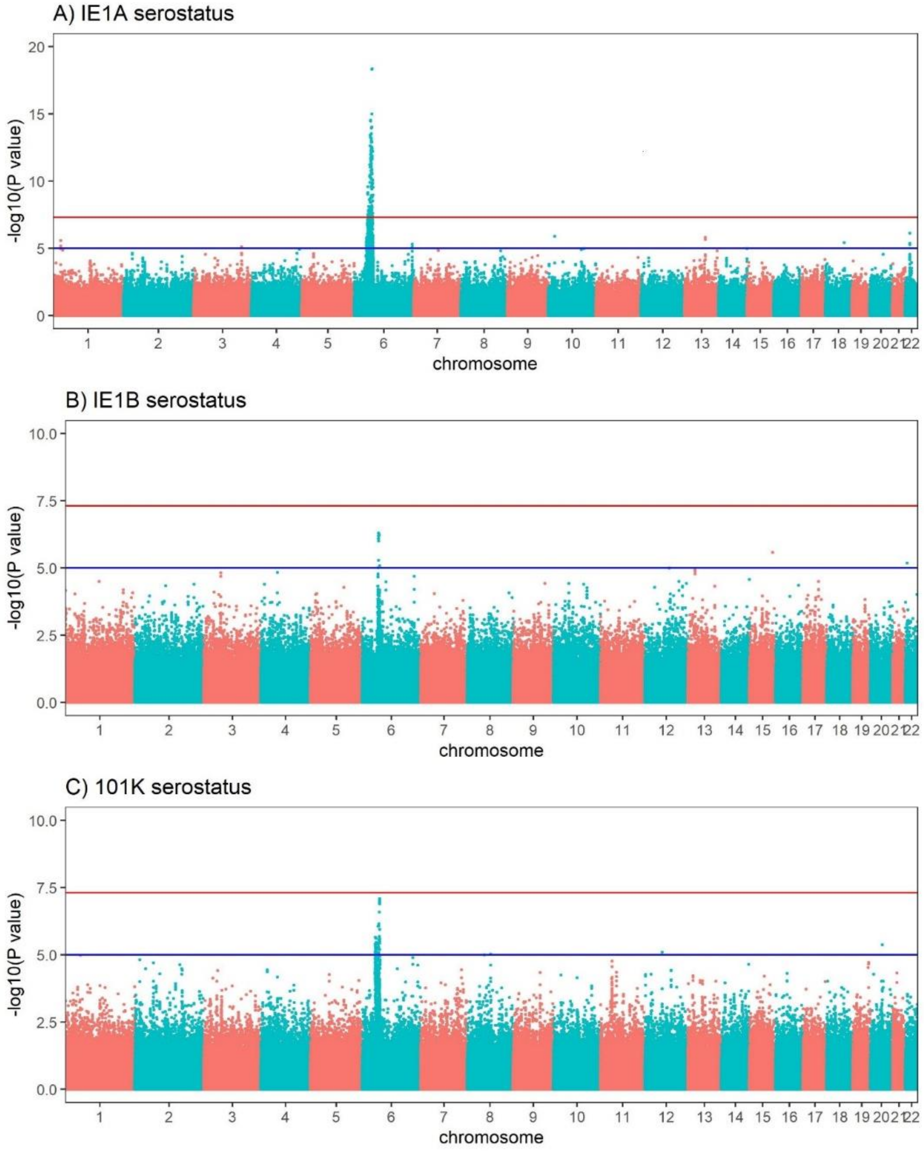


**Supplementary Figure S4. Manhattan plots visualizing associations between SNPs and anti-HHV-6A/6B protein IgG serostatus.** GWAS data showing associations between SNPs high/low IgG response, against (A) IE1A, (B) IE1B and (C) 101K. Red lines indicate GWAS significance level of $5x10^{-8}$ (-log10 = 7.3 on the y-axis) and blue lines indicate suggestive association ($p=10^{-5}$). Association analysis was carried out in established MS cohort with available genotypes (n=6,396 persons with MS and n=5,530 controls).

**Supplementary Figure S5**

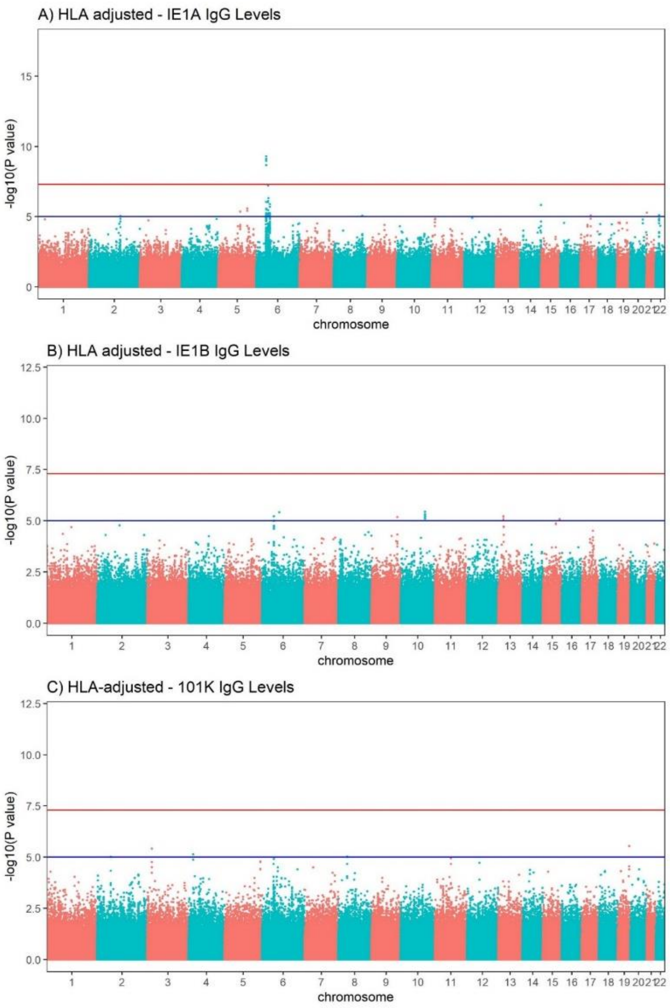


**Supplementary Figure S5. Manhattan plots of the association results when associated HLA alleles are added as covariates.** GWAS was performed between SNPs and A) IE1A, B) IE1B and C) 101K with linear regression models where HLA alleles associated to the responses were added to the model and adjusted for. Red lines indicate GWAS significance level of 5x10^-8 (-log10 = 7.3 on the y-axis) and blue lines indicate suggestive association ($p=10^{-5}$). In A) SNPs remaining significant after conditionen for HLA are rs1610678, rs1611149, rs173693 and rs915669. Association analysis was carried out in established MS cohort with available genotypes (n=6,396 persons with MS and n=5,530 controls).

**Supplementary Figure S6**

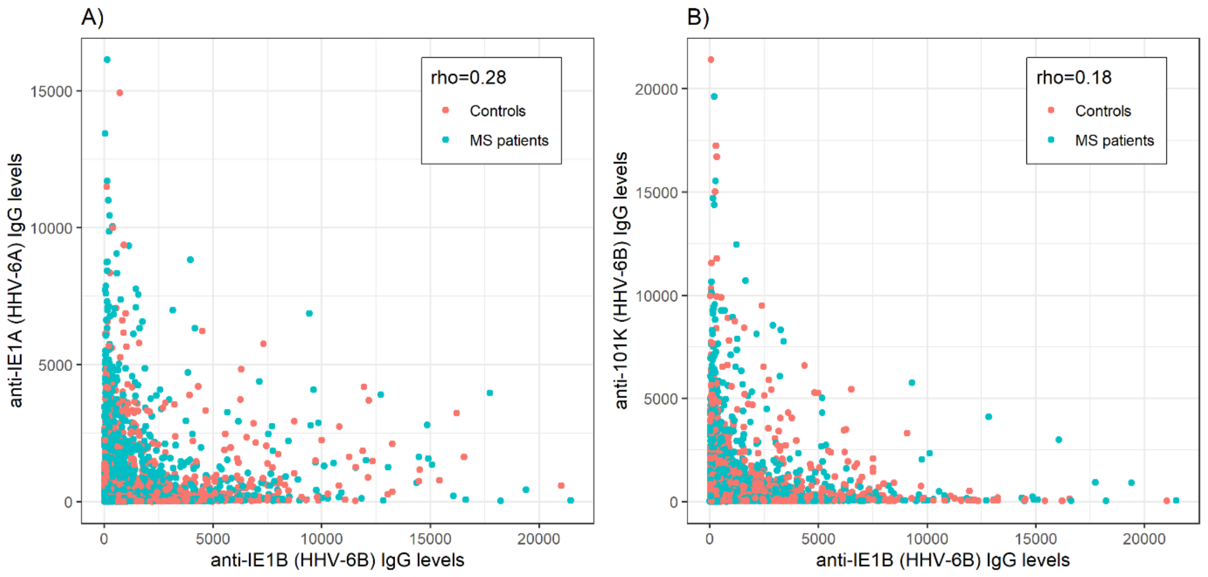


**Supplementary Figure S6. Correlation between different antibody responses for HHV-6A and B antigens.** Antibody levels measured as Median Fluorescence Intensity [MFI] against (A) IE1B and IE1A, (B) 101K and IE1B are plotted. Rho = Spearman's rank correlation coefficient. There is an inverse correlation between IgG levels against the different HHV-6 antigens, suggesting that different individuals respond with differently to the various antigens. Analysis was carried out in established MS cohort (n=8742 persons with MS and n=7215 controls).

**Supplementary Figure S7**

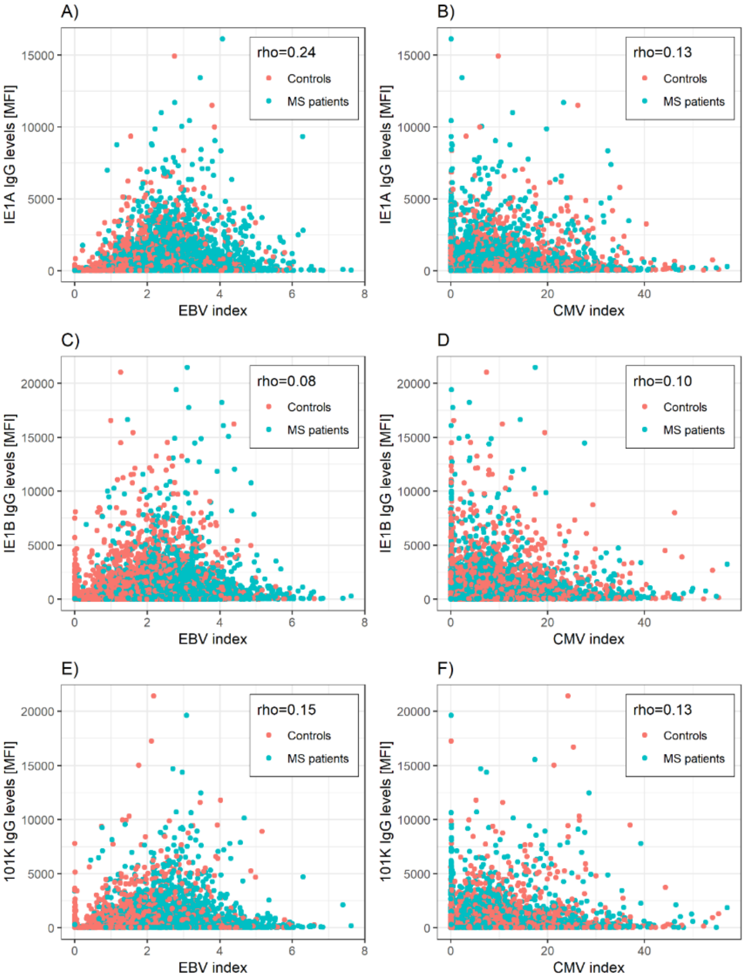


**Supplementary Figure S7. Correlation between anti-HHV-6A/6B, CMV and EBV serology.** Antibody levels [MFI] against IE1A, IE1B and 101K are plotted against EBV or CMV indexes. Rho = Spearman's rank correlation coefficient**.** There was a significant positive correlations between the different HHV-6 antibody responses and the EBV index (rho $\leq$ 0.23) as well as with the CMV index (rho $\leq$ 0.15). Being a strong IE1A responder was significantly associated with being a strong EBV responder, both in MS cases (OR = 3.41, p = $7x10^{-72}$) and in controls (OR = 2.68, p = $2x10^{-45}$). Analysis was carried out in established MS cohort (n=8742 persons with MS and n=7215 controls).

**Supplementary Figure S8**

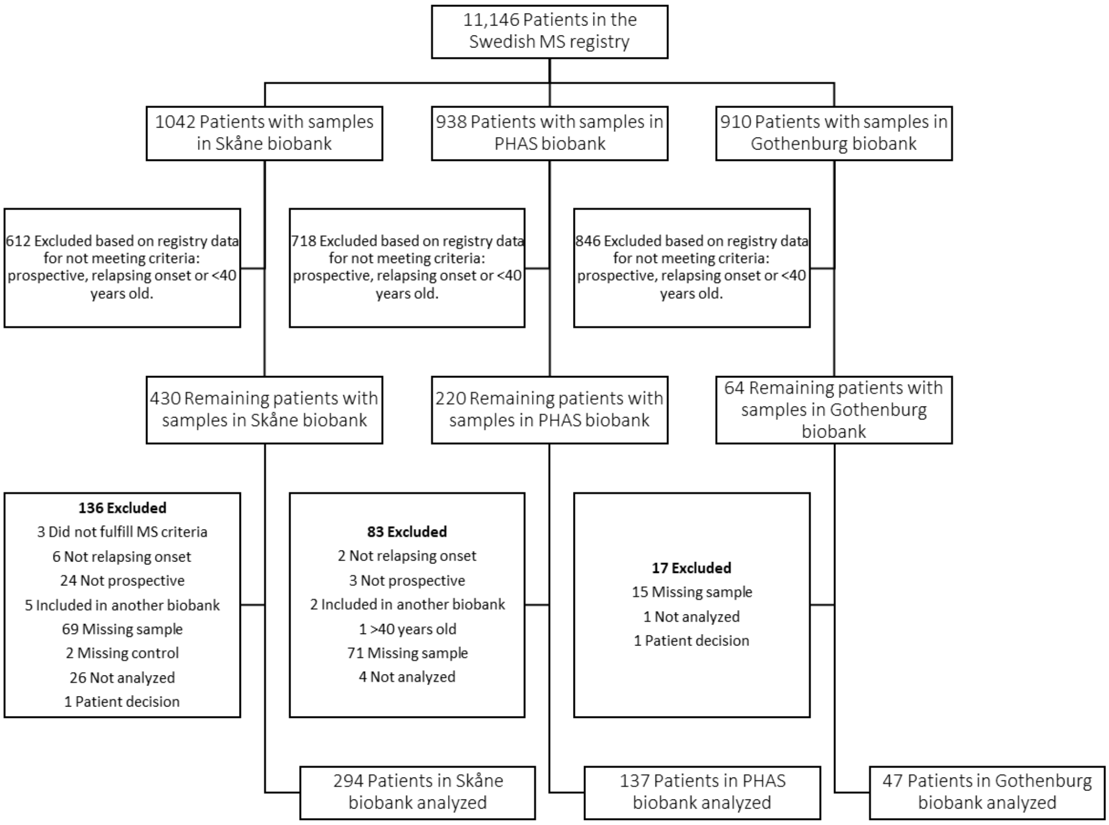


**Supplementary Figure S8. Flow chart depicting identification and selection of patients for inclusion in the Pre-MS cohort.** Through crosslinking between the Swedish MS registry and three Swedish microbiological biobanks potential study participants were identified. In the next step all patients that did not have a relapsing onset of MS, had not deposited serum sample before MS debut or were above 40 years old at the time of sampling were excluded. Validation of the information gathered from the Swedish MS registry was performed for 65 % of study participants. This, together with some samples missing or having too low volume, resulted in additional patients being excluded.

#### Supplementary Figure S9

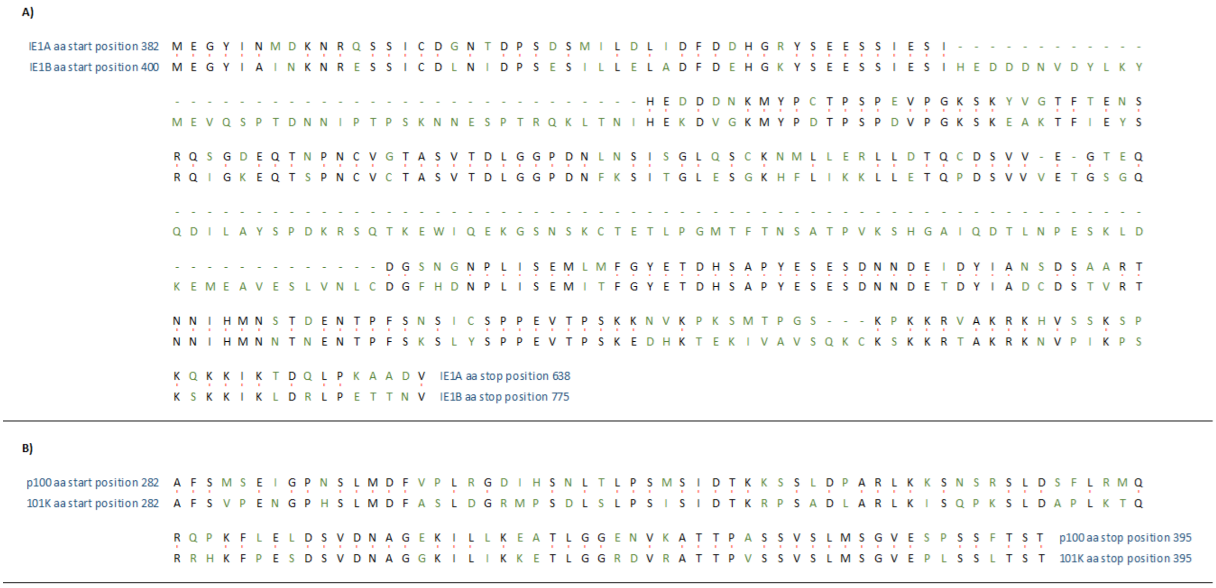


**Supplementary Figure S9. Alignment of amino acid sequences used in the multiplex serological assay.** Figure adapted from the NCBI BLAST. (A) Alignment of IE1A (Sequence ID = AGJ52065.1; length = 257 aa) with IE1B (NP_050266.1; length = 376 aa), (B) Alignment of p100 (NP_042902.1; length = 114 aa) with 101K (NP_050192.1; length = 114 aa). Identical amino acids from the two sequences are indicated with black font and a red connection between the amino acids. Green font indicates amino acid difference between the two sequences. aa = amino acid.

**Table S1. Difference in IgG response against investigated HHV-6A/6B proteins between MS cases and controls in the pre-MS cohort**

| | Model | - | Mann Whitney | Linear regression | | | | | | | | | |
|---|---|---|---|---|---|---|---|---|---|---|---|---|---|
| | Included | All measurements | | | | | | | | | | | |
| | Adjustment | - | | - | | AGE, SEX | | HLA | | EBV & CMV | | ALL | |
| | | Median MFI | p | β | p | β | p | β | p | β | p | β | p |
| IE1A | All subjects | 46 | **2E-08** | 0.18 | **7E-10** | 0.18 | **6E-10** | NA | | 0.14 | **3E-06** | 0.14 | **2E-06** |
| | Men | 46 | 4E-02 | 0.15 | 0.03 | 0.15 | 0.02 | | | 0.11 | 0.09 | 0.12 | 0.07 |
| | Women | 46 | **2E-07** | 0.19 | **8E-09** | 0.19 | **7E-09** | | | 0.14 | **2E-05** | 0.14 | **1E-05** |
| IE1B | All subjects | 125 | 8E-01 | -0.02 | 0.67 | -0.02 | 0.67 | NA | | -0.02 | 0.7 | -0.02 | 0.6 |
| | Men | 130 | 8E-01 | 0.00 | 0.98 | -0.01 | 0.95 | | | 0.04 | 0.7 | 0.04 | 0.7 |
| | Women | 124 | 7E-01 | -0.02 | 0.65 | -0.02 | 0.65 | | | -0.04 | 0.4 | -0.04 | 0.4 |
| 101K | All subjects | 75 | 3E-02 | 0.07 | 0.02 | 0.07 | 0.02 | NA | | 0.06 | 0.1 | 0.06 | 0.1 |
| | Men | 58 | 7E-01 | 0.08 | 0.28 | 0.08 | 0.28 | | | 0.09 | 0.2 | 0.09 | 0.2 |
| | Women | 78 | 4E-02 | 0.07 | 0.04 | 0.07 | 0.04 | | | 0.05 | 0.2 | 0.05 | 0.2 |

Mann Whitney U tests were performed on un-transformed MFI values but as the antibody levels can be affected by co-variables, the IgG responses were also investigated using linear regression models on Log10-transformed MFI values, adjusted for potential confounders. Measurements of EBV and CMV serology were converted into EBV and CMV indexes, respectively, and added as confounders. Significant p-values are highlighted in bold.

**Table S2. Difference in IgG response against investigated HHV-6A/6B proteins between MS cases and controls in the established MS cohort**

| | Model | - | Mann Whitney | Linear regression | | | | | | | | | |
|---|---|---|---|---|---|---|---|---|---|---|---|---|---|
| | Included | All measurements | | | | | | | | | | | |
| | Adjustment | - | | - | | AGE, SEX, COHORT | | HLA * | | EBV & CMV # | | ALL | |
| | | Median MFI | p | β | p | β | p | β | p | β | p | β | p |
| IE1A | All subjects | 47 | **6E-32** | 0.11 | **4E-33** | 0.10 | **9E-30** | 0.11 | **6E-26** | 0.05 | **7E-09** | 0.06 | **2E-10** |
| | Men | 51 | **8E-15** | 0.14 | **5E-15** | 0.15 | **5E-16** | 0.14 | **2E-11** | 0.08 | 2E-05 | 0.09 | **4E-06** |
| | Women | 47 | **3E-19** | 0.10 | **2E-20** | 0.09 | **2E-17** | 0.10 | **5E-16** | 0.04 | 4E-05 | 0.06 | **3E-06** |
| IE1B | All subjects | 123 | **1E-08** | - 0.06 | **2E-10** | -0.07 | **9E-14** | -0.07 | **6E-10** | -0.09 | **1E-17** | -0.09 | **3E-15** |
| | Men | 116 | 4E-02 | - 0.04 | 3E-02 | -0.05 | 4E-03 | -0.05 | 2E-02 | -0.07 | 3E-04 | -0.07 | 4E-04 |
| | Women | 125 | **1E-07** | - 0.07 | **6E-09** | -0.08 | **7E-12** | -0.07 | **3E-08** | -0.09 | **3E-14** | -0.09 | **4E-12** |
| 101K | All subjects | 46 | 7E-04 | 0.02 | 2E-02 | 0.02 | 4E-02 | 0.01 | 4E-01 | -0.002 | 8E-01 | -0.003 | 7E-01 |
| | Men | 35 | 2E-03 | 0.03 | 9E-02 | 0.02 | 2E-01 | 3E-04 | 0.99 | 0.01 | 6E-01 | -0.02 | 3E-01 |
| | Women | 51 | 6E-03 | 0.02 | 2E-02 | 0.02 | 7E-02 | 0.02 | 1E-01 | -0.001 | 9E-01 | 0.002 | 8E-01 |

Mann Whitney U tests were performed on un-transformed MFI values, but as the antibody levels can be affected by co-variables the IgG responses were also investigated using linear regression models on Log10-transformed MFI values, adjusted for potential confounders. *Models adjusted for carriage of HLA-DRB1*15:01, HLA-A*02 and HLA alleles associated to the respective measured IgG responses (marked in Table 2). # Measurements of EBV and CMV serology were converted into EBV and CMV indexes, respectively, and added as confounders. Number of individuals included were 15608 (pre-MS excluded), of which 4159 were men and 11449 were women. Significant p-values are highlighted.

**Table S3. Association between IgG levels and HLA haplotypes and IE1B**

| | **MS cases** | | | | **Controls** | | | | **All subjects** | | | |
|---|---|---|---|---|---|---|---|---|---|---|---|---|
| Adjustment[A] | Adjustment #1 | | Adjustment #2 | | Adjustment #1 | | Adjustment #2 | | Adjustment #1 | | Adjustment #2 | |
| | **β** | **p** | **β** | **p** | **β** | **p** | **β** | **p** | **β** | **p** | **β** | **p** |
| DRB1*13:02-DRB3*03:01-DQB1*06:04 | | | | | | | | | | | | |
| DRB1*13:02 | 0.082 | 0.002 | 0.076 | 0.004 | 0.078 | 0.003 | 0.079 | 0.002 | 0.079 | **2E-05** | 0.077 | **3E-05** |
| **DRB3*03:01** | 0.082 | 0.001 | 0.076 | 0.003 | 0.079 | 0.002 | 0.08 | 0.002 | 0.08 | **1E-05** | 0.078 | **2E-05** |
| DQB1*06:04 | 0.103 | **4E-04** | 0.097 | 0.001 | 0.071 | 0.009 | 0.074 | 0.007 | 0.084 | **2E-05** | 0.083 | **3E-05** |
| B*44:03-C*16:01-DRB1*07:01-DRB4*01:01-DQA1*02:01-DQB1*02:02 | | | | | | | | | | | | |
| **B*44:03** | 0.071 | 0.048 | 0.066 | 0.066 | 0.152 | **5E-05** | 0.153 | **5E-05** | 0.109 | **2E-05** | 0.107 | **4E-05** |
| C*16:01 | 0.071 | 0.107 | 0.064 | 0.146 | 0.125 | 0.008 | 0.123 | 0.009 | 0.096 | 0.003 | 0.092 | 0.004 |
| DRB1*07:01 | 0.004 | 0.853 | 0.003 | 0.89 | 0.098 | **8E-06** | 0.1 | **5E-06** | 0.053 | **6E-04** | 0.054 | **6E-04** |
| DRB4*01:01 | 0.024 | 0.393 | 0.022 | 0.43 | 0.144 | **7E-07** | 0.146 | **5E-07** | 0.083 | **3E-05** | 0.083 | **3E-05** |
| DQA1*02:01 | 0.016 | 0.467 | 0.015 | 0.497 | 0.098 | **6E-06** | 0.101 | **4E-06** | 0.06 | **1E-04** | 0.06 | **1E-04** |
| DQB1*02:02 | 0.011 | 0.691 | 0.008 | 0.753 | 0.149 | **6E-08** | 0.15 | **5E-08** | 0.081 | **2E-05** | 0.080 | **3E-05** |
| B*07:02-C*07:02 | | | | | | | | | | | | |
| **B*07:02** | -0.053 | **2E-04** | -0.047 | **1E-03** | -0.025 | 0.158 | -0.017 | 0.327 | -0.041 | **2E-04** | -0.035 | **2E-03** |
| C*07:02 | -0.045 | 0.001 | -0.04 | 0.005 | -0.028 | 0.105 | -0.02 | 0.245 | -0.038 | **5E-04** | -0.032 | **4E-03** |
| C*15:02 | | | | | | | | | | | | |
| C*15:02 | 0.003 | 0.917 | -0.005 | 0.857 | 0.148 | **8E-05** | 0.152 | **5E-05** | 0.062 | 0.008 | 0.059 | 0.012 |

The allele variants in a given HLA haplotype are presented. Underlined alleles were conferring the strongest effect, determined by stepwise conditional analyses. HLA genotype data was available for 7,063 MS cases and 6,098 controls. [A]Adjustment #1 = Age, sex, 5 PCA vectors, cohort type (incidence or prevalence). When adjusted for HLA, one associated HLA allele from each associated haplotype (underlined in the table) were added to the model in order to test for independent association of the investigated HLA allele. Adjustment #2 = Adjustment #1 + HLA. When all subjects were analyzed together, MS affection status were added as a covariable. Haplotypes associated to IE1B indicated in bold if any allele in it has a p-value <0.001.

**Table S4. Association between IgG levels and HLA haplotypes and 101K**

| | MS cases | | | | Controls | | | | All subjects | | | |
|---|---|---|---|---|---|---|---|---|---|---|---|---|
| Adjustment[A] | Adjustment #1 | | Adjustment #2 | | Adjustment #1 | | Adjustment #2 | | Adjustment #1 | | Adjustment #2 | |
| | **β** | **p** | **β** | **p** | **β** | **p** | **β** | **p** | **β** | **p** | **β** | **p** |
| B*40:01-DRB1*09:01 | | | | | | | | | | | | |
| **B*40:01** | -0.07 | **2E-04** | -0.06 | 0.001 | -0.07 | **1E-04** | -0.06 | **8E-04** | -0.07 | **6E-08** | -0.06 | **3E-06** |
| DRB1*09:01 | -0.06 | 0.158 | -0.05 | 0.234 | -0.11 | 0.003 | -0.10 | 0.006 | -0.09 | 0.001 | -0.08 | 0.004 |
| A*02:01-B*44:02-C*05:01 | | | | | | | | | | | | |
| **A*02:01** | -0.05 | **2E-04** | -0.04 | 0.000 | -0.03 | 0.012 | -0.03 | 0.010 | -0.04 | **6E-06** | -0.04 | **1E-05** |
| B*44:02 | -0.04 | 0.025 | -0.04 | 0.023 | -0.04 | 0.019 | -0.04 | 0.011 | -0.04 | **1E-03** | -0.04 | **5E-04** |
| C*05:01 | -0.04 | 0.031 | -0.04 | 0.025 | -0.05 | 0.003 | -0.05 | 0.002 | -0.05 | **3E-04** | -0.05 | **9E-05** |
| DRB1*04-DQA1*03:01-DQB1*03:02 | | | | | | | | | | | | |
| DRB1*04 | 0.06 | **1E-05** | 0.06 | **3E-06** | 0.02 | 0.245 | 0.02 | 0.074 | 0.04 | **8E-05** | 0.04 | **3E-06** |
| DQA1*03:01 | 0.06 | **2E-05** | 0.06 | **3E-06** | 0.01 | 0.303 | 0.02 | 0.104 | 0.04 | **2E-04** | 0.04 | **9E-06** |
| **DQB1*03:02** | 0.07 | **2E-06** | 0.07 | **4E-07** | 0.03 | 0.059 | 0.04 | 0.014 | 0.05 | **3E-06** | 0.05 | **9E-08** |

| | | | | | | | | | | | | |
|---|---|---|---|---|---|---|---|---|---|---|---|---|
| DRB1*13:02-DRB3*03:01-DQA1*01:02-DQB1*06:04 | | | | | | | | | | | | |
| DRB1*13:02 | -0.05 | 0.033 | -0.01 | 0.631 | -0.05 | 0.011 | -0.01 | 0.623 | -0.05 | **8E-04** | -0.01 | 0.479 |
| DRB3*03:01 | -0.05 | 0.036 | -0.01 | 0.648 | -0.05 | 0.013 | -0.01 | 0.661 | -0.05 | **1E-03** | -0.01 | 0.529 |
| DQA1*01:02 | -0.03 | 0.006 | -0.01 | 0.293 | -0.03 | 0.014 | -0.01 | 0.352 | -0.03 | 0.000 | -0.01 | 0.168 |
| **DQB1*06:04** | -0.07 | 0.004 | -0.03 | 0.327 | -0.06 | 0.004 | -0.02 | 0.437 | -0.07 | **4E-05** | -0.02 | 0.199 |
| DRB1*08:01-DQA1*04:01-DQB1*04:02 | | | | | | | | | | | | |
| DRB1*08:01 | 0.02 | 0.451 | 0.04 | 0.089 | 0.09 | **1E-04** | 0.10 | **1E-05** | 0.05 | 0.001 | 0.07 | **2E-05** |
| DQA*04:01 | 0.01 | 0.561 | 0.03 | 0.136 | 0.09 | **7E-05** | 0.10 | **8E-06** | 0.05 | 0.002 | 0.06 | **4E-05** |
| **DQB*04:02** | 0.02 | 0.294 | 0.04 | 0.051 | 0.09 | **5E-05** | 0.10 | **6E-06** | 0.05 | **5E-04** | 0.07 | **8E-06** |
| A*03:01 | | | | | | | | | | | | |
| A*03:01 | 0.03 | 0.023 | 0.02 | 0.122 | 0.04 | 0.011 | 0.03 | 0.064 | 0.03 | **6E-04** | 0.02 | 0.017 |
| C*03:03 | | | | | | | | | | | | |
| C*03:03 | 0.07 | **2E-04** | 0.06 | 0.003 | 0.04 | 0.037 | 0.03 | 0.094 | 0.06 | **5E-05** | 0.05 | 1E-03 |

The allele variants in a given HLA haplotype are presented. Underlined alleles were conferring the strongest effect, determined by stepwise conditional analyses. HLA genotype data was available for 7,063 MS cases and 6,098 controls. [A]Adjustment #1 = Age, sex, 5 PCA vectors, cohort type (incidence or prevalence). When adjusted for HLA, one associated HLA allele from each associated haplotype (underlined in the table) were added to the model in order to test for independent association of the investigated HLA allele. Adjustment #2 = Adjustment #1 + HLA. When all subjects were analyzed together, MS affection status were added as a covariable. Haplotypes associated to 101K are shown if any allele in it has a p-value <0.001 indicated in bold.

**Table S5. Anti-IE1A, IE1B, 101K and p100 IgG response in children with primary HHV-6B infection**

| | | | | | | | | | IE1A | | IE1B | | 101K | | p100 | |
|---|---|---|---|---|---|---|---|---|---|---|---|---|---|---|---|---|
| no | Age | Sex | Time of sampling * | Sample type | Isolation of HHV-6 from blood | Detection of HHV-6 DNA in plasma/ serum ^ | viral species | anti-HHV-6 IgG titer [IFA] ~ | MFI | Post/Pre | MFI | Post/Pre | MFI | Post/Pre | MFI | Post/Pre |
| 1 | 12 months | F | Acute | plasma | + | NT | B | 4 | 97 | | 26 | | 1 | | 3 | |
| | | | + 48 days | plasma | NT | NT | | 256 | 80 | 0.8 | 7 | 0.3 | 69 | **69.4** | 11 | 4.3 |
| 2 | 12 months | M | Acute | serum | + | NT | B | NT | 210 | | 50 | | 15 | | 5 | |
| | | | + 28 days | serum | NT | NT | | NT | 176 | 0.8 | 612 | **12.2** | 249 | **16.2** | 13 | 2.6 |
| 3 | 8 months | M | Acute | serum | + | NT | B | 4 | 91 | | 77 | | 3 | | 4 | |
| | | | + 16 days | serum | NT | NT | | 128 | 109 | 1.2 | 86 | 1.1 | 71 | **22.8** | 1 | 0.3 |
| 4 | 10 months | F | Acute | plasma | + | NT | B | 4 | 100 | | 74 | | 82 | | 10 | |
| | | | + 14 days | serum | NT | NT | | 128 | 106 | 1.1 | 26 | 0.4 | 186 | 2.3 | 11 | 1.1 |
| 5 | 14 months | F | Acute | plasma | + | + | B | 8 | 114 | | 42 | | 10 | | 1 | |
| | | | + 8 days | serum | NT | NT | | NT | 407 | 3.6 | 1466 | **35.3** | 101 | **10.5** | 10 | 9.5 |
| 6 | 14 months | F | Acute | plasma | NT | + | B | NT | 88 | | 16 | | 2 | | 14 | |
| | | | + 7 days | plasma | NT | NT | | 256 | 152 | 1.7 | 23 | 1.5 | 34 | **16.4** | 5 | 0.4 |
| 7 | 12 months | M | Acute | plasma | NT | + | B | 16 | 324 | | 105 | | 7 | | 29 | |
| | | | + 6 days | serum | NT | NT | | 256 | 190 | 0.6 | 71 | 0.7 | 35 | 4.9 | 1 | 0.04 |
| 8 | 15 months | M | Acute | serum | + | + | B | 8 | 252 | | 29 | | 1 | | 2 | |
| | | | + 6 days | serum | NT | NT | | 32 | 313 | 1.2 | 43 | 1.5 | 1 | 1.0 | 1 | 0.6 |
| 9 | 11 months | F | Acute | plasma | + | + | B | 4 | 140 | | 60 | | 75 | | 15 | |
| | | | + 6 days | plasma | NT | NT | | 16 | 104 | 0.7 | 29 | 0.5 | 56 | 0.7 | 15 | 1.0 |
| 10 | 12 months | M | Acute | plasma | + | + | B | NT | 74 | | 27 | | 3 | | 16 | |
| | | | + 5 days | plasma | NT | NT | | NT | 84 | 1.1 | 22 | 0.8 | 32 | 9.5 | 10 | 0.7 |

The first sample (Pre) is collected during Exanthema subitum (ES) acute phase, the second sample (Post) is collected in the convalescent phase and days between the two samples are given in the table. When ≤ 7 days between the samples, the samples are marked grey. NT = not tested. Isolation of viable HHV-6 from blood is described in (Asano Y, et al 1989 journal of pediatrics114:535-539). ^ Presence of DNA in cell-free plasma/serum was detected using the LAMP method (ref: Ihira 2004). If isolation of HHV-6 from blood sample was positive, the identification of HHV-6 species was performed by specific immunofluorescence staining with monoclonal antibodies to HHV-6 (Asano Y, et al 1989 journal of pediatrics114:535-539) and if detection of HHV-6 DNA from serum using LAMP was positive, the identification of HHV-6 species was followed RFLP method by LAMP products. (Ihira et al J virol Methods 2008 154(1-2):223-225). ~ IFA was used for detection of antibody against the representative strain of HHV-6B isolated (FG-1) from patients with ES (Suga et al 1989 pediatrics 83(6)1003-6.). A serum dilution of 1:100 was used in the assay and with that dilution MFI values <50 may be in the technical noise of the assay and should be interpreted with caution. The ability of this secondary antibody to detect IgM is limited, but we cannot be excluded

**Table S6. Amino acid sequences used as antigens in the multiplex serological assay.**

| Virus species | Antigen used (Sequence ID) | Protein sequence length | Amino acid start and stop position | Sequence | Most similar protein from another virus species (amino acid identity and Sequence ID) |
|---|---|---|---|---|---|
| **HHV-6A** | IE1A (AGJ52065.1) | 257 | 382-638 | MEGYINMDKNRQSSICDGNTDPSDSMILDLIDFDDHGRYSEESSI<br>ESIHEDDDNKMYPCTPSPEVPGKSKYVGTFTENSRQSGDEQTNP<br>NCVGTASVTDLGGPDNLNSISGLQSCKNMLLERLLDTQCDSVVE<br>GTEQDGSNGNPLISEMLMFGYETDHSAPYESESDNNDEIDYIAN<br>SDSAARTNNIHMNSTDENTPFSNSICSPPEVTPSKKNVKPKSMTP<br>GSKPKKRVAKRKHVSSKSPKQKKIKTDQLPKAADV | **IE1B**<br>(98/180 [54%] AAP81093.1 )<br>(97/176 [55%] NP_050266.1) |
| **HHV-6B** | IE1B (NP_050266.1) | 376 | 400-775 | MEGYIAINKNRESSICDLNIDPSESILLELADFDEHGKYSEESSIESI<br>HEDDDNVDYLKYMEVQSPTDNNIPTPSKNNESPTRQKLTNIHE<br>KDVGKMYPDTPSPDVPGKSKEAKTFIEYSRQIGKEQTSPNCVCT<br>ASVTDLGGPDNFKSITGLESGKHFLIKKLLETQPDSVVVETGSGQ<br>QDILAYSPDKRSQTKEWIQEKGSNSKCTETLPGMTFTNSATPVK<br>SHGAIQDTLNPESKLDKEMEAVESLVNLCDGFHDNPLISEMITF | **IE1A**<br>(94/173 [54%] CAA58338.1)<br>(93/173 [54%] AGJ52065.1) |

| | | | | | |
|---|---|---|---|---|---|
| | | | | GYETDHSAPYESESDNNDETDYIADCDSTVRTNNIHMNNTNE<br>NTPFSKSLYSPPEVTPSKEDHKTEKIVAVSQKCKSKKRTAKRKN<br>VPIKPSKSKKIKLDRLPETTNV | |
| **HHV-6B** | 101K (NP_050192.1) | 114 | 282-395 | AFSVPENGPHSLMDFASLDGRMPSDLSLPSISIDTKRPSADLA<br>RLKISQPKSLDAPLKTQRRHKFPESDSVDNAGGKILIKKETLGG<br>RDVRATTPVSSVSLMSGVEPLSSLTST | **p100**<br>(81/114 [71%] AKZ18118.1)<br>(74/114 [65%] NP_042902.1) |
| **HHV-6A** | p100 (NP_042902.1) | 114 | 282-395 | AFSMSEIGPNSLMDFVPLRGDIHSNLTLPSMSIDTKKSSLDPA<br>RLKKSNSRSLDSFLRMQRQPKFLELDSVDNAGEKILLKEATLG<br>GENVKATTPASSVSLMSGVESPSSFTST | **101K**<br>(74/114 [65%] NP_050192.1) |

Amino acid similarity to the top-scored protein from another virus species, obtained with the NCBI BLAST function. If more than one Sequence IDs are given, the second ID is of special interest. The % identity is given by the protein BLAST matching, please note that the matched amino acid length is not always the same as the length as the used protein sequence.

### Table S7. Antibody responses analyzed for association with non-HLA SNPs serolevels

| Antigen | Subject group | Chromosome | SNP marker | Basepair position | Minor allele | β | p |
|---|---|---|---|---|---|---|---|
| IE1A | MS cases | 2 | RS17450916 | 205 697 101 | A | -0.06911 | 2.58E-06 |
| IE1A | MS cases | 3 | RS17129314 | 197 031 608 | G | -0.12 | 7.50E-06 |
| IE1A | MS cases | 5 | RS6894855 | 111 149 480 | C | 0.05547 | 5.38E-06 |
| IE1A | MS cases | 6 | RS2247056 | 31 265 490 | A | -0.04979 | 3.50E-06 |
| IE1A | MS cases | 6 | RS2596487 | 31 325 056 | A | -0.0665 | 2.24E-06 |
| IE1A | MS cases | 6 | RS2523573 | 31 328 988 | C | -0.07431 | 2.92E-06 |
| IE1A | MS cases | 6 | RS9266669 | 31 348 077 | A | -0.06755 | 6.81E-06 |
| IE1A | MS cases | 6 | RS3094013 | 31 434 366 | A | -0.07616 | 1.21E-06 |
| IE1A | MS cases | 6 | RS3131618 | 31 434 621 | G | -0.0751 | 1.90E-06 |
| IE1A | MS cases | 6 | RS3131643 | 31 442 782 | A | -0.06924 | 2.03E-06 |
| IE1A | MS cases | 6 | RS3099844 | 31 448 976 | A | -0.08391 | 8.61E-08 |
| IE1A | MS cases | 6 | RS3094011 | 31 451 836 | G | -0.08265 | 1.61E-07 |
| IE1A | MS cases | 6 | RS9267445 | 31 483 481 | C | -0.07734 | 1.20E-06 |
| IE1A | MS cases | 6 | RS2734583 | 31 505 480 | G | -0.07186 | 3.96E-06 |
| IE1A | MS cases | 6 | RS3117582 | 31 620 520 | C | -0.07401 | 4.49E-06 |
| IE1A | MS cases | 6 | RS9267531 | 31 636 742 | G | -0.07392 | 4.59E-06 |
| IE1A | MS cases | 6 | **RS17201046** | 31 699 573 | A | 0.1805 | **9.19E-09** |
| IE1A | MS cases | 6 | RS3101018 | 31 705 864 | A | -0.0748 | 4.29E-06 |
| IE1A | MS cases | 6 | RS3132445 | 31 712 196 | A | -0.07315 | 5.91E-06 |
| IE1A | MS cases | 6 | RS3130484 | 31 715 882 | G | -0.07357 | 5.17E-06 |
| IE1A | MS cases | 6 | RS3131379 | 31 721 033 | A | -0.07301 | 6.32E-06 |
| IE1A | MS cases | 6 | RS3117574 | 31 725 230 | A | -0.07258 | 6.87E-06 |
| IE1A | MS cases | 6 | RS3131378 | 31 725 285 | G | -0.07354 | 5.25E-06 |
| IE1A | MS cases | 6 | RS3117577 | 31 727 474 | G | -0.07395 | 4.65E-06 |
| IE1A | MS cases | 6 | RS3101017 | 31 733 466 | G | -0.07217 | 7.77E-06 |
| IE1A | MS cases | 6 | RS2075798 | 31 846 741 | A | 0.088 | 4.54E-06 |
| IE1A | MS cases | 6 | RS555007 | 31 850 332 | G | 0.08493 | 8.65E-06 |
| IE1A | MS cases | 6 | RS558702 | 31 870 326 | A | -0.07658 | 2.10E-06 |
| IE1A | MS cases | 6 | RS519417 | 31 878 433 | A | -0.07899 | 1.07E-06 |
| IE1A | MS cases | 6 | RS497309 | 31 892 484 | C | -0.07807 | 1.28E-06 |
| IE1A | MS cases | 6 | RS1270942 | 31 918 860 | G | -0.07857 | 1.09E-06 |
| IE1A | MS cases | 6 | RS17201466 | 31 926 600 | A | 0.1494 | 1.80E-07 |
| IE1A | MS cases | 6 | **RS6455** | 32 006 896 | G | 0.1939 | **4.35E-09** |
| IE1A | MS cases | 6 | **RS11969759** | 32 021 130 | A | 0.1332 | **1.44E-08** |
| IE1A | MS cases | 6 | RS1150755 | 32 038 550 | A | -0.06753 | 2.02E-06 |
| IE1A | MS cases | 6 | **RS7774197** | 32 046 275 | C | 0.136 | **6.67E-09** |
| IE1A | MS cases | 6 | RS1150754 | 32 050 758 | A | -0.06824 | 1.51E-06 |
| IE1A | MS cases | 6 | RS1150753 | 32 059 867 | G | -0.07879 | 1.10E-06 |

| | | | | | | | |
|---|---|---|---|---|---|---|---|
| IE1A | MS cases | 6 | RS1150752 | 32 064 726 | G | -0.0783 | 1.47E-06 |
| IE1A | MS cases | 6 | RS1269852 | 32 080 191 | C | -0.07622 | 2.60E-06 |
| IE1A | MS cases | 6 | RS3130288 | 32 096 001 | A | -0.07667 | 2.13E-06 |
| IE1A | MS cases | 6 | RS3134942 | 32 168 771 | A | -0.07209 | 3.73E-06 |
| IE1A | MS cases | 6 | RS3131296 | 32 172 993 | A | -0.07384 | 2.43E-06 |
| IE1A | MS cases | 6 | RS206018 | 32 177 880 | C | 0.06652 | 1.35E-06 |
| IE1A | MS cases | 6 | RS3132956 | 32 179 438 | A | -0.07153 | 4.50E-06 |
| IE1A | MS cases | 6 | RS3131290 | 32 183 175 | A | -0.05254 | 2.57E-06 |
| IE1A | MS cases | 6 | RS915894 | 32 190 390 | C | -0.05629 | 8.88E-07 |
| IE1A | MS cases | 6 | RS6901158 | 32 205 942 | A | -0.07489 | 7.00E-07 |
| IE1A | MS cases | 6 | RS28895018 | 32 224 378 | G | 0.1245 | 3.11E-07 |
| IE1A | MS cases | 6 | RS3132971 | 32 230 256 | C | -0.0781 | 1.48E-06 |
| IE1A | MS cases | 6 | RS7775397 | 32 261 252 | C | -0.07617 | 2.54E-06 |
| IE1A | MS cases | 6 | RS6906662 | 32 266 506 | A | 0.1231 | 4.03E-07 |
| IE1A | MS cases | 6 | RS9268219 | 32 284 108 | C | -0.07356 | 9.65E-06 |
| IE1A | MS cases | 6 | RS9268235 | 32 290 208 | A | -0.07543 | 3.27E-06 |
| IE1A | MS cases | 6 | RS2073045 | 32 339 548 | A | -0.05299 | 3.27E-06 |
| IE1A | MS cases | 6 | **RS7758128** | 32 345 283 | A | 0.1643 | **2.40E-08** |
| IE1A | MS cases | 6 | RS2894254 | 32 345 689 | C | -0.07478 | 4.16E-06 |
| IE1A | MS cases | 6 | **RS17423649** | 32 357 133 | A | 0.0891 | **9.21E-09** |
| IE1A | MS cases | 6 | **RS17202259** | 32 357 489 | C | 0.08966 | **8.19E-09** |
| IE1A | MS cases | 6 | **RS16870123** | 32 359 460 | A | 0.08766 | **1.47E-08** |
| IE1A | MS cases | 6 | RS3817969 | 32 361 388 | A | 0.08306 | 6.62E-08 |
| IE1A | MS cases | 6 | **RS3763326** | 32 413 557 | G | 0.1593 | **3.18E-08** |
| IE1A | MS cases | 6 | RS9272219 | 32 602 269 | A | -0.05687 | 2.64E-06 |
| IE1A | MS cases | 6 | RS9273012 | 32 611 641 | G | -0.05651 | 3.06E-06 |
| IE1A | MS cases | 6 | RS17843604 | 32 620 283 | A | -0.05423 | 1.04E-06 |
| IE1A | MS cases | 6 | **RS9273349** | 32 625 869 | G | -0.06591 | **2.68E-09** |
| IE1A | MS cases | 6 | RS9273363 | 32 626 272 | A | -0.05966 | 3.54E-07 |
| IE1A | MS cases | 6 | RS6906021 | 32 626 311 | G | -0.05084 | 8.96E-06 |
| IE1A | MS cases | 6 | **RS1063355** | 32 627 714 | C | -0.06631 | **2.18E-09** |
| IE1A | MS cases | 6 | RS28986337 | 32 748 398 | A | 0.089 | 4.20E-06 |
| IE1A | MS cases | 6 | RS28986338 | 32 748 810 | A | 0.09066 | 2.71E-06 |
| IE1A | MS cases | 6 | RS28986359 | 32 751 349 | C | 0.08728 | 6.28E-06 |
| IE1A | MS cases | 6 | RS28986364 | 32 751 691 | C | 0.09062 | 3.08E-06 |
| IE1A | MS cases | 6 | RS28986373 | 32 752 150 | A | 0.08992 | 3.23E-06 |
| IE1A | MS cases | 6 | RS13204139 | 32 752 685 | A | 0.08955 | 3.93E-06 |
| IE1A | MS cases | 6 | RS28986396 | 32 754 430 | A | 0.08915 | 4.03E-06 |
| IE1A | MS cases | 6 | RS9276722 | 32 762 391 | G | 0.0887 | 4.32E-06 |
| IE1A | MS cases | 6 | RS11244 | 32 780 724 | A | -0.05508 | 4.62E-06 |
| IE1A | MS cases | 6 | RS2621321 | 32 789 480 | G | 0.06177 | 1.21E-06 |

| | | | | | | | |
|---|---|---|---|---|---|---|---|
| IE1A | MS cases | 6 | RS13209654 | 32 792 659 | G | 0.08885 | 4.18E-06 |
| IE1A | MS cases | 6 | RS2857101 | 32 794 676 | G | 0.06138 | 1.44E-06 |
| IE1A | MS cases | 6 | RS241456 | 32 795 965 | A | 0.05971 | 2.61E-06 |
| IE1A | MS cases | 6 | RS241454 | 32 796 144 | G | 0.0608 | 1.66E-06 |
| IE1A | MS cases | 6 | RS241453 | 32 796 226 | A | 0.06025 | 2.07E-06 |
| IE1A | MS cases | 6 | RS241448 | 32 796 685 | G | 0.0588 | 3.67E-06 |
| IE1A | MS cases | 6 | RS241447 | 32 796 751 | G | 0.06057 | 1.82E-06 |
| IE1A | MS cases | 6 | RS241446 | 32 796 967 | A | 0.06009 | 2.29E-06 |
| IE1A | MS cases | 6 | RS241445 | 32 797 072 | A | 0.06043 | 1.95E-06 |
| IE1A | MS cases | 6 | RS241441 | 32 797 297 | G | 0.06057 | 1.86E-06 |
| IE1A | MS cases | 6 | RS241440 | 32 797 361 | A | 0.06005 | 2.34E-06 |
| IE1A | MS cases | 6 | RS3819714 | 32 804 217 | A | 0.05551 | 5.45E-07 |
| IE1A | MS cases | 6 | RS3819715 | 32 804 219 | A | 0.0549 | 7.18E-07 |
| IE1A | MS cases | 6 | RS3819720 | 32 804 570 | A | -0.06002 | 1.35E-07 |
| IE1A | MS cases | 6 | RS241425 | 32 804 909 | G | -0.0526 | 5.28E-07 |
| IE1A | MS cases | 6 | RS241424 | 32 804 934 | A | -0.05336 | 4.68E-07 |
| IE1A | MS cases | 6 | RS2239701 | 32 805 049 | G | -0.04773 | 8.04E-06 |
| IE1A | MS cases | 6 | RS2071538 | 32 818 678 | A | 0.06102 | 3.64E-07 |
| IE1A | MS cases | 6 | RS1383266 | 32 834 732 | A | 0.05743 | 9.65E-07 |
| IE1A | MS cases | 6 | **RS423196** | 33 003 562 | G | -0.08783 | **3.55E-09** |
| IE1A | MS cases | 6 | **RS406477** | 33 005 644 | G | -0.08903 | **1.65E-09** |
| IE1A | MS cases | 6 | RS2395309 | 33 026 246 | G | -0.07036 | 2.65E-06 |
| IE1A | MS cases | 6 | RS17214519 | 33 032 188 | A | -0.07162 | 1.91E-06 |
| IE1A | MS cases | 6 | RS7905 | 33 032 975 | G | -0.08074 | 9.06E-07 |
| IE1A | MS cases | 6 | RS3077 | 33 033 022 | G | -0.07039 | 2.66E-06 |
| IE1A | MS cases | 6 | RS9469341 | 33 035 877 | G | -0.07277 | 1.26E-06 |
| IE1A | MS cases | 6 | RS1042190 | 33 036 999 | G | -0.07034 | 2.72E-06 |
| IE1A | MS cases | 6 | RS10214910 | 33 037 675 | A | -0.07121 | 2.20E-06 |
| IE1A | MS cases | 6 | RS2301224 | 33 038 369 | A | -0.07074 | 2.49E-06 |
| IE1A | MS cases | 6 | RS2301220 | 33 038 766 | A | -0.07256 | 1.36E-06 |
| IE1A | MS cases | 6 | RS6914849 | 33 040 715 | A | -0.07198 | 1.61E-06 |
| IE1A | MS cases | 6 | RS1431399 | 33 041 034 | G | -0.07134 | 2.03E-06 |
| IE1A | MS cases | 6 | RS1431400 | 33 041 176 | A | -0.07156 | 1.86E-06 |
| IE1A | MS cases | 6 | RS1431401 | 33 041 186 | A | -0.07145 | 1.93E-06 |
| IE1A | MS cases | 6 | **RS987870** | 33 042 880 | G | -0.09732 | **2.24E-08** |
| IE1A | MS cases | 6 | RS2071351 | 33 043 930 | G | -0.07137 | 2.02E-06 |
| IE1A | MS cases | 6 | RS2071353 | 33 044 257 | G | -0.08087 | 1.51E-07 |
| IE1A | MS cases | 6 | **RS2071354** | 33 044 388 | G | -0.09685 | **2.65E-08** |
| IE1A | MS cases | 6 | **RS1042151** | 33 048 661 | G | -0.08181 | **2.65E-10** |
| IE1A | MS cases | 6 | **RS9277378** | 33 050 279 | G | -0.07791 | **8.68E-12** |
| IE1A | MS cases | 6 | **RS9277386** | 33 050 499 | G | -0.08023 | **2.20E-12** |

| | | | | | | | |
|---|---|---|---|---|---|---|---|
| IE1A | MS cases | 6 | **RS9277396** | 33 051 139 | A | -0.07419 | **5.25E-11** |
| IE1A | MS cases | 6 | **RS9277426** | 33 051 910 | A | -0.07374 | **6.61E-11** |
| IE1A | MS cases | 6 | **RS9277469** | 33 053 468 | A | -0.0746 | **3.93E-11** |
| IE1A | MS cases | 6 | **RS9277471** | 33 053 682 | A | -0.07295 | **1.07E-10** |
| IE1A | MS cases | 6 | **RS9277533** | 33 054 721 | A | -0.07373 | **6.69E-11** |
| IE1A | MS cases | 6 | **RS9277546** | 33 055 346 | C | -0.07405 | **5.54E-11** |
| IE1A | MS cases | 6 | **RS9277554** | 33 055 538 | A | -0.07386 | **6.04E-11** |
| IE1A | MS cases | 6 | **RS9277555** | 33 055 605 | A | -0.07125 | **4.87E-10** |
| IE1A | MS cases | 6 | **RS3117228** | 33 056 435 | A | -0.07389 | **5.87E-11** |
| IE1A | MS cases | 6 | **RS3130188** | 33 057 176 | G | -0.07399 | **5.72E-11** |
| IE1A | MS cases | 6 | **RS3091282** | 33 057 198 | C | -0.07446 | **4.26E-11** |
| IE1A | MS cases | 6 | **RS3117226** | 33 057 659 | A | -0.07616 | **4.67E-11** |
| IE1A | MS cases | 6 | **RS3117225** | 33 057 711 | A | -0.07456 | **4.74E-11** |
| IE1A | MS cases | 6 | **RS3097652** | 33 057 835 | A | -0.07396 | **6.00E-11** |
| IE1A | MS cases | 6 | **RS1367730** | 33 058 114 | A | -0.0721 | **3.22E-10** |
| IE1A | MS cases | 6 | **RS3128972** | 33 058 774 | G | -0.07101 | **5.77E-10** |
| IE1A | MS cases | 6 | **RS2179920** | 33 058 874 | A | -0.07765 | **1.86E-10** |
| IE1A | MS cases | 6 | **RS2179919** | 33 059 262 | G | -0.0715 | **4.33E-10** |
| IE1A | MS cases | 6 | **RS3128917** | 33 059 996 | C | -0.07178 | **3.63E-10** |
| IE1A | MS cases | 6 | **RS3117222** | 33 060 949 | A | -0.07143 | **4.56E-10** |
| IE1A | MS cases | 6 | **RS3130190** | 33 061 690 | G | -0.0708 | **6.62E-10** |
| IE1A | MS cases | 6 | **RS3130191** | 33 061 871 | G | -0.07081 | **6.87E-10** |
| IE1A | MS cases | 6 | RS6914616 | 33 063 592 | A | -0.1296 | 5.52E-08 |
| IE1A | MS cases | 6 | **RS3130198** | 33 063 931 | G | -0.07189 | **3.30E-10** |
| IE1A | MS cases | 6 | **RS3117213** | 33 064 605 | A | -0.07187 | **3.56E-10** |
| IE1A | MS cases | 6 | RS2179915 | 33 065 734 | A | -0.13 | 5.22E-08 |
| IE1A | MS cases | 6 | **RS2395319** | 33 067 211 | A | -0.07192 | **3.44E-10** |
| IE1A | MS cases | 6 | **RS3117239** | 33 071 777 | G | -0.07716 | **2.19E-10** |
| IE1A | MS cases | 6 | **RS2064479** | 33 072 240 | A | -0.06877 | **2.49E-09** |
| IE1A | MS cases | 6 | **RS2064478** | 33 072 266 | A | -0.07746 | **1.89E-10** |
| IE1A | MS cases | 6 | **RS2064476** | 33 073 322 | G | -0.07383 | **6.05E-11** |
| IE1A | MS cases | 6 | **RS3117234** | 33 073 984 | G | -0.08027 | **2.72E-11** |
| IE1A | MS cases | 6 | **RS3117231** | 33 074 908 | G | -0.07594 | **4.85E-11** |
| IE1A | MS cases | 6 | **RS3117230** | 33 075 635 | G | -0.07713 | **2.21E-10** |
| IE1A | MS cases | 6 | **RS3128930** | 33 075 666 | A | -0.07784 | **7.12E-11** |
| IE1A | MS cases | 6 | RS6924545 | 33 077 607 | A | -0.1282 | 7.98E-08 |
| IE1A | MS cases | 9 | RS2578246 | 90 835 589 | G | -0.06121 | 2.45E-06 |
| IE1A | MS cases | 9 | RS893494 | 90 875 879 | G | -0.05975 | 1.49E-06 |
| IE1A | Controls | 5 | RS2304028 | 150 891 733 | A | 0.05312 | 1.63E-06 |
| IE1A | Controls | 5 | RS12717853 | 150 894 920 | A | 0.05378 | 1.25E-06 |

| | | | | | | | |
|---|---|---|---|---|---|---|---|
| IE1A | Controls | 5 | RS3822699 | 150 901 111 | A | 0.05304 | 1.77E-06 |
| IE1A | Controls | 5 | RS2053028 | 150 901 613 | A | 0.05455 | 9.35E-07 |
| IE1A | Controls | 6 | RS1633041 | 29 733 223 | A | -0.05464 | 1.63E-06 |
| IE1A | Controls | 6 | RS1737041 | 29 736 229 | A | -0.05459 | 1.65E-06 |
| IE1A | Controls | 6 | RS1737031 | 29 737 481 | A | -0.0549 | 1.01E-06 |
| IE1A | Controls | 6 | RS1615962 | 29 745 726 | A | -0.05446 | 1.88E-06 |
| IE1A | Controls | 6 | RS1633030 | 29 745 794 | G | -0.05497 | 9.69E-07 |
| IE1A | Controls | 6 | RS1633028 | 29 746 367 | C | -0.05477 | 1.09E-06 |
| IE1A | Controls | 6 | RS2735042 | 29 747 373 | G | -0.05473 | 1.07E-06 |
| IE1A | Controls | 6 | RS1002046 | 29 754 016 | A | -0.05449 | 1.74E-06 |
| IE1A | Controls | 6 | RS1610644 | 29 759 445 | A | -0.05421 | 1.97E-06 |
| IE1A | Controls | 6 | RS1611205 | 29 759 823 | G | -0.04935 | 7.90E-06 |
| IE1A | Controls | 6 | RS1610723 | 29 761 473 | C | -0.04881 | 9.45E-06 |
| IE1A | Controls | 6 | RS1633011 | 29 762 693 | A | -0.05455 | 1.84E-06 |
| IE1A | Controls | 6 | RS1630969 | 29 765 001 | A | -0.05697 | 9.53E-07 |
| IE1A | Controls | 6 | RS1610649 | 29 768 917 | G | -0.05527 | 3.41E-07 |
| IE1A | Controls | 6 | RS1610711 | 29 769 497 | C | -0.05469 | 1.65E-06 |
| IE1A | Controls | 6 | RS1632988 | 29 772 395 | A | -0.05667 | 1.24E-06 |
| IE1A | Controls | 6 | RS1632987 | 29 772 549 | G | -0.05521 | 1.31E-06 |
| IE1A | Controls | 6 | RS1632983 | 29 773 509 | A | -0.0543 | 1.11E-06 |
| IE1A | Controls | 6 | RS1736971 | 29 776 322 | A | -0.05446 | 1.78E-06 |
| IE1A | Controls | 6 | RS1736969 | 29 776 390 | A | -0.05538 | 1.22E-06 |
| IE1A | Controls | 6 | RS1736959 | 29 782 470 | A | -0.05341 | 3.49E-06 |
| IE1A | Controls | 6 | RS1736957 | 29 782 633 | G | -0.05266 | 2.71E-06 |
| IE1A | Controls | 6 | RS1077432 | 29 782 868 | G | -0.05392 | 2.27E-06 |
| IE1A | Controls | 6 | RS1620173 | 29 785 149 | C | -0.05463 | 1.65E-06 |
| IE1A | Controls | 6 | RS1619379 | 29 785 235 | A | -0.04883 | 3.89E-06 |
| IE1A | Controls | 6 | RS1610675 | 29 788 606 | A | -0.05209 | 3.64E-06 |
| IE1A | Controls | 6 | **RS1610678** | 29 789 190 | G | -0.05937 | **6.06E-09** |
| IE1A | Controls | 6 | **RS1611149** | 29 789 999 | A | -0.05959 | **5.15E-09** |
| IE1A | Controls | 6 | **RS1736936** | 29 794 317 | A | -0.06359 | **5.38E-10** |
| IE1A | Controls | 6 | **RS915669** | 29 798 425 | A | -0.06098 | **2.20E-09** |
| IE1A | Controls | 6 | RS1611133 | 29 809 382 | A | -0.06603 | 1.76E-07 |
| IE1A | Controls | 6 | RS1611750 | 29 814 778 | C | -0.06792 | 7.33E-08 |
| IE1A | Controls | 6 | RS2734985 | 29 818 662 | G | -0.05673 | 8.79E-06 |
| IE1A | Controls | 6 | RS2247504 | 29 819 077 | A | -0.05006 | 1.71E-06 |
| IE1A | Controls | 6 | RS2523756 | 29 820 717 | G | -0.05078 | 1.11E-06 |
| IE1A | Controls | 6 | RS2428510 | 29 823 027 | A | -0.05055 | 1.23E-06 |
| IE1A | Controls | 6 | RS2517854 | 29 823 296 | G | -0.0515 | 8.21E-07 |
| IE1A | Controls | 6 | RS2844826 | 29 823 319 | G | -0.05076 | 1.10E-06 |
| IE1A | Controls | 6 | RS6919513 | 29 823 994 | A | -0.05086 | 1.07E-06 |

| | | | | | | | |
|---|---|---|---|---|---|---|---|
| IE1A | Controls | 6 | RS1611684 | 29 825 846 | A | -0.05076 | 1.10E-06 |
| IE1A | Controls | 6 | RS1611717 | 29 829 577 | G | -0.05063 | 1.19E-06 |
| IE1A | Controls | 6 | RS1611732 | 29 831 008 | A | -0.05099 | 9.90E-07 |
| IE1A | Controls | 6 | RS765649 | 29 831 058 | A | -0.05003 | 1.69E-06 |
| IE1A | Controls | 6 | RS1611737 | 29 831 571 | G | -0.05309 | 3.55E-07 |
| IE1A | Controls | 6 | RS1362104 | 30 101 656 | A | -0.0514 | 7.91E-07 |
| IE1A | Controls | 6 | RS3130557 | 31 094 703 | A | -0.06998 | 8.09E-06 |
| IE1A | Controls | 6 | RS3132510 | 31 172 151 | G | -0.07342 | 6.09E-06 |
| IE1A | Controls | 6 | RS9263911 | 31 175 667 | G | 0.04637 | 4.69E-06 |
| IE1A | Controls | 6 | RS2844603 | 31 250 854 | A | -0.04853 | 3.88E-06 |
| IE1A | Controls | 6 | RS2853933 | 31 254 088 | A | -0.04808 | 4.66E-06 |
| IE1A | Controls | 6 | RS2524040 | 31 257 625 | A | -0.04897 | 3.13E-06 |
| IE1A | Controls | 6 | RS2524163 | 31 259 579 | G | -0.04794 | 5.02E-06 |
| IE1A | Controls | 6 | RS2524156 | 31 260 397 | A | -0.04751 | 6.22E-06 |
| IE1A | Controls | 6 | RS2243868 | 31 261 276 | A | -0.04812 | 4.73E-06 |
| IE1A | Controls | 6 | RS2247056 | 31 265 490 | A | -0.05518 | 9.07E-07 |
| IE1A | Controls | 6 | RS2853922 | 31 266 190 | A | -0.04879 | 3.52E-06 |
| IE1A | Controls | 6 | RS2524089 | 31 266 522 | C | -0.04907 | 3.04E-06 |
| IE1A | Controls | 6 | RS2524066 | 31 269 154 | A | -0.0488 | 3.35E-06 |
| IE1A | Controls | 6 | RS2523589 | 31 327 334 | C | -0.04523 | 5.55E-06 |
| IE1A | Controls | 6 | RS2844575 | 31 334 945 | G | 0.05234 | 2.13E-07 |
| IE1A | Controls | 6 | RS9266669 | 31 348 077 | A | -0.06755 | 4.06E-06 |
| IE1A | Controls | 6 | RS3131618 | 31 434 621 | G | -0.07055 | 9.24E-06 |
| IE1A | Controls | 6 | RS3099844 | 31 448 976 | A | -0.07435 | 2.02E-06 |
| IE1A | Controls | 6 | RS3094011 | 31 451 836 | G | -0.07683 | 9.80E-07 |
| IE1A | Controls | 6 | RS9267445 | 31 483 481 | C | -0.0729 | 4.42E-06 |
| IE1A | Controls | 6 | RS2734583 | 31 505 480 | G | -0.0715 | 4.51E-06 |
| IE1A | Controls | 6 | **RS805262** | 31 628 733 | A | -0.05813 | **1.26E-08** |
| IE1A | Controls | 6 | RS511294 | 31 888 869 | C | 0.1009 | 4.14E-06 |
| IE1A | Controls | 6 | RS498240 | 31 892 592 | A | 0.102 | 3.83E-06 |
| IE1A | Controls | 6 | RS2257331 | 31 898 285 | G | 0.1011 | 4.35E-06 |
| IE1A | Controls | 6 | RS9332735 | 31 901 773 | A | 0.1278 | 4.53E-06 |
| IE1A | Controls | 6 | RS635348 | 31 902 289 | C | 0.09978 | 6.23E-06 |
| IE1A | Controls | 6 | RS621701 | 31 903 121 | A | 0.09823 | 8.52E-06 |
| IE1A | Controls | 6 | **RS1042663** | 31 905 130 | A | 0.1002 | **4.90E-08** |
| IE1A | Controls | 6 | **RS2507953** | 31 905 328 | A | 0.1003 | **4.70E-08** |
| IE1A | Controls | 6 | RS550605 | 31 907 147 | G | 0.09945 | 6.21E-08 |
| IE1A | Controls | 6 | **RS653414** | 31 907 168 | A | 0.1003 | **4.70E-08** |
| IE1A | Controls | 6 | **RS497239** | 31 908 761 | G | 0.1003 | **4.70E-08** |
| IE1A | Controls | 6 | RS609061 | 31 910 162 | A | 0.09993 | 5.06E-08 |
| IE1A | Controls | 6 | **RS2242572** | 31 910 929 | A | 0.1004 | **4.28E-08** |

| | | | | | | | |
|---|---|---|---|---|---|---|---|
| IE1A | Controls | 6 | **RS547154** | 31 910 938 | A | 0.1003 | **4.53E-08** |
| IE1A | Controls | 6 | **RS541862** | 31 916 951 | G | 0.1009 | **3.92E-08** |
| IE1A | Controls | 6 | RS522162 | 31 919 917 | G | 0.1 | 5.11E-08 |
| IE1A | Controls | 6 | **RS760070** | 31 919 956 | G | 0.1006 | **4.31E-08** |
| IE1A | Controls | 6 | **RS550513** | 31 920 687 | A | 0.1006 | **4.35E-08** |
| IE1A | Controls | 6 | **RS403569** | 31 924 880 | A | 0.1009 | **4.00E-08** |
| IE1A | Controls | 6 | **RS438999** | 31 928 306 | G | 0.1001 | **4.61E-08** |
| IE1A | Controls | 6 | RS444921 | 31 932 177 | A | 0.07361 | 6.04E-06 |
| IE1A | Controls | 6 | RS406936 | 31 933 161 | A | 0.07374 | 5.79E-06 |
| IE1A | Controls | 6 | RS454212 | 31 934 372 | A | 0.07354 | 6.22E-06 |
| IE1A | Controls | 6 | RS453821 | 31 935 311 | A | 0.07322 | 6.80E-06 |
| IE1A | Controls | 6 | RS449643 | 31 936 679 | A | 0.07368 | 5.91E-06 |
| IE1A | Controls | 6 | RS387608 | 31 941 557 | A | 0.07303 | 6.95E-06 |
| IE1A | Controls | 6 | RS12333245 | 32 019 769 | A | 0.08628 | 9.04E-07 |
| IE1A | Controls | 6 | RS2269429 | 32 029 183 | A | 0.08671 | 8.64E-07 |
| IE1A | Controls | 6 | RS1150755 | 32 038 550 | A | -0.06335 | 6.31E-06 |
| IE1A | Controls | 6 | RS1150754 | 32 050 758 | A | -0.06325 | 6.50E-06 |
| IE1A | Controls | 6 | RS169496 | 32 052 983 | A | 0.08675 | 7.84E-07 |
| IE1A | Controls | 6 | RS204900 | 32 056 580 | C | 0.08665 | 8.05E-07 |
| IE1A | Controls | 6 | RS204899 | 32 057 627 | A | 0.08686 | 8.33E-07 |
| IE1A | Controls | 6 | RS204896 | 32 064 098 | A | 0.08419 | 1.84E-06 |
| IE1A | Controls | 6 | RS439844 | 32 072 940 | A | 0.09022 | 3.11E-07 |
| IE1A | Controls | 6 | **RS411337** | 32 077 380 | A | 0.1069 | **8.59E-09** |
| IE1A | Controls | 6 | **RS9469084** | 32 080 383 | A | 0.1041 | **2.11E-08** |
| IE1A | Controls | 6 | **RS204888** | 32 089 142 | A | 0.1039 | **2.22E-08** |
| IE1A | Controls | 6 | **RS169494** | 32 097 876 | A | 0.1057 | **1.31E-08** |
| IE1A | Controls | 6 | RS2071287 | 32 170 433 | A | -0.04708 | 3.46E-06 |
| IE1A | Controls | 6 | RS2071277 | 32 171 683 | G | -0.04538 | 6.59E-06 |
| IE1A | Controls | 6 | RS206017 | 32 176 211 | G | 0.04839 | 2.13E-06 |
| IE1A | Controls | 6 | RS206018 | 32 177 880 | C | 0.06599 | 2.34E-07 |
| IE1A | Controls | 6 | RS3131290 | 32 183 175 | A | -0.05024 | 1.33E-06 |
| IE1A | Controls | 6 | RS3134799 | 32 184 221 | A | -0.04906 | 1.56E-06 |
| IE1A | Controls | 6 | RS436388 | 32 186 264 | A | 0.04518 | 9.96E-06 |
| IE1A | Controls | 6 | RS379464 | 32 186 348 | A | 0.1063 | 3.17E-06 |
| IE1A | Controls | 6 | RS394657 | 32 187 023 | G | -0.04698 | 3.96E-06 |
| IE1A | Controls | 6 | RS8192584 | 32 189 192 | A | 0.122 | 2.76E-06 |
| IE1A | Controls | 6 | RS915894 | 32 190 390 | C | -0.05219 | 8.86E-07 |
| IE1A | Controls | 6 | RS3132971 | 32 230 256 | C | -0.07087 | 9.26E-06 |
| IE1A | Controls | 6 | **RS17843604** | 32 620 283 | G | 0.0553 | **4.80E-08** |
| IE1A | Controls | 6 | **RS9273349** | 32 625 869 | A | 0.06037 | **2.41E-09** |
| IE1A | Controls | 6 | **RS1063355** | 32 627 714 | A | 0.06106 | **1.74E-09** |

| | | | | | | | |
|---|---|---|---|---|---|---|---|
| IE1A | Controls | 12 | RS4765808 | 5 319 541 | A | -0.05014 | 8.44E-06 |
| IE1A | Controls | 16 | RS277903 | 25 175 122 | G | -0.05188 | 9.86E-06 |

| | | | | | | | |
|---|---|---|---|---|---|---|---|
| IE1A | All subjects | 0 | RS3997868 | 0 | G | -0.04484 | 5.94E-07 |
| IE1A | All subjects | 0 | **RS602875** | 0 | G | -0.05105 | **1.71E-08** |
| IE1A | All subjects | 1 | RS4659411 | 26 516 845 | G | -0.03239 | 8.87E-06 |
| IE1A | All subjects | 3 | RS433317 | 28 060 456 | A | 0.03335 | 5.30E-06 |
| IE1A | All subjects | 5 | RS6894855 | 111 149 480 | C | 0.04042 | 2.25E-06 |
| IE1A | All subjects | 5 | RS7718446 | 145 749 535 | G | 0.03621 | 5.03E-06 |
| IE1A | All subjects | 6 | RS444189 | 29 605 935 | C | -0.03411 | 6.22E-06 |
| IE1A | All subjects | 6 | RS396660 | 29 646 165 | A | -0.03648 | 3.82E-06 |
| IE1A | All subjects | 6 | RS445150 | 29 646 879 | G | -0.03616 | 4.60E-06 |
| IE1A | All subjects | 6 | RS2747430 | 29 648 506 | A | -0.03657 | 3.83E-06 |
| IE1A | All subjects | 6 | RS1610736 | 29 718 544 | G | 0.03287 | 6.52E-06 |
| IE1A | All subjects | 6 | RS909728 | 29 719 561 | A | 0.03303 | 5.90E-06 |
| IE1A | All subjects | 6 | RS1633041 | 29 733 223 | A | -0.03909 | 1.17E-06 |
| IE1A | All subjects | 6 | RS2735048 | 29 733 701 | A | 0.03855 | 6.02E-07 |
| IE1A | All subjects | 6 | RS2735046 | 29 734 098 | A | 0.03629 | 1.60E-06 |
| IE1A | All subjects | 6 | RS1737041 | 29 736 229 | A | -0.0391 | 1.16E-06 |
| IE1A | All subjects | 6 | RS1737031 | 29 737 481 | A | -0.03933 | 7.41E-07 |
| IE1A | All subjects | 6 | RS1362068 | 29 742 108 | G | -0.03777 | 1.04E-06 |
| IE1A | All subjects | 6 | RS1362070 | 29 742 299 | G | -0.03826 | 7.47E-07 |
| IE1A | All subjects | 6 | RS1615962 | 29 745 726 | A | -0.03845 | 1.81E-06 |
| IE1A | All subjects | 6 | RS1633030 | 29 745 794 | G | -0.03939 | 7.07E-07 |
| IE1A | All subjects | 6 | RS1633028 | 29 746 367 | C | -0.03932 | 7.51E-07 |
| IE1A | All subjects | 6 | RS2735042 | 29 747 373 | G | -0.03923 | 7.80E-07 |
| IE1A | All subjects | 6 | RS1002046 | 29 754 016 | A | -0.0391 | 1.17E-06 |
| IE1A | All subjects | 6 | RS1610641 | 29 759 066 | G | -0.03801 | 9.97E-07 |
| IE1A | All subjects | 6 | RS1610644 | 29 759 445 | A | -0.03918 | 1.10E-06 |
| IE1A | All subjects | 6 | RS1611205 | 29 759 823 | G | -0.03903 | 5.49E-07 |
| IE1A | All subjects | 6 | RS1610719 | 29 760 677 | G | -0.03823 | 8.49E-07 |
| IE1A | All subjects | 6 | RS1610723 | 29 761 473 | C | -0.03918 | 4.53E-07 |
| IE1A | All subjects | 6 | RS1610724 | 29 761 516 | G | -0.0386 | 6.65E-07 |
| IE1A | All subjects | 6 | RS1611220 | 29 761 835 | G | -0.03776 | 1.04E-06 |
| IE1A | All subjects | 6 | RS1633011 | 29 762 693 | A | -0.03889 | 1.39E-06 |
| IE1A | All subjects | 6 | RS1633005 | 29 764 472 | A | -0.04716 | 1.40E-06 |
| IE1A | All subjects | 6 | RS1633003 | 29 764 759 | G | -0.03733 | 1.37E-06 |
| IE1A | All subjects | 6 | RS1630969 | 29 765 001 | A | -0.03905 | 1.90E-06 |
| IE1A | All subjects | 6 | **RS1610649** | 29 768 917 | G | -0.04533 | **4.04E-09** |
| IE1A | All subjects | 6 | RS1610711 | 29 769 497 | C | -0.03927 | 1.06E-06 |
| IE1A | All subjects | 6 | RS1632995 | 29 769 621 | T | -0.03717 | 1.61E-06 |

| | | | | | | | |
|---|---|---|---|---|---|---|---|
| IE1A | All subjects | 6 | RS1610657 | 29 771 066 | G | -0.03771 | 1.07E-06 |
| IE1A | All subjects | 6 | RS1611192 | 29 771 879 | G | -0.04067 | 2.49E-06 |
| IE1A | All subjects | 6 | RS1632988 | 29 772 395 | A | -0.03931 | 1.86E-06 |
| IE1A | All subjects | 6 | RS1632987 | 29 772 549 | G | -0.03894 | 1.30E-06 |
| IE1A | All subjects | 6 | RS1632983 | 29 773 509 | A | -0.03864 | 1.02E-06 |
| IE1A | All subjects | 6 | RS1736976 | 29 773 999 | G | -0.03773 | 1.19E-06 |
| IE1A | All subjects | 6 | RS2523409 | 29 775 662 | G | 0.0362 | 2.13E-06 |
| IE1A | All subjects | 6 | RS1736971 | 29 776 322 | A | -0.03919 | 1.11E-06 |
| IE1A | All subjects | 6 | RS1736969 | 29 776 390 | A | -0.04005 | 6.56E-07 |
| IE1A | All subjects | 6 | RS1610663 | 29 778 626 | A | -0.03785 | 1.38E-06 |
| IE1A | All subjects | 6 | RS1610699 | 29 778 987 | C | -0.03696 | 2.22E-06 |
| IE1A | All subjects | 6 | RS1610669 | 29 780 093 | A | -0.03777 | 1.04E-06 |
| IE1A | All subjects | 6 | RS1736959 | 29 782 470 | A | -0.03742 | 4.21E-06 |
| IE1A | All subjects | 6 | RS1736957 | 29 782 633 | G | -0.03586 | 6.40E-06 |
| IE1A | All subjects | 6 | RS1077432 | 29 782 868 | G | -0.03939 | 1.02E-06 |
| IE1A | All subjects | 6 | RS1620173 | 29 785 149 | C | -0.03902 | 1.22E-06 |
| IE1A | All subjects | 6 | **RS1619379** | 29 785 235 | A | -0.04206 | **1.70E-08** |
| IE1A | All subjects | 6 | RS2735028 | 29 785 538 | A | -0.03604 | 3.17E-06 |
| IE1A | All subjects | 6 | RS1610675 | 29 788 606 | A | -0.03697 | 3.14E-06 |
| IE1A | All subjects | 6 | **RS1610678** | 29 789 190 | G | -0.05089 | **2.39E-12** |
| IE1A | All subjects | 6 | **RS1611149** | 29 789 999 | A | -0.05145 | **1.32E-12** |
| IE1A | All subjects | 6 | **RS1736936** | 29 794 317 | A | -0.05145 | **1.79E-12** |
| IE1A | All subjects | 6 | **RS915669** | 29 798 425 | A | -0.05191 | **8.45E-13** |
| IE1A | All subjects | 6 | RS1611133 | 29 809 382 | A | -0.0438 | 6.19E-07 |
| IE1A | All subjects | 6 | RS1611750 | 29 814 778 | C | -0.04473 | 3.60E-07 |
| IE1A | All subjects | 6 | RS2734985 | 29 818 662 | G | -0.04604 | 2.24E-07 |
| IE1A | All subjects | 6 | **RS2247504** | 29 819 077 | A | -0.04203 | **1.40E-08** |
| IE1A | All subjects | 6 | RS2523759 | 29 819 093 | G | -0.0448 | 4.25E-07 |
| IE1A | All subjects | 6 | RS3115627 | 29 820 278 | G | 0.03638 | 2.90E-06 |
| IE1A | All subjects | 6 | RS5013093 | 29 820 586 | A | -0.04509 | 3.54E-07 |
| IE1A | All subjects | 6 | **RS2523756** | 29 820 717 | G | -0.04332 | **4.51E-09** |
| IE1A | All subjects | 6 | RS2734981 | 29 821 692 | A | -0.04449 | 5.39E-07 |
| IE1A | All subjects | 6 | RS2734980 | 29 821 896 | A | -0.04509 | 3.58E-07 |
| IE1A | All subjects | 6 | RS2517861 | 29 821 982 | A | -0.04515 | 3.35E-07 |
| IE1A | All subjects | 6 | RS2517860 | 29 822 521 | G | -0.04592 | 2.10E-07 |
| IE1A | All subjects | 6 | **RS2428510** | 29 823 027 | A | -0.04286 | **6.65E-09** |
| IE1A | All subjects | 6 | **RS2517854** | 29 823 296 | G | -0.04405 | **2.54E-09** |
| IE1A | All subjects | 6 | **RS2844826** | 29 823 319 | G | -0.04306 | **5.48E-09** |
| IE1A | All subjects | 6 | **RS6919513** | 29 823 994 | A | -0.04305 | **5.67E-09** |
| IE1A | All subjects | 6 | RS1611674 | 29 825 434 | G | -0.03556 | 3.28E-06 |
| IE1A | All subjects | 6 | **RS1611684** | 29 825 846 | A | -0.04309 | **5.31E-09** |

| | | | | | | | |
|---|---|---|---|---|---|---|---|
| IE1A | All subjects | 6 | RS1611704 | 29 828 467 | A | 0.03688 | 2.21E-06 |
| IE1A | All subjects | 6 | **RS1611717** | 29 829 577 | G | -0.04279 | **6.85E-09** |
| IE1A | All subjects | 6 | **RS1611732** | 29 831 008 | A | -0.04301 | **5.74E-09** |
| IE1A | All subjects | 6 | **RS765649** | 29 831 058 | A | -0.04262 | **8.58E-09** |
| IE1A | All subjects | 6 | **RS1611737** | 29 831 571 | G | -0.04428 | **2.05E-09** |
| IE1A | All subjects | 6 | RS3132682 | 30 044 388 | C | 0.03468 | 1.74E-06 |
| IE1A | All subjects | 6 | RS7382061 | 30 047 965 | A | 0.03517 | 1.24E-06 |
| IE1A | All subjects | 6 | RS2057728 | 30 054 757 | A | 0.03569 | 9.30E-07 |
| IE1A | All subjects | 6 | RS6904455 | 30 055 377 | G | 0.03548 | 1.04E-06 |
| IE1A | All subjects | 6 | RS9261394 | 30 064 562 | A | 0.03766 | 2.14E-07 |
| IE1A | All subjects | 6 | RS1116222 | 30 071 279 | C | -0.03957 | 2.11E-06 |
| IE1A | All subjects | 6 | RS1116221 | 30 071 330 | A | -0.03906 | 2.96E-06 |
| IE1A | All subjects | 6 | RS1264341 | 30 802 465 | G | -0.05024 | 6.89E-06 |
| IE1A | All subjects | 6 | RS2535340 | 30 838 497 | G | -0.05072 | 5.64E-06 |
| IE1A | All subjects | 6 | RS1264322 | 30 857 894 | A | -0.0502 | 7.16E-06 |
| IE1A | All subjects | 6 | RS886422 | 30 864 279 | A | -0.05024 | 6.86E-06 |
| IE1A | All subjects | 6 | RS1049633 | 30 867 527 | A | -0.05014 | 7.32E-06 |
| IE1A | All subjects | 6 | RS1264312 | 30 872 982 | A | -0.0501 | 8.17E-06 |
| IE1A | All subjects | 6 | RS1264310 | 30 873 605 | A | -0.04992 | 8.44E-06 |
| IE1A | All subjects | 6 | RS1264308 | 30 879 987 | A | -0.05026 | 6.77E-06 |
| IE1A | All subjects | 6 | RS1264304 | 30 882 415 | A | -0.05049 | 6.15E-06 |
| IE1A | All subjects | 6 | RS3131921 | 30 907 335 | G | -0.04759 | 7.38E-06 |
| IE1A | All subjects | 6 | RS3094086 | 30 919 391 | A | -0.04967 | 5.34E-06 |
| IE1A | All subjects | 6 | RS3131934 | 30 931 844 | G | -0.05249 | 6.09E-07 |
| IE1A | All subjects | 6 | RS3131783 | 30 932 068 | A | -0.0532 | 5.18E-07 |
| IE1A | All subjects | 6 | RS1634721 | 30 977 680 | A | -0.05657 | 4.16E-07 |
| IE1A | All subjects | 6 | **RS3130544** | 31 058 340 | A | -0.06234 | **1.76E-08** |
| IE1A | All subjects | 6 | RS2233974 | 31 080 016 | C | -0.0488 | 6.24E-07 |
| IE1A | All subjects | 6 | RS3823402 | 31 081 743 | T | 0.04574 | 5.57E-06 |
| IE1A | All subjects | 6 | **RS3130557** | 31 094 703 | A | -0.06454 | **5.43E-09** |
| IE1A | All subjects | 6 | **RS7750641** | 31 129 310 | A | -0.06444 | **5.46E-09** |
| IE1A | All subjects | 6 | RS6921948 | 31 171 257 | C | 0.03708 | 2.92E-07 |
| IE1A | All subjects | 6 | **RS3132510** | 31 172 151 | G | -0.0692 | **1.83E-09** |
| IE1A | All subjects | 6 | RS2894181 | 31 174 527 | G | 0.03708 | 2.89E-07 |
| IE1A | All subjects | 6 | RS9263911 | 31 175 667 | G | 0.03965 | 5.12E-08 |
| IE1A | All subjects | 6 | RS3132505 | 31 177 503 | A | -0.03891 | 9.28E-06 |
| IE1A | All subjects | 6 | **RS3869109** | 31 184 196 | A | -0.04091 | **2.23E-08** |
| IE1A | All subjects | 6 | RS9263964 | 31 186 039 | G | -0.03983 | 5.11E-08 |
| IE1A | All subjects | 6 | RS2394895 | 31 206 979 | G | -0.03918 | 7.98E-06 |
| IE1A | All subjects | 6 | RS3868082 | 31 207 692 | A | -0.03914 | 1.05E-07 |
| IE1A | All subjects | 6 | RS4084262 | 31 218 889 | A | 0.04689 | 2.70E-06 |

| | | | | | | | |
|---|---|---|---|---|---|---|---|
| IE1A | All subjects | 6 | RS2394944 | 31 220 450 | A | -0.04118 | 3.92E-06 |
| IE1A | All subjects | 6 | RS1793891 | 31 221 698 | A | -0.04086 | 2.18E-06 |
| IE1A | All subjects | 6 | RS1986997 | 31 228 410 | A | -0.03911 | 1.02E-07 |
| IE1A | All subjects | 6 | RS2245822 | 31 230 800 | A | -0.04147 | 2.07E-06 |
| IE1A | All subjects | 6 | **RS1049281** | 31 236 567 | A | -0.04098 | **3.84E-08** |
| IE1A | All subjects | 6 | RS2844613 | 31 243 846 | A | -0.05071 | 5.59E-07 |
| IE1A | All subjects | 6 | **RS2524074** | 31 244 021 | G | -0.04413 | **5.88E-09** |
| IE1A | All subjects | 6 | **RS2844603** | 31 250 854 | A | -0.04116 | **2.32E-08** |
| IE1A | All subjects | 6 | **RS2853933** | 31 254 088 | A | -0.04086 | **2.77E-08** |
| IE1A | All subjects | 6 | **RS2524040** | 31 257 625 | A | -0.0417 | **1.47E-08** |
| IE1A | All subjects | 6 | **RS2524163** | 31 259 579 | G | -0.04091 | **2.69E-08** |
| IE1A | All subjects | 6 | **RS2524156** | 31 260 397 | A | -0.04072 | **3.26E-08** |
| IE1A | All subjects | 6 | **RS2243868** | 31 261 276 | A | -0.04094 | **2.68E-08** |
| IE1A | All subjects | 6 | RS2246954 | 31 265 262 | A | -0.03549 | 1.15E-06 |
| IE1A | All subjects | 6 | **RS2247056** | 31 265 490 | A | -0.05209 | **2.00E-11** |
| IE1A | All subjects | 6 | **RS2853922** | 31 266 190 | A | -0.04197 | **1.23E-08** |
| IE1A | All subjects | 6 | **RS2524089** | 31 266 522 | C | -0.04204 | **1.12E-08** |
| IE1A | All subjects | 6 | **RS2524066** | 31 269 154 | A | -0.04197 | **1.14E-08** |
| IE1A | All subjects | 6 | RS9264904 | 31 272 553 | A | 0.04298 | 1.39E-06 |
| IE1A | All subjects | 6 | RS3094691 | 31 274 693 | A | 0.03823 | 4.21E-07 |
| IE1A | All subjects | 6 | **RS2596487** | 31 325 056 | A | -0.06346 | **1.76E-10** |
| IE1A | All subjects | 6 | RS2523589 | 31 327 334 | A | 0.03841 | 9.16E-08 |
| IE1A | All subjects | 6 | **RS2523573** | 31 328 988 | C | -0.06997 | **5.93E-10** |
| IE1A | All subjects | 6 | RS2844575 | 31 334 945 | G | 0.03776 | 2.16E-07 |
| IE1A | All subjects | 6 | **RS9266669** | 31 348 077 | A | -0.06718 | **1.82E-10** |
| IE1A | All subjects | 6 | **RS3094013** | 31 434 366 | A | -0.07145 | **1.67E-10** |
| IE1A | All subjects | 6 | **RS3131618** | 31 434 621 | G | -0.07275 | **9.38E-11** |
| IE1A | All subjects | 6 | RS2518028 | 31 436 047 | A | -0.04051 | 5.95E-08 |
| IE1A | All subjects | 6 | RS1055569 | 31 440 082 | A | 0.03638 | 7.49E-06 |
| IE1A | All subjects | 6 | RS3828886 | 31 440 552 | C | 0.03735 | 9.62E-06 |
| IE1A | All subjects | 6 | **RS3131643** | 31 442 782 | A | -0.06235 | **7.85E-10** |
| IE1A | All subjects | 6 | **RS3099844** | 31 448 976 | A | -0.07919 | **1.03E-12** |
| IE1A | All subjects | 6 | RS2523706 | 31 451 567 | G | -0.03435 | 9.73E-06 |
| IE1A | All subjects | 6 | **RS3094011** | 31 451 836 | G | -0.07964 | **9.86E-13** |
| IE1A | All subjects | 6 | RS9267280 | 31 457 633 | A | 0.06073 | 6.24E-06 |
| IE1A | All subjects | 6 | RS2855812 | 31 472 720 | A | -0.04357 | 1.17E-07 |
| IE1A | All subjects | 6 | **RS9267445** | 31 483 481 | C | -0.07497 | **3.07E-11** |
| IE1A | All subjects | 6 | RS9469021 | 31 490 970 | A | 0.08315 | 4.33E-06 |
| IE1A | All subjects | 6 | RS3093988 | 31 492 453 | A | -0.04483 | 5.05E-06 |
| IE1A | All subjects | 6 | RS2516482 | 31 496 569 | G | -0.04383 | 7.24E-06 |
| IE1A | All subjects | 6 | **RS2734583** | 31 505 480 | G | -0.07134 | **1.10E-10** |

| | | | | | | | |
|---|---|---|---|---|---|---|---|
| IE1A | All subjects | 6 | RS3093542 | 31 540 693 | G | 0.08865 | 1.95E-06 |
| IE1A | All subjects | 6 | RS1800629 | 31 543 031 | A | -0.04322 | 9.40E-06 |
| IE1A | All subjects | 6 | **RS3117582** | 31 620 520 | C | -0.07007 | **8.78E-10** |
| IE1A | All subjects | 6 | **RS805262** | 31 628 733 | A | -0.05099 | **5.15E-12** |
| IE1A | All subjects | 6 | RS707920 | 31 629 096 | G | 0.04234 | 6.22E-07 |
| IE1A | All subjects | 6 | RS7029 | 31 629 953 | G | 0.04212 | 7.51E-07 |
| IE1A | All subjects | 6 | RS7992 | 31 630 241 | A | 0.04249 | 5.81E-07 |
| IE1A | All subjects | 6 | **RS9267531** | 31 636 742 | G | -0.06994 | **9.58E-10** |
| IE1A | All subjects | 6 | RS805290 | 31 648 403 | A | 0.04324 | 3.68E-07 |
| IE1A | All subjects | 6 | RS805282 | 31 659 731 | C | 0.04356 | 2.96E-07 |
| IE1A | All subjects | 6 | RS805281 | 31 661 489 | G | 0.0432 | 3.74E-07 |
| IE1A | All subjects | 6 | RS805277 | 31 662 957 | A | 0.04353 | 2.93E-07 |
| IE1A | All subjects | 6 | RS805274 | 31 665 194 | G | 0.04293 | 4.41E-07 |
| IE1A | All subjects | 6 | RS805287 | 31 678 730 | G | 0.03896 | 3.35E-06 |
| IE1A | All subjects | 6 | **RS17201046** | 31 699 573 | A | 0.115 | **2.30E-08** |
| IE1A | All subjects | 6 | **RS3101018** | 31 705 864 | A | -0.07111 | **6.34E-10** |
| IE1A | All subjects | 6 | **RS3132445** | 31 712 196 | A | -0.06921 | **1.43E-09** |
| IE1A | All subjects | 6 | **RS3130484** | 31 715 882 | G | -0.06951 | **1.19E-09** |
| IE1A | All subjects | 6 | **RS3131379** | 31 721 033 | A | -0.06897 | **1.71E-09** |
| IE1A | All subjects | 6 | **RS3117574** | 31 725 230 | A | -0.06895 | **1.61E-09** |
| IE1A | All subjects | 6 | **RS3131378** | 31 725 285 | G | -0.06967 | **1.11E-09** |
| IE1A | All subjects | 6 | **RS3117577** | 31 727 474 | G | -0.0698 | **1.03E-09** |
| IE1A | All subjects | 6 | **RS3101017** | 31 733 466 | G | -0.06891 | **1.69E-09** |
| IE1A | All subjects | 6 | RS707926 | 31 748 820 | A | 0.04336 | 3.78E-06 |
| IE1A | All subjects | 6 | RS707924 | 31 751 727 | A | 0.04265 | 5.44E-06 |
| IE1A | All subjects | 6 | RS480092 | 31 764 899 | G | 0.04337 | 3.72E-06 |
| IE1A | All subjects | 6 | **RS2075798** | 31 846 741 | A | 0.07719 | **4.87E-09** |
| IE1A | All subjects | 6 | **RS555007** | 31 850 332 | G | 0.07459 | **1.42E-08** |
| IE1A | All subjects | 6 | RS652888 | 31 851 234 | G | -0.04254 | 2.92E-06 |
| IE1A | All subjects | 6 | **RS558702** | 31 870 326 | A | -0.07072 | **6.01E-10** |
| IE1A | All subjects | 6 | **RS519417** | 31 878 433 | A | -0.07196 | **3.21E-10** |
| IE1A | All subjects | 6 | **RS511294** | 31 888 869 | C | 0.08785 | **4.80E-08** |
| IE1A | All subjects | 6 | **RS497309** | 31 892 484 | C | -0.07158 | **3.64E-10** |
| IE1A | All subjects | 6 | **RS498240** | 31 892 592 | A | 0.08946 | **3.36E-08** |
| IE1A | All subjects | 6 | **RS2257331** | 31 898 285 | G | 0.08886 | **3.92E-08** |
| IE1A | All subjects | 6 | **RS9332735** | 31 901 773 | A | 0.1145 | **4.64E-08** |
| IE1A | All subjects | 6 | RS635348 | 31 902 289 | C | 0.08785 | 6.21E-08 |
| IE1A | All subjects | 6 | RS621701 | 31 903 121 | A | 0.08796 | 5.50E-08 |
| IE1A | All subjects | 6 | **RS1042663** | 31 905 130 | A | 0.089 | **3.11E-11** |
| IE1A | All subjects | 6 | **RS2507953** | 31 905 328 | A | 0.08916 | **2.84E-11** |
| IE1A | All subjects | 6 | **RS550605** | 31 907 147 | G | 0.08912 | **2.93E-11** |

| | | | | | | | |
|---|---|---|---|---|---|---|---|
| IE1A | All subjects | 6 | **RS653414** | 31 907 168 | A | 0.08911 | **2.91E-11** |
| IE1A | All subjects | 6 | **RS497239** | 31 908 761 | G | 0.08849 | **3.86E-11** |
| IE1A | All subjects | 6 | **RS609061** | 31 910 162 | A | 0.08895 | **2.96E-11** |
| IE1A | All subjects | 6 | **RS2242572** | 31 910 929 | A | 0.08909 | **2.73E-11** |
| IE1A | All subjects | 6 | **RS547154** | 31 910 938 | A | 0.08901 | **2.87E-11** |
| IE1A | All subjects | 6 | **RS541862** | 31 916 951 | G | 0.08937 | **2.51E-11** |
| IE1A | All subjects | 6 | **RS1270942** | 31 918 860 | G | -0.07234 | **2.39E-10** |
| IE1A | All subjects | 6 | **RS522162** | 31 919 917 | G | 0.08855 | **3.77E-11** |
| IE1A | All subjects | 6 | **RS760070** | 31 919 956 | G | 0.08896 | **3.06E-11** |
| IE1A | All subjects | 6 | **RS550513** | 31 920 687 | A | 0.08955 | **2.38E-11** |
| IE1A | All subjects | 6 | **RS403569** | 31 924 880 | A | 0.08959 | **2.30E-11** |
| IE1A | All subjects | 6 | RS17201466 | 31 926 600 | A | 0.09887 | 1.42E-07 |
| IE1A | All subjects | 6 | RS440454 | 31 927 342 | A | -0.03654 | 1.40E-06 |
| IE1A | All subjects | 6 | **RS438999** | 31 928 306 | G | 0.08924 | **2.30E-11** |
| IE1A | All subjects | 6 | RS419788 | 31 928 799 | A | -0.03646 | 1.50E-06 |
| IE1A | All subjects | 6 | RS437179 | 31 929 014 | A | -0.03632 | 1.63E-06 |
| IE1A | All subjects | 6 | RS429608 | 31 930 462 | A | 0.05766 | 7.06E-08 |
| IE1A | All subjects | 6 | RS444921 | 31 932 177 | A | 0.05985 | 3.24E-07 |
| IE1A | All subjects | 6 | RS406936 | 31 933 161 | A | 0.05999 | 3.06E-07 |
| IE1A | All subjects | 6 | RS492899 | 31 933 518 | G | 0.06368 | 7.34E-06 |
| IE1A | All subjects | 6 | RS454212 | 31 934 372 | A | 0.05991 | 3.18E-07 |
| IE1A | All subjects | 6 | RS453821 | 31 935 311 | A | 0.05972 | 3.45E-07 |
| IE1A | All subjects | 6 | RS410851 | 31 936 668 | A | -0.03693 | 1.08E-06 |
| IE1A | All subjects | 6 | RS449643 | 31 936 679 | A | 0.05989 | 3.18E-07 |
| IE1A | All subjects | 6 | RS387608 | 31 941 557 | A | 0.05923 | 4.19E-07 |
| IE1A | All subjects | 6 | RS389883 | 31 947 460 | C | -0.0363 | 1.64E-06 |
| IE1A | All subjects | 6 | **RS6455** | 32 006 896 | G | 0.1333 | **1.32E-09** |
| IE1A | All subjects | 6 | RS12333245 | 32 019 769 | A | 0.06075 | 1.37E-06 |
| IE1A | All subjects | 6 | **RS11969759** | 32 021 130 | A | 0.08565 | **3.66E-08** |
| IE1A | All subjects | 6 | RS2269429 | 32 029 183 | A | 0.06028 | 1.77E-06 |
| IE1A | All subjects | 6 | **RS1150755** | 32 038 550 | A | -0.06558 | **6.14E-11** |
| IE1A | All subjects | 6 | **RS7774197** | 32 046 275 | C | 0.08616 | **2.88E-08** |
| IE1A | All subjects | 6 | **RS1150754** | 32 050 758 | A | -0.06593 | **4.64E-11** |
| IE1A | All subjects | 6 | RS169496 | 32 052 983 | A | 0.06026 | 1.65E-06 |
| IE1A | All subjects | 6 | RS204900 | 32 056 580 | C | 0.06043 | 1.55E-06 |
| IE1A | All subjects | 6 | RS204899 | 32 057 627 | A | 0.0614 | 1.22E-06 |
| IE1A | All subjects | 6 | **RS1150753** | 32 059 867 | G | -0.0717 | **3.71E-10** |
| IE1A | All subjects | 6 | RS204896 | 32 064 098 | A | 0.06018 | 1.97E-06 |
| IE1A | All subjects | 6 | **RS1150752** | 32 064 726 | G | -0.07329 | **1.89E-10** |
| IE1A | All subjects | 6 | RS439844 | 32 072 940 | A | 0.06215 | 8.39E-07 |
| IE1A | All subjects | 6 | **RS411337** | 32 077 380 | A | 0.07306 | **3.30E-08** |

| | | | | | | | |
|---|---|---|---|---|---|---|---|
| IE1A | All subjects | 6 | **RS1269852** | 32 080 191 | C | -0.06916 | **1.53E-09** |
| IE1A | All subjects | 6 | RS9469084 | 32 080 383 | A | 0.07175 | 5.80E-08 |
| IE1A | All subjects | 6 | RS204888 | 32 089 142 | A | 0.0716 | 6.20E-08 |
| IE1A | All subjects | 6 | **RS3130288** | 32 096 001 | A | -0.06985 | **9.75E-10** |
| IE1A | All subjects | 6 | **RS169494** | 32 097 876 | A | 0.0736 | **2.70E-08** |
| IE1A | All subjects | 6 | RS3134608 | 32 117 971 | C | -0.04683 | 9.10E-07 |
| IE1A | All subjects | 6 | RS2269425 | 32 123 639 | A | 0.05567 | 7.96E-08 |
| IE1A | All subjects | 6 | RS3096697 | 32 134 510 | A | -0.04712 | 8.05E-07 |
| IE1A | All subjects | 6 | RS3130347 | 32 134 656 | G | -0.04712 | 7.83E-07 |
| IE1A | All subjects | 6 | RS3130284 | 32 140 487 | G | -0.04722 | 7.67E-07 |
| IE1A | All subjects | 6 | RS408359 | 32 141 883 | A | 0.07315 | 1.23E-06 |
| IE1A | All subjects | 6 | RS3134947 | 32 145 205 | A | -0.04722 | 7.47E-07 |
| IE1A | All subjects | 6 | RS3134945 | 32 146 492 | A | -0.047 | 8.47E-07 |
| IE1A | All subjects | 6 | **RS3130349** | 32 147 696 | A | -0.05448 | **3.74E-08** |
| IE1A | All subjects | 6 | **RS1800625** | 32 152 442 | G | -0.05439 | **3.81E-08** |
| IE1A | All subjects | 6 | RS204995 | 32 154 285 | G | -0.04329 | 4.31E-06 |
| IE1A | All subjects | 6 | RS204993 | 32 155 581 | G | -0.04492 | 4.18E-07 |
| IE1A | All subjects | 6 | RS204992 | 32 156 908 | A | -0.04296 | 6.08E-06 |
| IE1A | All subjects | 6 | RS176095 | 32 158 319 | G | -0.04249 | 8.16E-06 |
| IE1A | All subjects | 6 | RS204991 | 32 161 366 | G | -0.04803 | 1.36E-06 |
| IE1A | All subjects | 6 | RS204990 | 32 161 430 | A | -0.04918 | 9.34E-07 |
| IE1A | All subjects | 6 | **RS2071278** | 32 165 444 | G | -0.06063 | **8.32E-09** |
| IE1A | All subjects | 6 | **RS3134942** | 32 168 771 | A | -0.06875 | **3.04E-10** |
| IE1A | All subjects | 6 | **RS2071287** | 32 170 433 | A | -0.04163 | **2.17E-08** |
| IE1A | All subjects | 6 | RS3132935 | 32 171 075 | G | -0.04722 | 8.87E-07 |
| IE1A | All subjects | 6 | **RS2071277** | 32 171 683 | G | -0.04267 | **8.14E-09** |
| IE1A | All subjects | 6 | **RS3131296** | 32 172 993 | A | -0.06988 | **1.89E-10** |
| IE1A | All subjects | 6 | **RS206017** | 32 176 211 | G | 0.04042 | **4.97E-08** |
| IE1A | All subjects | 6 | **RS206018** | 32 177 880 | C | 0.06657 | **1.62E-12** |
| IE1A | All subjects | 6 | **RS3132956** | 32 179 438 | A | -0.06847 | **3.64E-10** |
| IE1A | All subjects | 6 | **RS3131290** | 32 183 175 | A | -0.05167 | **1.52E-11** |
| IE1A | All subjects | 6 | **RS3134799** | 32 184 221 | A | -0.04677 | **4.86E-10** |
| IE1A | All subjects | 6 | RS436388 | 32 186 264 | A | 0.03911 | 1.40E-07 |
| IE1A | All subjects | 6 | **RS379464** | 32 186 348 | A | 0.1051 | **5.96E-10** |
| IE1A | All subjects | 6 | **RS394657** | 32 187 023 | G | -0.0447 | **2.46E-09** |
| IE1A | All subjects | 6 | RS1109771 | 32 187 605 | G | -0.03547 | 1.65E-06 |
| IE1A | All subjects | 6 | **RS8192584** | 32 189 192 | A | 0.1223 | **4.69E-10** |
| IE1A | All subjects | 6 | **RS915894** | 32 190 390 | C | -0.05418 | **4.83E-12** |
| IE1A | All subjects | 6 | RS3134930 | 32 191 620 | A | -0.04748 | 2.08E-07 |
| IE1A | All subjects | 6 | RS483574 | 32 194 956 | A | 0.04739 | 6.44E-06 |
| IE1A | All subjects | 6 | RS482759 | 32 195 017 | G | 0.04748 | 6.20E-06 |

| | | | | | | | |
|---|---|---|---|---|---|---|---|
| IE1A | All subjects | 6 | RS377763 | 32 199 144 | A | -0.04214 | 4.78E-06 |
| IE1A | All subjects | 6 | **RS6901158** | 32 205 942 | A | -0.06911 | **5.93E-11** |
| IE1A | All subjects | 6 | RS424232 | 32 208 324 | A | -0.04258 | 9.21E-07 |
| IE1A | All subjects | 6 | **RS28895018** | 32 224 378 | G | 0.08694 | **2.64E-08** |
| IE1A | All subjects | 6 | **RS3132971** | 32 230 256 | C | -0.07448 | **7.29E-11** |
| IE1A | All subjects | 6 | **RS7775397** | 32 261 252 | C | -0.07297 | **1.62E-10** |
| IE1A | All subjects | 6 | RS17422797 | 32 265 528 | A | 0.06858 | 7.21E-07 |
| IE1A | All subjects | 6 | **RS6906662** | 32 266 506 | A | 0.08641 | **3.17E-08** |
| IE1A | All subjects | 6 | **RS9268219** | 32 284 108 | C | -0.06752 | **7.72E-09** |
| IE1A | All subjects | 6 | **RS9268235** | 32 290 208 | A | -0.0728 | **1.88E-10** |
| IE1A | All subjects | 6 | **RS2073045** | 32 339 548 | A | -0.04301 | **3.97E-08** |
| IE1A | All subjects | 6 | **RS7758128** | 32 345 283 | A | 0.1299 | **1.27E-11** |
| IE1A | All subjects | 6 | **RS2894254** | 32 345 689 | C | -0.07211 | **3.15E-10** |
| IE1A | All subjects | 6 | **RS17423649** | 32 357 133 | A | 0.07176 | **4.71E-12** |
| IE1A | All subjects | 6 | **RS17202259** | 32 357 489 | C | 0.07204 | **4.39E-12** |
| IE1A | All subjects | 6 | RS3117099 | 32 358 270 | A | -0.03995 | 4.83E-06 |
| IE1A | All subjects | 6 | **RS16870123** | 32 359 460 | A | 0.07033 | **1.07E-11** |
| IE1A | All subjects | 6 | **RS3817969** | 32 361 388 | A | 0.06703 | **7.61E-11** |
| IE1A | All subjects | 6 | RS4248166 | 32 366 421 | G | 0.04613 | 6.15E-07 |
| IE1A | All subjects | 6 | RS2294884 | 32 367 259 | C | 0.04616 | 5.54E-07 |
| IE1A | All subjects | 6 | RS2294882 | 32 367 515 | G | 0.04329 | 2.33E-06 |
| IE1A | All subjects | 6 | RS2294881 | 32 367 604 | G | 0.04283 | 2.96E-06 |
| IE1A | All subjects | 6 | RS3763304 | 32 369 355 | A | 0.04604 | 5.88E-07 |
| IE1A | All subjects | 6 | **RS3129843** | 32 395 726 | G | -0.07057 | **7.62E-10** |
| IE1A | All subjects | 6 | **RS3763326** | 32 413 557 | G | 0.1189 | **1.63E-10** |
| IE1A | All subjects | 6 | RS9268877 | 32 431 147 | A | 0.03788 | 4.28E-07 |
| IE1A | All subjects | 6 | RS9268979 | 32 435 044 | A | 0.0381 | 4.15E-07 |
| IE1A | All subjects | 6 | **RS9270665** | 32 566 232 | A | -0.05394 | **5.96E-09** |
| IE1A | All subjects | 6 | RS660895 | 32 577 380 | G | -0.04462 | 6.62E-07 |
| IE1A | All subjects | 6 | **RS642093** | 32 582 075 | A | -0.05064 | **2.36E-08** |
| IE1A | All subjects | 6 | RS17533090 | 32 590 722 | A | 0.04805 | 8.44E-06 |
| IE1A | All subjects | 6 | **RS3129763** | 32 590 925 | A | -0.05371 | **6.96E-09** |
| IE1A | All subjects | 6 | **RS9272219** | 32 602 269 | A | -0.05302 | **2.81E-10** |
| IE1A | All subjects | 6 | RS2187668 | 32 605 884 | A | -0.05422 | 7.17E-07 |
| IE1A | All subjects | 6 | **RS9273012** | 32 611 641 | G | -0.05296 | **2.93E-10** |
| IE1A | All subjects | 6 | **RS17843604** | 32 620 283 | A | -0.05466 | **4.08E-13** |
| IE1A | All subjects | 6 | RS9273327 | 32 623 223 | C | -0.05349 | 1.31E-06 |
| IE1A | All subjects | 6 | **RS9273349** | 32 625 869 | G | -0.06273 | **7.47E-17** |
| IE1A | All subjects | 6 | RS6928482 | 32 626 249 | G | -0.0375 | 1.49E-06 |
| IE1A | All subjects | 6 | **RS9273363** | 32 626 272 | A | -0.05312 | **4.22E-11** |
| IE1A | All subjects | 6 | **RS6906021** | 32 626 311 | G | -0.04782 | **8.20E-10** |

| | | | | | | | |
|---|---|---|---|---|---|---|---|
| IE1A | All subjects | 6 | **RS1063355** | 32 627 714 | C | -0.06328 | **4.43E-17** |
| IE1A | All subjects | 6 | **RS9273448** | 32 627 747 | A | 0.04561 | **5.10E-09** |
| IE1A | All subjects | 6 | RS2854275 | 32 628 428 | A | -0.05527 | 4.28E-07 |
| IE1A | All subjects | 6 | RS9274741 | 32 637 994 | G | -0.03668 | 1.90E-06 |
| IE1A | All subjects | 6 | RS9275141 | 32 651 117 | C | -0.04201 | 5.90E-08 |
| IE1A | All subjects | 6 | RS2856695 | 32 651 894 | G | -0.04158 | 8.16E-08 |
| IE1A | All subjects | 6 | RS3021058 | 32 652 359 | C | -0.04209 | 5.71E-08 |
| IE1A | All subjects | 6 | RS4947342 | 32 653 070 | A | -0.04391 | 6.27E-07 |
| IE1A | All subjects | 6 | RS2856692 | 32 653 385 | C | -0.0442 | 5.48E-07 |
| IE1A | All subjects | 6 | **RS4642516** | 32 657 543 | A | -0.04238 | **4.56E-08** |
| IE1A | All subjects | 6 | RS5000634 | 32 663 564 | G | -0.04323 | 5.47E-08 |
| IE1A | All subjects | 6 | **RS6457622** | 32 664 163 | C | -0.04395 | **3.22E-08** |
| IE1A | All subjects | 6 | RS9275312 | 32 665 728 | G | -0.04889 | 3.61E-07 |
| IE1A | All subjects | 6 | **RS1794282** | 32 666 526 | A | -0.06743 | **4.49E-09** |
| IE1A | All subjects | 6 | RS9275328 | 32 666 822 | A | -0.0482 | 5.31E-07 |
| IE1A | All subjects | 6 | RS9275330 | 32 666 875 | G | -0.04836 | 4.91E-07 |
| IE1A | All subjects | 6 | RS9275333 | 32 666 968 | G | -0.04881 | 3.81E-07 |
| IE1A | All subjects | 6 | **RS3135006** | 32 667 119 | A | 0.04601 | **4.10E-09** |
| IE1A | All subjects | 6 | RS4947344 | 32 677 846 | A | 0.03882 | 2.77E-07 |
| IE1A | All subjects | 6 | RS9275601 | 32 682 664 | A | -0.03613 | 1.79E-06 |
| IE1A | All subjects | 6 | RS3873444 | 32 682 724 | A | 0.06443 | 3.65E-07 |
| IE1A | All subjects | 6 | RS1383265 | 32 739 888 | G | 0.04684 | 2.89E-07 |
| IE1A | All subjects | 6 | RS2857212 | 32 740 411 | G | 0.03882 | 3.48E-07 |
| IE1A | All subjects | 6 | **RS28986337** | 32 748 398 | A | 0.07994 | **2.51E-09** |
| IE1A | All subjects | 6 | **RS28986338** | 32 748 810 | A | 0.0806 | **1.83E-09** |
| IE1A | All subjects | 6 | **RS28986359** | 32 751 349 | C | 0.07823 | **5.88E-09** |
| IE1A | All subjects | 6 | **RS28986364** | 32 751 691 | C | 0.08022 | **2.76E-09** |
| IE1A | All subjects | 6 | **RS9276689** | 32 751 962 | A | -0.06756 | **1.94E-08** |
| IE1A | All subjects | 6 | **RS28986373** | 32 752 150 | A | 0.08036 | **2.02E-09** |
| IE1A | All subjects | 6 | **RS13204139** | 32 752 685 | A | 0.0803 | **2.33E-09** |
| IE1A | All subjects | 6 | **RS28986396** | 32 754 430 | A | 0.07998 | **2.47E-09** |
| IE1A | All subjects | 6 | **RS7762279** | 32 755 290 | G | -0.06748 | **2.00E-08** |
| IE1A | All subjects | 6 | **RS9276722** | 32 762 391 | G | 0.08029 | **1.98E-09** |
| IE1A | All subjects | 6 | RS9276726 | 32 763 888 | G | -0.0379 | 1.10E-06 |
| IE1A | All subjects | 6 | RS1383261 | 32 765 451 | A | -0.03421 | 9.71E-06 |
| IE1A | All subjects | 6 | RS6932969 | 32 765 671 | G | -0.03485 | 6.41E-06 |
| IE1A | All subjects | 6 | RS7383606 | 32 766 593 | A | -0.03468 | 7.03E-06 |
| IE1A | All subjects | 6 | RS9276734 | 32 766 854 | A | -0.03473 | 6.80E-06 |
| IE1A | All subjects | 6 | RS7382714 | 32 767 496 | G | -0.03473 | 6.92E-06 |
| IE1A | All subjects | 6 | RS4947350 | 32 767 620 | G | -0.0465 | 1.07E-07 |
| IE1A | All subjects | 6 | RS7381376 | 32 767 673 | G | -0.03525 | 5.21E-06 |

| | | | | | | | |
|---|---|---|---|---|---|---|---|
| IE1A | All subjects | 6 | RS2621367 | 32 768 721 | A | 0.03929 | 6.70E-07 |
| IE1A | All subjects | 6 | RS2199874 | 32 769 926 | G | 0.04007 | 3.52E-07 |
| IE1A | All subjects | 6 | RS6899857 | 32 770 482 | G | -0.03529 | 4.73E-06 |
| IE1A | All subjects | 6 | RS2621358 | 32 770 808 | A | 0.03781 | 1.91E-06 |
| IE1A | All subjects | 6 | RS6912414 | 32 774 465 | A | -0.03771 | 1.19E-06 |
| IE1A | All subjects | 6 | RS7382619 | 32 775 809 | A | -0.03434 | 8.59E-06 |
| IE1A | All subjects | 6 | RS7382649 | 32 776 087 | A | -0.03463 | 7.26E-06 |
| IE1A | All subjects | 6 | RS2621338 | 32 776 583 | A | 0.03788 | 1.82E-06 |
| IE1A | All subjects | 6 | RS2857118 | 32 777 900 | A | 0.03787 | 2.04E-06 |
| IE1A | All subjects | 6 | RS2857115 | 32 778 281 | A | 0.03964 | 9.59E-07 |
| IE1A | All subjects | 6 | **RS11244** | 32 780 724 | A | -0.04657 | **2.30E-08** |
| IE1A | All subjects | 6 | RS2071474 | 32 782 582 | A | 0.04235 | 1.74E-07 |
| IE1A | All subjects | 6 | RS2071472 | 32 784 620 | A | 0.04238 | 1.77E-07 |
| IE1A | All subjects | 6 | RS2071469 | 32 784 783 | A | 0.03576 | 2.17E-06 |
| IE1A | All subjects | 6 | RS1894406 | 32 787 056 | A | 0.03687 | 1.89E-06 |
| IE1A | All subjects | 6 | RS2621321 | 32 789 480 | G | 0.04648 | 7.14E-08 |
| IE1A | All subjects | 6 | **RS13209654** | 32 792 659 | G | 0.08061 | **1.85E-09** |
| IE1A | All subjects | 6 | RS2857101 | 32 794 676 | G | 0.04603 | 9.78E-08 |
| IE1A | All subjects | 6 | RS241456 | 32 795 965 | A | 0.04672 | 6.01E-08 |
| IE1A | All subjects | 6 | **RS241454** | 32 796 144 | G | 0.04715 | **4.45E-08** |
| IE1A | All subjects | 6 | RS241453 | 32 796 226 | A | 0.04659 | 6.37E-08 |
| IE1A | All subjects | 6 | RS241448 | 32 796 685 | G | 0.04623 | 8.30E-08 |
| IE1A | All subjects | 6 | **RS241447** | 32 796 751 | G | 0.04699 | **4.92E-08** |
| IE1A | All subjects | 6 | RS241446 | 32 796 967 | A | 0.04668 | 6.35E-08 |
| IE1A | All subjects | 6 | **RS241445** | 32 797 072 | A | 0.04709 | **4.61E-08** |
| IE1A | All subjects | 6 | RS241441 | 32 797 297 | G | 0.04664 | 6.31E-08 |
| IE1A | All subjects | 6 | RS241440 | 32 797 361 | A | 0.04614 | 8.94E-08 |
| IE1A | All subjects | 6 | **RS3819714** | 32 804 217 | A | 0.04267 | **4.91E-08** |
| IE1A | All subjects | 6 | RS3819715 | 32 804 219 | A | 0.0423 | 6.32E-08 |
| IE1A | All subjects | 6 | **RS3819720** | 32 804 570 | A | -0.05035 | **1.27E-10** |
| IE1A | All subjects | 6 | **RS241425** | 32 804 909 | A | 0.04212 | **9.94E-09** |
| IE1A | All subjects | 6 | RS241424 | 32 804 934 | A | -0.03762 | 3.47E-07 |
| IE1A | All subjects | 6 | RS2239701 | 32 805 049 | G | -0.03998 | 7.53E-08 |
| IE1A | All subjects | 6 | RS1480380 | 32 913 246 | A | -0.06895 | 1.60E-07 |
| IE1A | All subjects | 6 | RS9276933 | 32 930 795 | G | -0.0659 | 1.90E-07 |
| IE1A | All subjects | 6 | **RS423196** | 33 003 562 | G | -0.06196 | **2.76E-09** |
| IE1A | All subjects | 6 | **RS406477** | 33 005 644 | G | -0.06265 | **1.47E-09** |
| IE1A | All subjects | 6 | RS376877 | 33 024 606 | C | -0.04424 | 6.42E-06 |
| IE1A | All subjects | 6 | RS2395309 | 33 026 246 | G | -0.05448 | 2.29E-07 |
| IE1A | All subjects | 6 | RS17214519 | 33 032 188 | A | -0.05521 | 1.68E-07 |
| IE1A | All subjects | 6 | RS7905 | 33 032 975 | G | -0.05866 | 3.10E-07 |

| | | | | | | | |
|---|---|---|---|---|---|---|---|
| IE1A | All subjects | 6 | RS3077 | 33 033 022 | G | -0.05424 | 2.58E-07 |
| IE1A | All subjects | 6 | RS9469341 | 33 035 877 | G | -0.05555 | 1.36E-07 |
| IE1A | All subjects | 6 | RS1042190 | 33 036 999 | G | -0.05423 | 2.61E-07 |
| IE1A | All subjects | 6 | RS10214910 | 33 037 675 | A | -0.05442 | 2.49E-07 |
| IE1A | All subjects | 6 | RS2301224 | 33 038 369 | A | -0.05416 | 2.67E-07 |
| IE1A | All subjects | 6 | RS2301220 | 33 038 766 | A | -0.05542 | 1.39E-07 |
| IE1A | All subjects | 6 | RS6914849 | 33 040 715 | A | -0.0552 | 1.55E-07 |
| IE1A | All subjects | 6 | RS1431399 | 33 041 034 | G | -0.05453 | 2.18E-07 |
| IE1A | All subjects | 6 | RS1431400 | 33 041 176 | A | -0.05487 | 1.81E-07 |
| IE1A | All subjects | 6 | RS1431401 | 33 041 186 | A | -0.05484 | 1.85E-07 |
| IE1A | All subjects | 6 | **RS987870** | 33 042 880 | G | -0.07253 | **1.82E-09** |
| IE1A | All subjects | 6 | RS2071351 | 33 043 930 | G | -0.05449 | 2.30E-07 |
| IE1A | All subjects | 6 | **RS2071353** | 33 044 257 | G | -0.06394 | **3.28E-09** |
| IE1A | All subjects | 6 | **RS2071354** | 33 044 388 | G | -0.07234 | **2.02E-09** |
| IE1A | All subjects | 6 | **RS1042151** | 33 048 661 | G | -0.05617 | **1.15E-09** |
| IE1A | All subjects | 6 | **RS9277378** | 33 050 279 | G | -0.05042 | **4.77E-10** |
| IE1A | All subjects | 6 | **RS9277386** | 33 050 499 | G | -0.05196 | **1.41E-10** |
| IE1A | All subjects | 6 | **RS9277396** | 33 051 139 | A | -0.04625 | **8.54E-09** |
| IE1A | All subjects | 6 | **RS9277426** | 33 051 910 | A | -0.04599 | **1.02E-08** |
| IE1A | All subjects | 6 | **RS9277469** | 33 053 468 | A | -0.04682 | **5.53E-09** |
| IE1A | All subjects | 6 | **RS9277471** | 33 053 682 | A | -0.04583 | **1.20E-08** |
| IE1A | All subjects | 6 | **RS9277533** | 33 054 721 | A | -0.04582 | **1.15E-08** |
| IE1A | All subjects | 6 | **RS9277546** | 33 055 346 | C | -0.04616 | **9.20E-09** |
| IE1A | All subjects | 6 | **RS9277554** | 33 055 538 | A | -0.04606 | **9.61E-09** |
| IE1A | All subjects | 6 | RS9277555 | 33 055 605 | A | -0.04389 | 8.14E-08 |
| IE1A | All subjects | 6 | **RS3117228** | 33 056 435 | A | -0.04612 | **9.17E-09** |
| IE1A | All subjects | 6 | **RS3130188** | 33 057 176 | G | -0.04611 | **9.38E-09** |
| IE1A | All subjects | 6 | **RS3091282** | 33 057 198 | C | -0.04649 | **7.13E-09** |
| IE1A | All subjects | 6 | **RS3117226** | 33 057 659 | A | -0.0487 | **3.75E-09** |
| IE1A | All subjects | 6 | **RS3117225** | 33 057 711 | A | -0.0464 | **8.40E-09** |
| IE1A | All subjects | 6 | **RS3097652** | 33 057 835 | A | -0.04595 | **1.09E-08** |
| IE1A | All subjects | 6 | RS1367730 | 33 058 114 | A | -0.04427 | 6.53E-08 |
| IE1A | All subjects | 6 | RS3128972 | 33 058 774 | G | -0.04352 | 1.05E-07 |
| IE1A | All subjects | 6 | **RS2179920** | 33 058 874 | A | -0.0487 | **2.02E-08** |
| IE1A | All subjects | 6 | RS2179919 | 33 059 262 | G | -0.04412 | 6.92E-08 |
| IE1A | All subjects | 6 | RS3128917 | 33 059 996 | C | -0.04435 | 5.91E-08 |
| IE1A | All subjects | 6 | RS3117222 | 33 060 949 | A | -0.04411 | 7.10E-08 |
| IE1A | All subjects | 6 | RS3130190 | 33 061 690 | G | -0.04365 | 9.67E-08 |
| IE1A | All subjects | 6 | RS3130191 | 33 061 871 | G | -0.04352 | 1.06E-07 |
| IE1A | All subjects | 6 | **RS6914616** | 33 063 592 | A | -0.1102 | **1.68E-11** |
| IE1A | All subjects | 6 | RS3130198 | 33 063 931 | G | -0.04453 | 5.03E-08 |

| | | | | | | | |
|---|---|---|---|---|---|---|---|
| IE1A | All subjects | 6 | **RS3117213** | 33 064 605 | A | -0.04466 | **4.75E-08** |
| IE1A | All subjects | 6 | **RS2179915** | 33 065 734 | A | -0.1104 | **1.58E-11** |
| IE1A | All subjects | 6 | **RS2395319** | 33 067 211 | A | -0.04497 | **3.88E-08** |
| IE1A | All subjects | 6 | **RS3117239** | 33 071 777 | G | -0.04914 | **1.38E-08** |
| IE1A | All subjects | 6 | RS2064479 | 33 072 240 | A | -0.04345 | 1.33E-07 |
| IE1A | All subjects | 6 | **RS2064478** | 33 072 266 | A | -0.04933 | **1.23E-08** |
| IE1A | All subjects | 6 | **RS2064476** | 33 073 322 | G | -0.04644 | **7.16E-09** |
| IE1A | All subjects | 6 | **RS3117234** | 33 073 984 | G | -0.05084 | **3.19E-09** |
| IE1A | All subjects | 6 | **RS3117231** | 33 074 908 | G | -0.0492 | **2.37E-09** |
| IE1A | All subjects | 6 | **RS3117230** | 33 075 635 | G | -0.04914 | **1.38E-08** |
| IE1A | All subjects | 6 | **RS3128930** | 33 075 666 | A | -0.05013 | **2.99E-09** |
| IE1A | All subjects | 6 | **RS6924545** | 33 077 607 | A | -0.1072 | **5.64E-11** |
| IE1A | All subjects | 6 | RS6457713 | 33 077 776 | A | -0.06478 | 2.11E-07 |
| IE1A | All subjects | 6 | RS12529876 | 167 461 501 | A | 0.03555 | 1.61E-06 |
| IE1A | All subjects | 6 | RS6905911 | 167 521 624 | A | 0.03366 | 4.54E-06 |
| IE1A | All subjects | 6 | RS6907666 | 167 523 395 | A | 0.03363 | 6.16E-06 |
| IE1A | All subjects | 7 | RS6461830 | 24 969 081 | A | 0.05543 | 9.77E-06 |
| IE1A | All subjects | 14 | RS2582516 | 105 314 093 | C | 0.04642 | 1.24E-06 |
| IE1A | All subjects | 14 | RS7157970 | 105 463 936 | A | 0.03492 | 8.32E-06 |
| IE1A | All subjects | 21 | RS460014 | 16 091 441 | A | 0.03771 | 4.10E-06 |
| IE1A | All subjects | 22 | RS2530682 | 30 103 476 | A | -0.0342 | 8.89E-06 |
| IE1B | MS cases | 6 | RS3016014 | 31 351 153 | G | -0.05517 | 7.38E-07 |
| IE1B | MS cases | 6 | RS2256175 | 31 380 449 | A | 0.05208 | 1.19E-06 |
| IE1B | MS cases | 6 | RS10948679 | 52 012 873 | G | -0.04987 | 1.87E-06 |
| IE1B | MS cases | 7 | RS2598404 | 33 611 480 | C | 0.0523 | 4.91E-06 |
| IE1B | MS cases | 8 | RS1829402 | 134 574 971 | A | 0.08937 | 8.01E-06 |
| IE1B | MS cases | 12 | RS7958938 | 76 287 239 | A | -0.05894 | 8.92E-06 |
| IE1B | MS cases | 13 | RS1606713 | 61 581 129 | A | -0.06691 | 3.33E-06 |
| IE1B | MS cases | 13 | RS2323081 | 61 585 068 | A | -0.06698 | 3.64E-06 |
| IE1B | Controls | 2 | RS1374109 | 137 312 484 | A | 0.1229 | 4.09E-06 |
| IE1B | Controls | 5 | RS10079074 | 111 102 229 | A | 0.05946 | 8.68E-06 |
| IE1B | Controls | 6 | RS479536 | 32 193 678 | A | 0.1664 | 5.27E-07 |
| IE1B | Controls | 6 | RS374205 | 32 196 873 | G | 0.1261 | 5.90E-06 |
| IE1B | Controls | 6 | RS511027 | 32 206 687 | A | 0.09395 | 4.38E-06 |
| IE1B | Controls | 6 | RS28366191 | 32 364 190 | G | 0.1588 | 2.15E-06 |
| IE1B | Controls | 6 | RS6911777 | 32 409 996 | G | 0.1727 | 1.57E-07 |
| IE1B | Controls | 6 | RS13204672 | 32 582 796 | G | 0.09352 | 5.24E-06 |
| IE1B | Controls | 6 | RS2858308 | 32 670 000 | A | 0.1191 | 3.50E-06 |
| IE1B | Controls | 6 | RS16898262 | 32 677 397 | G | 0.1198 | 3.14E-06 |

| | | | | | | | |
|---|---|---|---|---|---|---|---|
| IE1B | Controls | 6 | RS2647089 | 32 681 568 | G | 0.06964 | 7.52E-06 |
| IE1B | Controls | 8 | RS12056386 | 26 050 530 | G | -0.07173 | 7.70E-06 |
| IE1B | Controls | 11 | RS10832934 | 18 435 700 | A | 0.07591 | 7.10E-06 |
| IE1B | All subjects | 2 | RS11896138 | 231 199 033 | A | 0.05443 | 7.84E-06 |
| IE1B | All subjects | 4 | RS7667582 | 36 154 555 | A | 0.03636 | 5.18E-06 |
| IE1B | All subjects | 6 | RS2394888 | 31 202 246 | A | -0.03805 | 7.05E-06 |
| IE1B | All subjects | 6 | RS3016014 | 31 351 153 | G | -0.04297 | 3.79E-07 |
| IE1B | All subjects | 6 | RS2256028 | 31 379 198 | A | 0.05436 | 7.24E-06 |
| IE1B | All subjects | 6 | RS2516463 | 31 416 536 | G | 0.09309 | 4.51E-06 |
| IE1B | All subjects | 6 | RS2596460 | 31 417 510 | G | 0.09267 | 4.79E-06 |
| IE1B | All subjects | 6 | RS2596458 | 31 417 647 | A | 0.09257 | 4.82E-06 |
| IE1B | All subjects | 6 | RS2516454 | 31 420 863 | T | 0.09579 | 2.44E-06 |
| IE1B | All subjects | 6 | RS2395028 | 31 424 757 | A | 0.09285 | 4.79E-06 |
| IE1B | All subjects | 6 | RS2596480 | 31 425 985 | A | 0.09373 | 3.77E-06 |
| IE1B | All subjects | 6 | RS2534678 | 31 463 963 | A | 0.07393 | 1.12E-07 |
| IE1B | All subjects | 6 | RS2534667 | 31 468 099 | A | 0.07811 | 1.52E-07 |
| IE1B | All subjects | 6 | RS2534657 | 31 472 459 | A | 0.07356 | 1.45E-07 |
| IE1B | All subjects | 6 | RS2844496 | 31 480 398 | A | 0.08718 | 7.98E-08 |
| IE1B | All subjects | 6 | **RS2246986** | 31 482 203 | G | 0.08893 | **4.40E-08** |
| IE1B | All subjects | 6 | RS2844509 | 31 510 924 | G | 0.05425 | 6.26E-07 |
| IE1B | All subjects | 6 | RS2239705 | 31 513 402 | A | 0.05972 | 1.38E-06 |
| IE1B | All subjects | 6 | RS693797 | 32 204 433 | G | 0.03651 | 9.46E-06 |
| IE1B | All subjects | 6 | **RS511027** | 32 206 687 | A | 0.08366 | **1.13E-08** |
| IE1B | All subjects | 6 | RS6910071 | 32 282 854 | G | -0.04378 | 7.88E-06 |
| IE1B | All subjects | 6 | RS9268557 | 32 389 305 | G | -0.03555 | 8.62E-06 |
| IE1B | All subjects | 7 | RS2598404 | 33 611 480 | C | 0.03855 | 7.35E-06 |
| IE1B | All subjects | 10 | RS1334461 | 99 868 267 | G | -0.03609 | 7.29E-06 |
| IE1B | All subjects | 10 | RS1334465 | 99 876 121 | G | -0.03667 | 5.21E-06 |
| IE1B | All subjects | 10 | RS7911699 | 99 890 470 | G | -0.03655 | 5.63E-06 |
| IE1B | All subjects | 10 | RS7088359 | 99 891 010 | G | -0.03598 | 7.56E-06 |
| IE1B | All subjects | 10 | RS10748715 | 99 914 261 | G | -0.03614 | 8.17E-06 |
| IE1B | All subjects | 10 | RS10450402 | 99 990 493 | G | -0.0376 | 3.41E-06 |
| IE1B | All subjects | 10 | RS3793692 | 100 008 436 | A | -0.03668 | 5.94E-06 |
| IE1B | All subjects | 12 | RS7958938 | 76 287 239 | A | -0.04589 | 3.82E-06 |
| IE1B | All subjects | 13 | RS9576177 | 37 679 729 | A | -0.03461 | 9.33E-06 |
| IE1B | All subjects | 13 | RS11147624 | 37 679 937 | A | -0.03463 | 8.36E-06 |
| IE1B | All subjects | 13 | RS11147625 | 37 680 081 | A | -0.03462 | 8.38E-06 |
| 101K | MS cases | 1 | RS12091891 | 9 000 220 | G | -0.08138 | 7.38E-06 |
| 101K | MS cases | 3 | RS12495466 | 13 282 611 | G | 0.1228 | 1.54E-06 |

| | | | | | | | |
|---|---|---|---|---|---|---|---|
| 101K | MS cases | 6 | RS9275224 | 32 659 878 | G | 0.04213 | 4.55E-06 |
| 101K | MS cases | 6 | RS9275245 | 32 660 943 | G | 0.04119 | 7.95E-06 |
| 101K | MS cases | 6 | RS6457617 | 32 663 851 | A | 0.04363 | 2.00E-06 |
| 101K | MS cases | 6 | RS9275371 | 32 668 296 | G | 0.04765 | 2.26E-06 |
| 101K | MS cases | 6 | RS9275374 | 32 668 526 | A | 0.04759 | 2.26E-06 |
| 101K | MS cases | 6 | RS9275388 | 32 669 084 | G | 0.04726 | 2.69E-06 |
| 101K | MS cases | 6 | RS9275390 | 32 669 156 | G | 0.04814 | 1.81E-06 |
| 101K | MS cases | 6 | RS9275393 | 32 669 439 | A | 0.0473 | 2.85E-06 |
| 101K | MS cases | 6 | RS9275406 | 32 669 955 | A | 0.04729 | 2.70E-06 |
| 101K | MS cases | 6 | RS9275407 | 32 670 037 | A | 0.04783 | 2.02E-06 |
| 101K | MS cases | 6 | RS9275418 | 32 670 244 | G | 0.0475 | 2.37E-06 |
| 101K | MS cases | 6 | RS9275424 | 32 670 576 | G | 0.04749 | 2.41E-06 |
| 101K | MS cases | 6 | RS9275428 | 32 670 978 | G | 0.04675 | 3.74E-06 |
| 101K | MS cases | 6 | RS9275439 | 32 671 521 | G | 0.04799 | 2.06E-06 |
| 101K | MS cases | 6 | RS9275523 | 32 674 994 | A | 0.04874 | 6.97E-06 |
| 101K | MS cases | 6 | RS9275582 | 32 680 070 | A | 0.0506 | 3.17E-06 |
| 101K | MS cases | 6 | RS9275595 | 32 681 355 | G | 0.04939 | 5.48E-06 |
| 101K | MS cases | 6 | RS17335164 | 44 662 629 | G | 0.08871 | 2.56E-06 |
| 101K | MS cases | 6 | RS7762246 | 51 099 256 | G | 0.05713 | 7.15E-06 |
| 101K | MS cases | 6 | RS12210804 | 104 100 800 | A | 0.139 | 7.47E-06 |
| 101K | MS cases | 7 | RS6947751 | 18 025 423 | G | -0.0418 | 2.16E-06 |
| 101K | MS cases | 7 | RS6947763 | 18 025 440 | G | -0.0422 | 1.73E-06 |
| 101K | MS cases | 9 | RS2821545 | 15 564 904 | A | -0.03938 | 7.51E-06 |
| 101K | MS cases | 9 | RS192466 | 15 585 395 | G | -0.04013 | 4.80E-06 |
| 101K | MS cases | 9 | RS1355171 | 15 681 694 | A | -0.03907 | 8.99E-06 |
| 101K | MS cases | 9 | RS10733295 | 15 681 901 | G | -0.03898 | 9.05E-06 |
| 101K | MS cases | 16 | RS7194313 | 50 059 500 | G | 0.07224 | 1.50E-06 |
| 101K | MS cases | 16 | RS17213264 | 50 060 203 | G | 0.06841 | 5.00E-06 |
| 101K | MS cases | 16 | RS6500279 | 50 073 908 | G | 0.07042 | 6.74E-06 |
| 101K | MS cases | 22 | RS4823817 | 46 857 705 | G | 0.04215 | 6.70E-06 |
| 101K | Controls | 3 | RS7616406 | 12 862 257 | G | 0.04442 | 3.92E-06 |
| 101K | Controls | 3 | RS6442333 | 12 864 386 | A | 0.04729 | 1.31E-06 |
| 101K | Controls | 3 | RS7629133 | 12 866 725 | G | 0.04571 | 5.96E-06 |
| 101K | Controls | 6 | RS2844571 | 31 335 647 | G | -0.04392 | 8.05E-06 |
| 101K | Controls | 6 | RS2253908 | 31 336 891 | A | -0.0442 | 7.12E-06 |
| 101K | Controls | 6 | RS2596530 | 31 387 373 | G | -0.04317 | 3.38E-06 |
| 101K | Controls | 6 | RS2596531 | 31 387 557 | G | -0.04356 | 2.75E-06 |
| 101K | Controls | 6 | RS2844511 | 31 389 784 | A | -0.04345 | 2.89E-06 |
| 101K | Controls | 6 | RS2516448 | 31 390 410 | A | -0.04346 | 2.89E-06 |
| 101K | Controls | 6 | RS2269475 | 31 583 931 | A | 0.05551 | 9.37E-06 |

| | | | | | | | |
|---|---|---|---|---|---|---|---|
| 101K | Controls | 6 | RS3763295 | 31 587 938 | G | 0.05879 | 3.06E-06 |
| 101K | Controls | 6 | RS14365 | 31 635 710 | G | 0.05665 | 2.09E-06 |
| 101K | Controls | 6 | RS28421666 | 32 592 737 | G | 0.09872 | 6.31E-06 |
| 101K | Controls | 6 | RS7774434 | 32 657 578 | G | 0.04619 | 1.27E-06 |
| 101K | Controls | 6 | RS6931062 | 32 835 615 | A | -0.0437 | 7.19E-06 |
| 101K | Controls | 6 | RS1580345 | 32 837 696 | C | -0.04417 | 5.64E-06 |
| 101K | Controls | 6 | RS6906376 | 32 838 584 | A | -0.0441 | 5.83E-06 |
| 101K | Controls | 6 | RS6929078 | 32 838 664 | G | -0.04416 | 5.68E-06 |
| 101K | Controls | 6 | RS7453903 | 32 838 949 | G | -0.04345 | 7.98E-06 |
| 101K | Controls | 6 | RS9276845 | 32 840 139 | G | -0.04472 | 4.54E-06 |
| 101K | Controls | 6 | RS9276864 | 32 843 238 | A | -0.0447 | 4.74E-06 |
| 101K | Controls | 6 | RS7757767 | 32 845 873 | G | -0.04427 | 5.31E-06 |
| 101K | Controls | 6 | RS2621426 | 32 846 570 | A | -0.0434 | 8.40E-06 |
| 101K | Controls | 6 | RS9276899 | 32 848 168 | A | -0.04387 | 6.76E-06 |
| 101K | Controls | 6 | RS2170185 | 32 849 639 | G | -0.04381 | 6.68E-06 |
| 101K | Controls | 6 | RS2127675 | 32 850 850 | G | -0.04365 | 7.40E-06 |
| 101K | Controls | 6 | RS4947259 | 32 851 507 | A | -0.04391 | 6.37E-06 |
| 101K | Controls | 9 | RS9406736 | 17 793 903 | A | 0.04771 | 7.00E-06 |
| 101K | Controls | 13 | RS11148543 | 61 707 375 | A | 0.04839 | 5.90E-06 |
| 101K | Controls | 13 | RS9317162 | 61 737 576 | A | 0.04826 | 6.77E-06 |
| 101K | Controls | 13 | RS9539055 | 61 755 041 | A | 0.05113 | 2.17E-06 |
| 101K | Controls | 13 | RS9539057 | 61 769 280 | A | 0.05091 | 2.28E-06 |
| 101K | Controls | 15 | RS10775256 | 92 997 155 | G | 0.04579 | 6.05E-06 |
| 101K | Controls | 20 | RS775121 | 16 046 227 | A | -0.05066 | 8.21E-06 |
| 101K | All subjects | 2 | RS1569193 | 51 720 700 | A | -0.03719 | 5.29E-06 |
| 101K | All subjects | 4 | RS10604 | 17 488 132 | G | 0.03208 | 2.47E-06 |
| 101K | All subjects | 4 | RS2597775 | 17 503 382 | A | 0.03127 | 4.52E-06 |
| 101K | All subjects | 6 | RS2735048 | 29 733 701 | A | -0.03252 | 1.47E-06 |
| 101K | All subjects | 6 | RS2735046 | 29 734 098 | A | -0.03087 | 2.89E-06 |
| 101K | All subjects | 6 | RS2523409 | 29 775 662 | G | -0.03267 | 9.81E-07 |
| 101K | All subjects | 6 | RS3115627 | 29 820 278 | G | -0.03098 | 5.13E-06 |
| 101K | All subjects | 6 | RS1611704 | 29 828 467 | A | -0.03167 | 3.31E-06 |
| 101K | All subjects | 6 | RS2523822 | 29 828 660 | G | -0.03151 | 5.16E-06 |
| 101K | All subjects | 6 | RS2860580 | 29 906 691 | A | 0.0283 | 8.79E-06 |
| 101K | All subjects | 6 | RS4713270 | 29 934 697 | A | -0.0306 | 8.97E-06 |
| 101K | All subjects | 6 | RS4713275 | 29 937 580 | C | -0.03082 | 8.81E-06 |
| 101K | All subjects | 6 | RS2256543 | 29 937 833 | A | 0.02804 | 9.87E-06 |
| 101K | All subjects | 6 | RS6916422 | 29 938 110 | A | -0.03066 | 8.61E-06 |
| 101K | All subjects | 6 | RS3823355 | 29 942 083 | A | -0.03073 | 8.45E-06 |
| 101K | All subjects | 6 | RS3823358 | 29 942 205 | A | -0.03082 | 8.62E-06 |

| | | | | | | | |
|---|---|---|---|---|---|---|---|
| 101K | All subjects | 6 | RS6904029 | 29 943 067 | A | -0.0308 | 8.08E-06 |
| 101K | All subjects | 6 | RS4713281 | 29 978 352 | A | -0.03129 | 6.52E-06 |
| 101K | All subjects | 6 | RS9357092 | 29 984 252 | A | -0.03168 | 5.00E-06 |
| 101K | All subjects | 6 | RS4711206 | 29 987 184 | A | -0.03083 | 8.83E-06 |
| 101K | All subjects | 6 | RS3910312 | 30 008 746 | C | -0.03159 | 5.27E-06 |
| 101K | All subjects | 6 | RS9380150 | 30 010 492 | G | -0.03114 | 6.79E-06 |
| 101K | All subjects | 6 | RS9295829 | 30 028 800 | G | -0.03129 | 6.34E-06 |
| 101K | All subjects | 6 | RS3807031 | 30 033 884 | A | -0.03362 | 7.12E-06 |
| 101K | All subjects | 6 | RS9393989 | 30 040 084 | A | -0.03135 | 6.22E-06 |
| 101K | All subjects | 6 | RS4711209 | 30 047 403 | A | -0.03116 | 7.08E-06 |
| 101K | All subjects | 6 | RS4959041 | 30 077 967 | G | -0.03281 | 1.90E-06 |
| 101K | All subjects | 6 | RS9261438 | 30 089 280 | A | -0.03135 | 4.76E-06 |
| 101K | All subjects | 6 | RS9261639 | 30 237 859 | A | -0.0318 | 7.16E-06 |
| 101K | All subjects | 6 | RS4713305 | 30 252 836 | A | -0.03165 | 7.35E-06 |
| 101K | All subjects | 6 | RS2844571 | 31 335 647 | G | -0.03392 | 7.19E-07 |
| 101K | All subjects | 6 | RS2253908 | 31 336 891 | A | -0.03409 | 6.39E-07 |
| 101K | All subjects | 6 | RS9295986 | 31 338 528 | A | 0.03241 | 8.86E-06 |
| 101K | All subjects | 6 | RS9266629 | 31 346 822 | G | -0.04199 | 8.10E-08 |
| 101K | All subjects | 6 | RS7741091 | 31 352 631 | G | 0.03793 | 1.03E-07 |
| 101K | All subjects | 6 | RS2523484 | 31 353 639 | A | 0.0336 | 8.90E-07 |
| 101K | All subjects | 6 | RS4711268 | 31 354 504 | A | 0.03823 | 1.14E-06 |
| 101K | All subjects | 6 | RS13437082 | 31 354 560 | A | 0.03898 | 6.92E-07 |
| 101K | All subjects | 6 | RS4711269 | 31 354 819 | A | 0.03892 | 7.15E-07 |
| 101K | All subjects | 6 | RS13437088 | 31 355 119 | A | 0.03885 | 7.51E-07 |
| 101K | All subjects | 6 | RS7751505 | 31 360 255 | C | 0.03748 | 1.99E-06 |
| 101K | All subjects | 6 | RS7751725 | 31 360 433 | G | 0.03826 | 1.24E-06 |
| 101K | All subjects | 6 | **RS2596530** | 31387373 | A | 0.03513 | **2.77E-08** |
| 101K | All subjects | 6 | **RS2596531** | 31387557 | A | 0.03552 | **1.87E-08** |
| 101K | All subjects | 6 | **RS2844511** | 31389784 | G | 0.03542 | **2.05E-08** |
| 101K | All subjects | 6 | **RS2516448** | 31390410 | G | 0.03558 | **1.77E-08** |
| 101K | All subjects | 6 | RS2523651 | 31 448 154 | A | -0.03294 | 3.86E-07 |
| 101K | All subjects | 6 | RS2523650 | 31 449 022 | A | -0.02988 | 8.51E-06 |
| 101K | All subjects | 6 | RS2269475 | 31 583 931 | A | 0.04192 | 2.02E-06 |
| 101K | All subjects | 6 | RS3763295 | 31 587 938 | G | 0.04406 | 6.74E-07 |
| 101K | All subjects | 6 | RS2242657 | 31 602 489 | G | 0.0406 | 3.82E-06 |
| 101K | All subjects | 6 | RS2295665 | 31 632 686 | A | 0.04194 | 1.78E-06 |
| 101K | All subjects | 6 | RS2295664 | 31 633 165 | A | 0.04167 | 2.06E-06 |
| 101K | All subjects | 6 | RS14365 | 31 635 710 | G | 0.04052 | 1.04E-06 |
| 101K | All subjects | 6 | RS2280800 | 31 646 398 | A | 0.04121 | 2.68E-06 |
| 101K | All subjects | 6 | RS2242653 | 31 675 765 | A | 0.04068 | 2.03E-06 |
| 101K | All subjects | 6 | RS2293861 | 31 711 124 | A | 0.04535 | 3.87E-07 |

| | | | | | | | |
|---|---|---|---|---|---|---|---|
| 101K | All subjects | 6 | RS2075788 | 31 712 181 | C | 0.04517 | 4.32E-07 |
| 101K | All subjects | 6 | RS3749953 | 31 713 124 | G | 0.04033 | 9.73E-06 |
| 101K | All subjects | 6 | RS3828922 | 31 713 454 | A | 0.04522 | 4.21E-07 |
| 101K | All subjects | 6 | RS6905572 | 31 731 881 | A | 0.0435 | 2.50E-06 |
| 101K | All subjects | 6 | RS35294112 | 31 794 443 | G | -0.0863 | 1.57E-06 |
| 101K | All subjects | 6 | RS17201248 | 31 803 130 | A | -0.0865 | 1.40E-06 |
| 101K | All subjects | 6 | RS397081 | 32 192 617 | G | -0.07245 | 1.21E-06 |
| 101K | All subjects | 6 | RS2294878 | 32 367 795 | A | 0.03214 | 8.22E-07 |
| 101K | All subjects | 6 | RS6926737 | 32 375 745 | A | 0.03213 | 6.59E-07 |
| 101K | All subjects | 6 | RS3763317 | 32 376 788 | A | 0.03332 | 3.26E-07 |
| 101K | All subjects | 6 | RS9268503 | 32 377 061 | A | 0.03161 | 1.02E-06 |
| 101K | All subjects | 6 | RS5007265 | 32 378 866 | C | 0.03267 | 4.29E-07 |
| 101K | All subjects | 6 | RS5007263 | 32 378 982 | G | 0.0322 | 6.35E-07 |
| 101K | All subjects | 6 | RS5007259 | 32 379 101 | G | 0.03208 | 6.94E-07 |
| 101K | All subjects | 6 | RS6932810 | 32 380 190 | A | 0.03152 | 1.10E-06 |
| 101K | All subjects | 6 | RS6932542 | 32 380 262 | G | 0.03225 | 6.07E-07 |
| 101K | All subjects | 6 | RS6937545 | 32 418 031 | A | -0.02979 | 4.45E-06 |
| 101K | All subjects | 6 | RS7763262 | 32 424 882 | A | -0.03026 | 3.31E-06 |
| 101K | All subjects | 6 | RS660895 | 32 577 380 | G | 0.04164 | 1.10E-07 |
| 101K | All subjects | 6 | RS4947342 | 32 653 070 | A | 0.03708 | 1.46E-06 |
| 101K | All subjects | 6 | RS2856692 | 32 653 385 | C | 0.03743 | 1.19E-06 |
| 101K | All subjects | 6 | **RS7774434** | 32657578 | G | 0.03808 | **1.48E-08** |
| 101K | All subjects | 6 | **RS9275224** | 32659878 | G | 0.03766 | **8.42E-09** |
| 101K | All subjects | 6 | RS2858324 | 32 660 375 | G | 0.0343 | 1.47E-07 |
| 101K | All subjects | 6 | **RS9275245** | 32660943 | G | 0.03877 | **3.47E-09** |
| 101K | All subjects | 6 | RS5000634 | 32 663 564 | G | 0.03433 | 7.96E-07 |
| 101K | All subjects | 6 | **RS6457617** | 32663851 | A | 0.03921 | **1.93E-09** |
| 101K | All subjects | 6 | RS6457622 | 32 664 163 | C | 0.03445 | 7.11E-07 |
| 101K | All subjects | 6 | RS2647012 | 32 664 458 | G | 0.03403 | 1.79E-07 |
| 101K | All subjects | 6 | RS2647003 | 32 664 880 | C | 0.03434 | 1.40E-07 |
| 101K | All subjects | 6 | **RS9275312** | 32665728 | G | 0.04905 | **4.92E-09** |
| 101K | All subjects | 6 | RS2856725 | 32 666 738 | A | 0.03393 | 1.95E-07 |
| 101K | All subjects | 6 | **RS9275328** | 32666822 | A | 0.04896 | **5.57E-09** |
| 101K | All subjects | 6 | **RS9275330** | 32666875 | G | 0.04862 | **7.10E-09** |
| 101K | All subjects | 6 | **RS9275333** | 32666968 | G | 0.04852 | **7.54E-09** |
| 101K | All subjects | 6 | **RS9275371** | 32668296 | G | 0.04466 | **3.64E-10** |
| 101K | All subjects | 6 | **RS9275374** | 32668526 | A | 0.04495 | **2.70E-10** |
| 101K | All subjects | 6 | RS1612904 | 32 669 018 | C | -0.03378 | 2.09E-07 |
| 101K | All subjects | 6 | **RS9275388** | 32669084 | G | 0.04475 | **3.26E-10** |
| 101K | All subjects | 6 | **RS9275390** | 32669156 | G | 0.04508 | **2.52E-10** |
| 101K | All subjects | 6 | **RS9275393** | 32669439 | A | 0.04478 | **3.53E-10** |

| | | | | | | | |
|---|---|---|---|---|---|---|---|
| 101K | All subjects | 6 | **RS9275406** | 32669955 | A | 0.04486 | **3.02E-10** |
| 101K | All subjects | 6 | **RS9275407** | 32670037 | A | 0.0451 | **2.38E-10** |
| 101K | All subjects | 6 | **RS9275418** | 32670244 | G | 0.04484 | **2.97E-10** |
| 101K | All subjects | 6 | RS2856717 | 32 670 308 | G | 0.03422 | 1.50E-07 |
| 101K | All subjects | 6 | RS2858305 | 32 670 464 | A | 0.03464 | 1.06E-07 |
| 101K | All subjects | 6 | **RS9275424** | 32670576 | G | 0.0448 | **3.15E-10** |
| 101K | All subjects | 6 | **RS9275428** | 32670978 | G | 0.04493 | **3.09E-10** |
| 101K | All subjects | 6 | **RS9275439** | 32671521 | G | 0.04487 | **3.21E-10** |
| 101K | All subjects | 6 | RS9275523 | 32 674 994 | A | 0.03628 | 1.62E-06 |
| 101K | All subjects | 6 | RS9275582 | 32 680 070 | A | 0.03789 | 6.16E-07 |
| 101K | All subjects | 6 | RS9275595 | 32 681 355 | G | 0.03688 | 1.22E-06 |
| 101K | All subjects | 6 | RS9275596 | 32 681 631 | G | -0.03509 | 7.08E-08 |
| 101K | All subjects | 6 | RS6931062 | 32 835 615 | A | -0.03068 | 5.12E-06 |
| 101K | All subjects | 6 | RS1580345 | 32 837 696 | C | -0.03091 | 4.26E-06 |
| 101K | All subjects | 6 | RS6906376 | 32 838 584 | A | -0.03078 | 4.68E-06 |
| 101K | All subjects | 6 | RS6929078 | 32 838 664 | G | -0.03106 | 3.82E-06 |
| 101K | All subjects | 6 | RS7453903 | 32 838 949 | G | -0.03059 | 5.37E-06 |
| 101K | All subjects | 6 | RS9276845 | 32 840 139 | G | -0.03121 | 3.65E-06 |
| 101K | All subjects | 6 | RS9276864 | 32 843 238 | A | -0.03066 | 5.57E-06 |
| 101K | All subjects | 6 | RS7757767 | 32 845 873 | G | -0.03112 | 3.63E-06 |
| 101K | All subjects | 6 | RS2621426 | 32 846 570 | A | -0.03045 | 6.13E-06 |
| 101K | All subjects | 6 | RS9276899 | 32 848 168 | A | -0.03079 | 4.81E-06 |
| 101K | All subjects | 6 | RS9276900 | 32 848 566 | C | -0.02999 | 9.61E-06 |
| 101K | All subjects | 6 | RS2170185 | 32 849 639 | G | -0.03081 | 4.60E-06 |
| 101K | All subjects | 6 | RS2127675 | 32 850 850 | G | -0.03075 | 4.89E-06 |
| 101K | All subjects | 6 | RS4947259 | 32 851 507 | A | -0.03088 | 4.34E-06 |
| 101K | All subjects | 19 | RS7254645 | 54 775 389 | G | 0.03379 | 2.57E-07 |
| 101K | All subjects | 19 | RS8100863 | 54 789 266 | C | 0.02862 | 8.04E-06 |
| 101K | All subjects | 19 | RS6509860 | 54 792 358 | A | 0.03142 | 2.23E-06 |

#### Table S8. Antibody responses analyzed for association with non-HLA SNPs serostatus

| Antigen | Subject group | Chromo-some | SNP marker | Basepair position | Minor allele | OR (L95-U95)* | p |
|---|---|---|---|---|---|---|---|
| IE1A | MS cases | 1 | RS12566754 | 166 444 011 | A | 0.7968 (0.7209-0.8808) | 8,83E-06 |
| IE1A | MS cases | 3 | RS17129314 | 197 031 608 | G | 0.5514 (0.4282-0.7101) | 3,99E-06 |
| IE1A | MS cases | 4 | RS17277682 | 161 479 466 | A | 0.7201 (0.6291-0.8243) | 1,91E-06 |
| IE1A | MS cases | 4 | RS11936687 | 163 288 053 | A | 1.31 (1.175-1.46) | 1,05E-06 |
| IE1A | MS cases | 6 | RS3130544 | 31 058 340 | A | 0.7114 (0.6139-0.8244) | 5,97E-06 |

| | | | | | | | |
|---|---|---|---|---|---|---|---|
| IE1A | MS cases | 6 | RS3130557 | 31 094 703 | A | 0.705 (0.6082-0.8172) | 3,48E-06 |
| IE1A | MS cases | 6 | RS7750641 | 31 129 310 | A | 0.7017 (0.6056-0.8131) | 2,46E-06 |
| IE1A | MS cases | 6 | RS3132510 | 31 172 151 | G | 0.6693 (0.5736-0.7811) | 3,49E-07 |
| IE1A | MS cases | 6 | RS2596487 | 31 325 056 | A | 0.6903 (0.6037-0.7894) | 6,13E-08 |
| IE1A | MS cases | 6 | **RS2523573** | 31 328 988 | C | 0.6381 (0.5495-0.7409) | **3,82E-09** |
| IE1A | MS cases | 6 | RS9266669 | 31 348 077 | A | 0.6781 (0.5882-0.7817) | 8,61E-08 |
| IE1A | MS cases | 6 | **RS3094013** | 31 434 366 | A | 0.6379 (0.5493-0.7408) | **3,76E-09** |
| IE1A | MS cases | 6 | **RS3131618** | 31 434 621 | G | 0.6413 (0.5517-0.7454) | **7,14E-09** |
| IE1A | MS cases | 6 | **RS3131643** | 31 442 782 | A | 0.6774 (0.5898-0.7781) | **3,63E-08** |
| IE1A | MS cases | 6 | **RS3099844** | 31 448 976 | A | 0.6261 (0.5393-0.7269) | **7,81E-10** |
| IE1A | MS cases | 6 | **RS3094011** | 31 451 836 | G | 0.6299 (0.5422-0.7317) | **1,49E-09** |
| IE1A | MS cases | 6 | **RS9267445** | 31 483 481 | C | 0.6372 (0.5476-0.7414) | **5,50E-09** |
| IE1A | MS cases | 6 | RS3093988 | 31 492 453 | A | 0.6958 (0.608-0.7964) | 1,38E-07 |
| IE1A | MS cases | 6 | RS2516482 | 31 496 569 | G | 0.7055 (0.6166-0.8071) | 3,75E-07 |
| IE1A | MS cases | 6 | RS2734583 | 31 505 480 | G | 0.6671 (0.5745-0.7747) | 1,12E-07 |
| IE1A | MS cases | 6 | RS1800629 | 31 543 031 | A | 0.7129 (0.6236-0.815) | 7,23E-07 |
| IE1A | MS cases | 6 | RS3117582 | 31 620 520 | C | 0.6607 (0.5668-0.7702) | 1,17E-07 |
| IE1A | MS cases | 6 | RS9267531 | 31 636 742 | G | 0.6607 (0.5668-0.7702) | 1,17E-07 |
| IE1A | MS cases | 6 | RS3101018 | 31 705 864 | A | 0.6578 (0.5635-0.7679) | 1,14E-07 |
| IE1A | MS cases | 6 | RS3132445 | 31 712 196 | A | 0.6635 (0.5691-0.7737) | 1,64E-07 |
| IE1A | MS cases | 6 | RS3130484 | 31 715 882 | G | 0.6619 (0.5677-0.7716) | 1,36E-07 |
| IE1A | MS cases | 6 | RS3131379 | 31 721 033 | A | 0.6643 (0.5697-0.7746) | 1,81E-07 |
| IE1A | MS cases | 6 | RS3117574 | 31 725 230 | A | 0.6649 (0.5703-0.7752) | 1,88E-07 |
| IE1A | MS cases | 6 | RS3131378 | 31 725 285 | G | 0.6612 (0.5671-0.7709) | 1,27E-07 |
| IE1A | MS cases | 6 | RS3117577 | 31 727 474 | G | 0.6611 (0.5671-0.7708) | 1,26E-07 |
| IE1A | MS cases | 6 | RS3101017 | 31 733 466 | G | 0.6653 (0.5707-0.7755) | 1,88E-07 |
| IE1A | MS cases | 6 | RS652888 | 31 851 234 | G | 0.755 (0.6673-0.8541) | 7,94E-06 |
| IE1A | MS cases | 6 | RS558702 | 31 870 326 | A | 0.6539 (0.5606-0.7628) | 6,46E-08 |
| IE1A | MS cases | 6 | **RS519417** | 31 878 433 | A | 0.6458 (0.5533-0.7538) | **2,98E-08** |
| IE1A | MS cases | 6 | **RS497309** | 31 892 484 | C | 0.6496 (0.5571-0.7575) | **3,72E-08** |
| IE1A | MS cases | 6 | **RS1270942** | 31 918 860 | G | 0.65 (0.5574-0.7578) | **3,82E-08** |
| IE1A | MS cases | 6 | RS1150755 | 32 038 550 | A | 0.7106 (0.6203-0.8141) | 8,42E-07 |
| IE1A | MS cases | 6 | RS7774197 | 32 046 275 | C | 1.728 (1.358-2.198) | 8,42E-06 |
| IE1A | MS cases | 6 | RS1150754 | 32 050 758 | A | 0.7087 (0.6186-0.8119) | 6,92E-07 |
| IE1A | MS cases | 6 | RS1150753 | 32 059 867 | G | 0.6557 (0.5621-0.7648) | 7,72E-08 |
| IE1A | MS cases | 6 | RS1150752 | 32 064 726 | G | 0.6557 (0.5617-0.7655) | 9,06E-08 |
| IE1A | MS cases | 6 | RS1269852 | 32 080 191 | C | 0.659 (0.5648-0.769) | 1,17E-07 |
| IE1A | MS cases | 6 | RS3130288 | 32 096 001 | A | 0.6606 (0.5663-0.7706) | 1,33E-07 |
| IE1A | MS cases | 6 | RS3134608 | 32 117 971 | C | 0.7446 (0.6537-0.8481) | 8,94E-06 |
| IE1A | MS cases | 6 | RS3096697 | 32 134 510 | A | 0.7413 (0.6507-0.8445) | 6,74E-06 |
| IE1A | MS cases | 6 | RS3130347 | 32 134 656 | G | 0.7438 (0.653-0.8472) | 8,37E-06 |

| | | | | | | | |
|---|---|---|---|---|---|---|---|
| IE1A | MS cases | 6 | RS3130284 | 32 140 487 | G | 0.7441 (0.6532-0.8477) | 8,77E-06 |
| IE1A | MS cases | 6 | RS3134947 | 32 145 205 | A | 0.744 (0.6531-0.8475) | 8,59E-06 |
| IE1A | MS cases | 6 | RS3134945 | 32 146 492 | A | 0.7439 (0.6529-0.8475) | 8,72E-06 |
| IE1A | MS cases | 6 | RS3130349 | 32 147 696 | A | 0.7167 (0.6263-0.82) | 1,26E-06 |
| IE1A | MS cases | 6 | RS1800625 | 32 152 442 | G | 0.7172 (0.6269-0.8206) | 1,32E-06 |
| IE1A | MS cases | 6 | RS204993 | 32 155 581 | G | 0.7449 (0.6591-0.8419) | 2,42E-06 |
| IE1A | MS cases | 6 | RS204991 | 32 161 366 | G | 0.7257 (0.6342-0.8304) | 3,14E-06 |
| IE1A | MS cases | 6 | RS204990 | 32 161 430 | A | 0.7195 (0.6279-0.8244) | 2,13E-06 |
| IE1A | MS cases | 6 | RS2071278 | 32 165 444 | G | 0.686 (0.5945-0.7916) | 2,45E-07 |
| IE1A | MS cases | 6 | **RS3134942** | 32 168 771 | A | 0.657 (0.5666-0.7619) | **2,68E-08** |
| IE1A | MS cases | 6 | RS2071277 | 32 171 683 | G | 0.7884 (0.7112-0.8741) | 6,24E-06 |
| IE1A | MS cases | 6 | **RS3131296** | 32 172 993 | A | 0.6535 (0.563-0.7585) | **2,20E-08** |
| IE1A | MS cases | 6 | **RS3132956** | 32 179 438 | A | 0.6583 (0.5677-0.7632) | **3,05E-08** |
| IE1A | MS cases | 6 | RS3131290 | 32 183 175 | A | 0.7554 (0.6789-0.8405) | 2,60E-07 |
| IE1A | MS cases | 6 | RS3134799 | 32 184 221 | A | 0.7765 (0.6995-0.862) | 2,05E-06 |
| IE1A | MS cases | 6 | RS436388 | 32 186 264 | G | 0.7947 (0.7176-0.88) | 9,98E-06 |
| IE1A | MS cases | 6 | RS394657 | 32 187 023 | G | 0.7797 (0.7025-0.8654) | 2,93E-06 |
| IE1A | MS cases | 6 | RS1109771 | 32 187 605 | G | 0.7939 (0.7169-0.8792) | 9,36E-06 |
| IE1A | MS cases | 6 | **RS915894** | 32 190 390 | C | 0.7146 (0.6406-0.797) | **1,62E-09** |
| IE1A | MS cases | 6 | RS3134930 | 32 191 620 | A | 0.7267 (0.641-0.8239) | 6,21E-07 |
| IE1A | MS cases | 6 | **RS6901158** | 32 205 942 | A | 0.6498 (0.563-0.7499) | **3,75E-09** |
| IE1A | MS cases | 6 | RS424232 | 32 208 324 | A | 0.7284 (0.6476-0.8192) | 1,26E-07 |
| IE1A | MS cases | 6 | **RS3132971** | 32 230 256 | C | 0.6315 (0.5409-0.7372) | **5,94E-09** |
| IE1A | MS cases | 6 | **RS7775397** | 32 261 252 | C | 0.6366 (0.5454-0.743) | **1,02E-08** |
| IE1A | MS cases | 6 | RS9268219 | 32 284 108 | C | 0.6475 (0.5525-0.7588) | 7,82E-08 |
| IE1A | MS cases | 6 | **RS9268235** | 32 290 208 | A | 0.6396 (0.5478-0.7468) | **1,57E-08** |
| IE1A | MS cases | 6 | RS1265760 | 32 321 872 | G | 0.797 (0.7207-0.8814) | 9,99E-06 |
| IE1A | MS cases | 6 | RS3129911 | 32 328 403 | A | 0.797 (0.7207-0.8814) | 9,99E-06 |
| IE1A | MS cases | 6 | RS3129922 | 32 333 098 | A | 0.7933 (0.7172-0.8775) | 6,77E-06 |
| IE1A | MS cases | 6 | RS2273017 | 32 337 630 | A | 0.7792 (0.7039-0.8624) | 1,46E-06 |
| IE1A | MS cases | 6 | **RS2073045** | 32 339 548 | A | 0.7202 (0.6463-0.8027) | **2,92E-09** |
| IE1A | MS cases | 6 | **RS2894254** | 32 345 689 | C | 0.6423 (0.5498-0.7504) | **2,42E-08** |
| IE1A | MS cases | 6 | RS3129843 | 32 395 726 | G | 0.6511 (0.5574-0.7606) | 6,28E-08 |
| IE1A | MS cases | 6 | RS3763326 | 32 413 557 | G | 2.179 (1.577-3.012) | 2,36E-06 |
| IE1A | MS cases | 6 | RS9270665 | 32 566 232 | A | 0.7426 (0.6527-0.8449) | 6,17E-06 |
| IE1A | MS cases | 6 | RS642093 | 32 582 075 | A | 0.7431 (0.6544-0.8438) | 4,69E-06 |
| IE1A | MS cases | 6 | RS3129763 | 32 590 925 | A | 0.7427 (0.6529-0.8449) | 6,14E-06 |
| IE1A | MS cases | 6 | RS9272219 | 32 602 269 | A | 0.7528 (0.6705-0.8451) | 1,50E-06 |
| IE1A | MS cases | 6 | RS2187668 | 32 605 884 | A | 0.7031 (0.607-0.8143) | 2,60E-06 |
| IE1A | MS cases | 6 | RS9273012 | 32 611 641 | G | 0.7504 (0.6685-0.8423) | 1,11E-06 |
| IE1A | MS cases | 6 | **RS17843604** | 32 620 283 | A | 0.7155 (0.6429-0.7963) | **8,66E-10** |

| | | | | | | | |
|---|---|---|---|---|---|---|---|
| IE1A | MS cases | 6 | RS9273327 | 32 623 223 | C | 0.7037 (0.6066-0.8163) | 3,52E-06 |
| IE1A | MS cases | 6 | **RS9273349** | 32 625 869 | G | 0.6862 (0.6161-0.7643) | **7,37E-12** |
| IE1A | MS cases | 6 | **RS9273363** | 32 626 272 | A | 0.7099 (0.6336-0.7955) | **3,60E-09** |
| IE1A | MS cases | 6 | **RS6906021** | 32 626 311 | G | 0.7379 (0.6617-0.8229) | **4,59E-08** |
| IE1A | MS cases | 6 | **RS1063355** | 32 627 714 | C | 0.6865 (0.6164-0.7647) | **7,91E-12** |
| IE1A | MS cases | 6 | RS9273448 | 32 627 747 | A | 1.335 (1.202-1.483) | 6,32E-08 |
| IE1A | MS cases | 6 | RS2854275 | 32 628 428 | A | 0.6999 (0.6043-0.8105) | 1,89E-06 |
| IE1A | MS cases | 6 | RS1794282 | 32 666 526 | A | 0.6545 (0.5601-0.7648) | 9,61E-08 |
| IE1A | MS cases | 6 | RS3135006 | 32 667 119 | A | 1.335 (1.202-1.484) | 7,37E-08 |
| IE1A | MS cases | 6 | RS4947344 | 32 677 846 | G | 0.7733 (0.6983-0.8564) | 8,04E-07 |
| IE1A | MS cases | 6 | RS9276689 | 32 751 962 | A | 0.6455 (0.5489-0.7591) | 1,22E-07 |
| IE1A | MS cases | 6 | RS7762279 | 32 755 290 | G | 0.6421 (0.5464-0.7545) | 7,31E-08 |
| IE1A | MS cases | 6 | RS4947350 | 32 767 620 | G | 0.742 (0.6573-0.8377) | 1,42E-06 |
| IE1A | MS cases | 6 | RS11244 | 32 780 724 | A | 0.7655 (0.6825-0.8586) | 5,02E-06 |
| IE1A | MS cases | 6 | **RS3819720** | 32 804 570 | A | 0.7304 (0.6548-0.8147) | **1,77E-08** |
| IE1A | MS cases | 6 | **RS241425** | 32 804 909 | G | 0.7515 (0.6797-0.8309) | **2,48E-08** |
| IE1A | MS cases | 6 | RS241424 | 32 804 934 | A | 0.7698 (0.6958-0.8516) | 3,84E-07 |
| IE1A | MS cases | 6 | RS2239701 | 32 805 049 | G | 0.7565 (0.683-0.8379) | 8,84E-08 |
| IE1A | MS cases | 6 | RS2071465 | 32 805 470 | C | 0.7743 (0.6987-0.8581) | 1,08E-06 |
| IE1A | MS cases | 6 | RS3763366 | 32 807 446 | C | 0.789 (0.7131-0.8729) | 4,32E-06 |
| IE1A | MS cases | 6 | RS3763349 | 32 808 232 | G | 0.7837 (0.7085-0.8669) | 2,20E-06 |
| IE1A | MS cases | 6 | RS9276810 | 32 810 443 | A | 0.7892 (0.7132-0.8734) | 4,70E-06 |
| IE1A | MS cases | 6 | RS406477 | 33 005 644 | G | 0.718 (0.6229-0.8275) | 4,81E-06 |
| IE1A | MS cases | 6 | RS987870 | 33 042 880 | G | 0.6726 (0.5686-0.7955) | 3,66E-06 |
| IE1A | MS cases | 6 | RS2071354 | 33 044 388 | G | 0.6733 (0.5692-0.7963) | 3,86E-06 |
| IE1A | MS cases | 6 | RS1042151 | 33 048 661 | G | 0.732 (0.6474-0.8276) | 6,39E-07 |
| IE1A | MS cases | 6 | RS9277378 | 33 050 279 | G | 0.7644 (0.6856-0.8523) | 1,31E-06 |
| IE1A | MS cases | 6 | RS9277386 | 33 050 499 | G | 0.7561 (0.678-0.8432) | 5,00E-07 |
| IE1A | MS cases | 6 | RS9277396 | 33 051 139 | A | 0.7774 (0.698-0.8658) | 4,65E-06 |
| IE1A | MS cases | 6 | RS9277426 | 33 051 910 | A | 0.7768 (0.6976-0.8651) | 4,23E-06 |
| IE1A | MS cases | 6 | RS9277469 | 33 053 468 | A | 0.7754 (0.6963-0.8636) | 3,66E-06 |
| IE1A | MS cases | 6 | RS9277471 | 33 053 682 | A | 0.7777 (0.6983-0.8661) | 4,70E-06 |
| IE1A | MS cases | 6 | RS9277533 | 33 054 721 | A | 0.7788 (0.6993-0.8674) | 5,36E-06 |
| IE1A | MS cases | 6 | RS9277546 | 33 055 346 | C | 0.7764 (0.6971-0.8647) | 4,11E-06 |
| IE1A | MS cases | 6 | RS9277554 | 33 055 538 | A | 0.7768 (0.6976-0.8651) | 4,23E-06 |
| IE1A | MS cases | 6 | RS3117228 | 33 056 435 | A | 0.7764 (0.6972-0.8647) | 4,05E-06 |
| IE1A | MS cases | 6 | RS3130188 | 33 057 176 | G | 0.7756 (0.6964-0.8637) | 3,68E-06 |
| IE1A | MS cases | 6 | RS3091282 | 33 057 198 | C | 0.7757 (0.6965-0.8639) | 3,80E-06 |
| IE1A | MS cases | 6 | RS3117226 | 33 057 659 | A | 0.7683 (0.6884-0.8576) | 2,58E-06 |
| IE1A | MS cases | 6 | RS3117225 | 33 057 711 | A | 0.7752 (0.6959-0.8635) | 3,71E-06 |
| IE1A | MS cases | 6 | RS3097652 | 33 057 835 | A | 0.7772 (0.6977-0.8658) | 4,70E-06 |

| | | | | | | | |
|---|---|---|---|---|---|---|---|
| IE1A | MS cases | 6 | RS2179920 | 33 058 874 | A | 0.7517 (0.6699-0.8435) | 1,20E-06 |
| IE1A | MS cases | 6 | RS6914616 | 33 063 592 | A | 0.5906 (0.4706-0.7412) | 5,50E-06 |
| IE1A | MS cases | 6 | RS2179915 | 33 065 734 | A | 0.591 (0.4709-0.7417) | 5,66E-06 |
| IE1A | MS cases | 6 | RS3117239 | 33 071 777 | G | 0.7571 (0.6748-0.8495) | 2,17E-06 |
| IE1A | MS cases | 6 | RS2064478 | 33 072 266 | A | 0.7572 (0.6748-0.8496) | 2,19E-06 |
| IE1A | MS cases | 6 | RS2064476 | 33 073 322 | G | 0.7769 (0.6977-0.8651) | 4,20E-06 |
| IE1A | MS cases | 6 | RS3117234 | 33 073 984 | G | 0.7479 (0.6674-0.838) | 5,67E-07 |
| IE1A | MS cases | 6 | RS3117231 | 33 074 908 | G | 0.7734 (0.6929-0.8632) | 4,57E-06 |
| IE1A | MS cases | 6 | RS3117230 | 33 075 635 | G | 0.7572 (0.6748-0.8496) | 2,19E-06 |
| IE1A | MS cases | 6 | RS3128930 | 33 075 666 | A | 0.746 (0.6663-0.8353) | 3,75E-07 |
| IE1A | MS cases | 10 | RS12255701 | 90 532 552 | A | 1.568 (1.292-1.902) | 5,22E-06 |
| IE1A | MS cases | 12 | RS706536 | 26 130 945 | G | 0.7792 (0.7026-0.8642) | 2,34E-06 |
| IE1A | MS cases | 22 | RS1061660 | 30 819 628 | G | 0.7739 (0.6931-0.8641) | 5,16E-06 |
| IE1A | Controls | 3 | RS7651713 | 111399209 | A | 1.382 (1.21-1.578) | 1,89E-06 |
| IE1A | Controls | 3 | RS1355767 | 111416310 | A | 1.342 (1.185-1.521) | 3,86E-06 |
| IE1A | Controls | 6 | RS1737031 | 29737481 | A | 0.7539 (0.6678-0.851) | 4,95E-06 |
| IE1A | Controls | 6 | RS1633030 | 29745794 | G | 0.7534 (0.6673-0.8505) | 4,68E-06 |
| IE1A | Controls | 6 | RS1633028 | 29746367 | C | 0.7555 (0.6692-0.853) | 5,94E-06 |
| IE1A | Controls | 6 | RS2735042 | 29747373 | G | 0.7547 (0.6686-0.852) | 5,36E-06 |
| IE1A | Controls | 6 | RS1610649 | 29768917 | G | 0.7582 (0.6743-0.8524) | 3,65E-06 |
| IE1A | Controls | 6 | RS1632983 | 29773509 | A | 0.7595 (0.6735-0.8564) | 7,17E-06 |
| IE1A | Controls | 6 | RS1610678 | 29789190 | G | 0.7423 (0.6642-0.8295) | 1,48E-07 |
| IE1A | Controls | 6 | RS1611149 | 29789999 | A | 0.7429 (0.6648-0.8301) | 1,53E-07 |
| IE1A | Controls | 6 | **RS1736936** | 29794317 | A | 0.7155 (0.6395-0.8005) | **5,09E-09** |
| IE1A | Controls | 6 | RS915669 | 29798425 | A | 0.7384 (0.6607-0.8253) | 9,04E-08 |
| IE1A | Controls | 6 | RS1611133 | 29809382 | A | 0.7241 (0.6329-0.8286) | 2,67E-06 |
| IE1A | Controls | 6 | RS1611750 | 29814778 | C | 0.7164 (0.6258-0.8201) | 1,34E-06 |
| IE1A | Controls | 6 | RS2247504 | 29819077 | A | 0.7688 (0.6867-0.8608) | 5,09E-06 |
| IE1A | Controls | 6 | RS2517854 | 29823296 | G | 0.7751 (0.6923-0.8678) | 9,85E-06 |
| IE1A | Controls | 6 | RS1611737 | 29831571 | G | 0.7713 (0.689-0.8634) | 6,41E-06 |
| IE1A | Controls | 6 | RS2844806 | 29933439 | A | 0.7791 (0.6978-0.8699) | 9,01E-06 |
| IE1A | Controls | 6 | RS1362104 | 30101656 | A | 0.7449 (0.6655-0.8337) | 3,02E-07 |
| IE1A | Controls | 6 | RS3130544 | 31058340 | A | 0.6651 (0.5605-0.7891) | 2,94E-06 |
| IE1A | Controls | 6 | RS3130557 | 31094703 | A | 0.6558 (0.5524-0.7785) | 1,44E-06 |
| IE1A | Controls | 6 | RS7750641 | 31129310 | A | 0.6638 (0.5594-0.7877) | 2,68E-06 |
| IE1A | Controls | 6 | RS3132510 | 31172151 | G | 0.6528 (0.5461-0.7805) | 2,87E-06 |
| IE1A | Controls | 6 | RS2523573 | 31328988 | C | 0.6521 (0.5482-0.7757) | 1,38E-06 |
| IE1A | Controls | 6 | RS9266669 | 31348077 | A | 0.6609 (0.5635-0.7751) | 3,53E-07 |
| IE1A | Controls | 6 | RS3094013 | 31434366 | A | 0.6559 (0.5517-0.7798) | 1,77E-06 |
| IE1A | Controls | 6 | RS3131618 | 31434621 | G | 0.647 (0.5436-0.7702) | 9,69E-07 |

| | | | | | | | |
|---|---|---|---|---|---|---|---|
| IE1A | Controls | 6 | RS3131643 | 31442782 | A | 0.707 (0.6076-0.8227) | 7,34E-06 |
| IE1A | Controls | 6 | RS3099844 | 31448976 | A | 0.6423 (0.5407-0.7631) | 4,75E-07 |
| IE1A | Controls | 6 | RS3094011 | 31451836 | G | 0.6333 (0.5327-0.7528) | 2,24E-07 |
| IE1A | Controls | 6 | RS9267445 | 31483481 | C | 0.6473 (0.5432-0.7714) | 1,16E-06 |
| IE1A | Controls | 6 | RS2734583 | 31505480 | G | 0.6514 (0.548-0.7744) | 1,18E-06 |
| IE1A | Controls | 6 | RS3117582 | 31620520 | C | 0.658 (0.5522-0.784) | 2,84E-06 |
| IE1A | Controls | 6 | **RS805262** | 31628733 | A | 0.7375 (0.662-0.8215) | **3,22E-08** |
| IE1A | Controls | 6 | RS9267531 | 31636742 | G | 0.6593 (0.5532-0.7859) | 3,34E-06 |
| IE1A | Controls | 6 | RS3101018 | 31705864 | A | 0.6566 (0.5504-0.7832) | 2,94E-06 |
| IE1A | Controls | 6 | RS3132445 | 31712196 | A | 0.6594 (0.5534-0.7857) | 3,21E-06 |
| IE1A | Controls | 6 | RS3130484 | 31715882 | G | 0.6585 (0.5526-0.7846) | 2,98E-06 |
| IE1A | Controls | 6 | RS3131379 | 31721033 | A | 0.6618 (0.5552-0.7889) | 4,13E-06 |
| IE1A | Controls | 6 | RS3117574 | 31725230 | A | 0.6585 (0.5526-0.7847) | 2,98E-06 |
| IE1A | Controls | 6 | RS3131378 | 31725285 | G | 0.6573 (0.5517-0.7833) | 2,71E-06 |
| IE1A | Controls | 6 | RS3117577 | 31727474 | G | 0.6581 (0.5523-0.7842) | 2,91E-06 |
| IE1A | Controls | 6 | RS3101017 | 31733466 | G | 0.6568 (0.5512-0.7827) | 2,62E-06 |
| IE1A | Controls | 6 | RS558702 | 31870326 | A | 0.6619 (0.5555-0.7886) | 3,86E-06 |
| IE1A | Controls | 6 | RS519417 | 31878433 | A | 0.6633 (0.5568-0.7902) | 4,30E-06 |
| IE1A | Controls | 6 | RS511294 | 31888869 | C | 1.728 (1.358-2.2) | 8,68E-06 |
| IE1A | Controls | 6 | RS497309 | 31892484 | C | 0.6607 (0.5546-0.7871) | 3,45E-06 |
| IE1A | Controls | 6 | RS1270942 | 31918860 | G | 0.6584 (0.5525-0.7845) | 2,96E-06 |
| IE1A | Controls | 6 | RS1150755 | 32038550 | A | 0.6912 (0.5938-0.8045) | 1,86E-06 |
| IE1A | Controls | 6 | RS1150754 | 32050758 | A | 0.6911 (0.5937-0.8044) | 1,85E-06 |
| IE1A | Controls | 6 | RS1150753 | 32059867 | G | 0.6595 (0.5538-0.7854) | 3,02E-06 |
| IE1A | Controls | 6 | RS1150752 | 32064726 | G | 0.6407 (0.5371-0.7644) | 7,70E-07 |
| IE1A | Controls | 6 | RS411337 | 32077380 | A | 1.597 (1.309-1.947) | 3,74E-06 |
| IE1A | Controls | 6 | RS1269852 | 32080191 | C | 0.6737 (0.5662-0.8016) | 8,44E-06 |
| IE1A | Controls | 6 | RS9469084 | 32080383 | A | 1.576 (1.293-1.922) | 6,83E-06 |
| IE1A | Controls | 6 | RS204888 | 32089142 | A | 1.576 (1.293-1.921) | 6,86E-06 |
| IE1A | Controls | 6 | RS3130288 | 32096001 | A | 0.6672 (0.5606-0.7941) | 5,19E-06 |
| IE1A | Controls | 6 | RS169494 | 32097876 | A | 1.589 (1.303-1.938) | 4,81E-06 |
| IE1A | Controls | 6 | RS3130349 | 32147696 | A | 0.7151 (0.6173-0.8283) | 7,75E-06 |
| IE1A | Controls | 6 | RS1800625 | 32152442 | G | 0.7157 (0.618-0.8289) | 8,03E-06 |
| IE1A | Controls | 6 | RS2071287 | 32170433 | A | 0.7567 (0.6795-0.8425) | 3,70E-07 |
| IE1A | Controls | 6 | RS2071277 | 32171683 | G | 0.7634 (0.6861-0.8493) | 6,97E-07 |
| IE1A | Controls | 6 | RS206017 | 32176211 | G | 1.335 (1.198-1.488) | 1,89E-07 |
| IE1A | Controls | 6 | RS206018 | 32177880 | C | 1.359 (1.186-1.557) | 9,79E-06 |
| IE1A | Controls | 6 | **RS3131290** | 32183175 | A | 0.7269 (0.6503-0.8126) | **2,03E-08** |
| IE1A | Controls | 6 | **RS3134799** | 32184221 | A | 0.7257 (0.6509-0.8091) | **7,57E-09** |
| IE1A | Controls | 6 | RS436388 | 32186264 | A | 1.343 (1.205-1.497) | 1,03E-07 |
| IE1A | Controls | 6 | RS379464 | 32186348 | A | 1.79 (1.39-2.306) | 6,57E-06 |

| | | | | | | | |
|---|---|---|---|---|---|---|---|
| IE1A | Controls | 6 | **RS394657** | 32187023 | G | 0.7305 (0.6554-0.8143) | **1,45E-08** |
| IE1A | Controls | 6 | RS1109771 | 32187605 | A | 1.303 (1.169-1.452) | 1,71E-06 |
| IE1A | Controls | 6 | RS8192584 | 32189192 | A | 2.057 (1.532-2.76) | 1,56E-06 |
| IE1A | Controls | 6 | RS915894 | 32190390 | C | 0.7427 (0.6617-0.8335) | 4,31E-07 |
| IE1A | Controls | 6 | RS3132971 | 32230256 | C | 0.638 (0.5351-0.7606) | 5,44E-07 |
| IE1A | Controls | 6 | RS7775397 | 32261252 | C | 0.6413 (0.5382-0.7642) | 6,81E-07 |
| IE1A | Controls | 6 | RS9268235 | 32290208 | A | 0.6407 (0.5377-0.7635) | 6,46E-07 |
| IE1A | Controls | 6 | RS2894254 | 32345689 | C | 0.6375 (0.5343-0.7608) | 5,97E-07 |
| IE1A | Controls | 6 | RS3129843 | 32395726 | G | 0.6385 (0.5351-0.7618) | 6,45E-07 |
| IE1A | Controls | 6 | RS9270665 | 32566232 | A | 0.7231 (0.6293-0.8309) | 4,82E-06 |
| IE1A | Controls | 6 | RS3129763 | 32590925 | A | 0.7249 (0.6311-0.8327) | 5,39E-06 |
| IE1A | Controls | 6 | RS9272219 | 32602269 | A | 0.7368 (0.6493-0.8362) | 2,21E-06 |
| IE1A | Controls | 6 | RS9273012 | 32611641 | G | 0.7348 (0.6476-0.8338) | 1,76E-06 |
| IE1A | Controls | 6 | RS17843604 | 32620283 | G | 1.354 (1.211-1.513) | 9,34E-08 |
| IE1A | Controls | 6 | **RS9273349** | 32625869 | A | 1.402 (1.254-1.567) | **2,52E-09** |
| IE1A | Controls | 6 | RS9273363 | 32626272 | A | 0.7303 (0.6489-0.8219) | 1,84E-07 |
| IE1A | Controls | 6 | **RS1063355** | 32627714 | A | 1.404 (1.256-1.569) | **2,25E-09** |
| IE1A | Controls | 6 | RS9273448 | 32627747 | A | 1.325 (1.176-1.493) | 3,91E-06 |
| IE1A | Controls | 6 | RS1794282 | 32666526 | A | 0.6358 (0.5326-0.7589) | 5,30E-07 |
| IE1A | Controls | 6 | RS3135006 | 32667119 | A | 1.333 (1.182-1.503) | 2,67E-06 |
| IE1A | Controls | 6 | RS9276689 | 32751962 | A | 0.6379 (0.5282-0.7703) | 3,00E-06 |
| IE1A | Controls | 6 | RS7762279 | 32755290 | G | 0.6387 (0.5289-0.7711) | 3,13E-06 |
| IE1A | Controls | 6 | RS3819720 | 32804570 | A | 0.7592 (0.6766-0.8518) | 2,72E-06 |
| IE1A | Controls | 6 | RS1480380 | 32913246 | A | 0.626 (0.5105-0.7675) | 6,67E-06 |
| IE1A | All subjects | 1 | RS595980 | 20 827 216 | A | 1.234 (1.126-1.353) | 6,76E-06 |
| IE1A | All subjects | 1 | RS476622 | 20 853 195 | A | 1.246 (1.137-1.366) | 2,58E-06 |
| IE1A | All subjects | 3 | RS7641571 | 162 749 836 | G | 1.389 (1.203-1.604) | 7,61E-06 |
| IE1A | All subjects | 6 | RS7767008 | 28 630 793 | C | 0.8246 (0.7584-0.8966) | 6,27E-06 |
| IE1A | All subjects | 6 | RS1233579 | 28 712 663 | G | 0.7476 (0.6608-0.8459) | 3,88E-06 |
| IE1A | All subjects | 6 | RS1233599 | 28 731 188 | C | 0.7506 (0.6634-0.8492) | 5,26E-06 |
| IE1A | All subjects | 6 | RS1233619 | 28 745 452 | A | 0.754 (0.6662-0.8535) | 7,91E-06 |
| IE1A | All subjects | 6 | RS1311918 | 28 753 646 | A | 0.7552 (0.6671-0.8549) | 9,14E-06 |
| IE1A | All subjects | 6 | RS7767099 | 28 768 698 | A | 0.7534 (0.6656-0.8527) | 7,41E-06 |
| IE1A | All subjects | 6 | RS2765220 | 28 769 671 | G | 0.778 (0.702-0.8622) | 1,70E-06 |
| IE1A | All subjects | 6 | RS3131343 | 28 775 564 | A | 0.7508 (0.6634-0.8498) | 5,71E-06 |
| IE1A | All subjects | 6 | RS4324798 | 28 776 117 | A | 0.7536 (0.6658-0.8531) | 7,71E-06 |
| IE1A | All subjects | 6 | RS3132389 | 28 831 021 | C | 0.7491 (0.6619-0.8478) | 4,78E-06 |
| IE1A | All subjects | 6 | RS3118370 | 28 833 101 | C | 0.7546 (0.6661-0.8548) | 9,58E-06 |
| IE1A | All subjects | 6 | RS3131093 | 28 837 437 | A | 0.7514 (0.664-0.8504) | 5,97E-06 |
| IE1A | All subjects | 6 | RS3132392 | 28 838 629 | G | 0.7513 (0.6639-0.8503) | 5,91E-06 |

| | | | | | | | |
|---|---|---|---|---|---|---|---|
| IE1A | All subjects | 6 | RS3135309 | 28 855 805 | C | 0.7491 (0.6624-0.8473) | 4,26E-06 |
| IE1A | All subjects | 6 | RS3135316 | 28 864 849 | G | 0.7489 (0.6622-0.847) | 4,16E-06 |
| IE1A | All subjects | 6 | RS2230683 | 28 891 176 | G | 0.75 (0.6631-0.8483) | 4,69E-06 |
| IE1A | All subjects | 6 | RS3118361 | 28 898 287 | A | 0.74 (0.6535-0.8379) | 2,04E-06 |
| IE1A | All subjects | 6 | RS3130895 | 28 905 791 | A | 0.7465 (0.66-0.8444) | 3,34E-06 |
| IE1A | All subjects | 6 | RS3131073 | 28 920 972 | A | 0.7428 (0.6565-0.8405) | 2,41E-06 |
| IE1A | All subjects | 6 | RS3130845 | 28 923 367 | G | 0.7479 (0.6613-0.8459) | 3,79E-06 |
| IE1A | All subjects | 6 | RS3130837 | 28 948 092 | A | 0.7403 (0.6545-0.8375) | 1,75E-06 |
| IE1A | All subjects | 6 | RS3129791 | 28 954 293 | A | 0.7471 (0.6605-0.8451) | 3,56E-06 |
| IE1A | All subjects | 6 | RS3130891 | 28 977 217 | C | 0.7455 (0.659-0.8435) | 3,11E-06 |
| IE1A | All subjects | 6 | RS3130893 | 28 980 707 | G | 0.7466 (0.66-0.8446) | 3,38E-06 |
| IE1A | All subjects | 6 | RS3117143 | 29 031 142 | A | 0.7403 (0.6544-0.8375) | 1,78E-06 |
| IE1A | All subjects | 6 | RS3129173 | 29 159 629 | C | 0.7381 (0.6527-0.8347) | 1,31E-06 |
| IE1A | All subjects | 6 | RS3116830 | 29 167 575 | A | 0.7394 (0.6536-0.8365) | 1,61E-06 |
| IE1A | All subjects | 6 | RS3117326 | 29 240 378 | A | 0.7392 (0.6534-0.8362) | 1,58E-06 |
| IE1A | All subjects | 6 | RS3130834 | 29 248 149 | G | 0.7403 (0.6543-0.8375) | 1,77E-06 |
| IE1A | All subjects | 6 | RS3117439 | 29 266 483 | A | 0.7573 (0.6709-0.8548) | 6,82E-06 |
| IE1A | All subjects | 6 | RS3117433 | 29 303 364 | G | 0.7861 (0.7092-0.8714) | 4,60E-06 |
| IE1A | All subjects | 6 | RS3749971 | 29 342 775 | A | 0.7496 (0.6639-0.8463) | 3,26E-06 |
| IE1A | All subjects | 6 | RS9257809 | 29 356 331 | G | 0.7575 (0.6718-0.8541) | 5,78E-06 |
| IE1A | All subjects | 6 | RS442694 | 29 356 687 | A | 0.7487 (0.6636-0.8448) | 2,63E-06 |
| IE1A | All subjects | 6 | RS429479 | 29 372 323 | G | 0.7598 (0.6742-0.8562) | 6,61E-06 |
| IE1A | All subjects | 6 | RS1535039 | 29 411 432 | G | 0.7442 (0.6596-0.8395) | 1,56E-06 |
| IE1A | All subjects | 6 | RS2746149 | 29 435 355 | G | 0.746 (0.6615-0.8413) | 1,79E-06 |
| IE1A | All subjects | 6 | RS2746150 | 29 442 701 | A | 0.7441 (0.6595-0.8394) | 1,54E-06 |
| IE1A | All subjects | 6 | RS404240 | 29 523 957 | G | 0.7435 (0.6592-0.8385) | 1,37E-06 |
| IE1A | All subjects | 6 | RS1235162 | 29 537 224 | G | 0.7433 (0.6608-0.8362) | 7,91E-07 |
| IE1A | All subjects | 6 | RS444189 | 29 605 935 | C | 0.832 (0.7711-0.8977) | 2,09E-06 |
| IE1A | All subjects | 6 | RS396660 | 29 646 165 | A | 0.8102 (0.7486-0.8768) | 1,79E-07 |
| IE1A | All subjects | 6 | RS445150 | 29 646 879 | G | 0.809 (0.7476-0.8756) | 1,49E-07 |
| IE1A | All subjects | 6 | RS2747430 | 29 648 506 | A | 0.81 (0.7483-0.8768) | 1,85E-07 |
| IE1A | All subjects | 6 | RS3117301 | 29 654 700 | A | 0.8207 (0.753-0.8946) | 6,94E-06 |
| IE1A | All subjects | 6 | RS2747457 | 29 656 417 | C | 0.8201 (0.7525-0.8938) | 6,32E-06 |
| IE1A | All subjects | 6 | RS2747460 | 29 657 127 | A | 0.8209 (0.7532-0.8948) | 7,10E-06 |
| IE1A | All subjects | 6 | RS3129039 | 29 658 864 | A | 0.8185 (0.7494-0.8939) | 8,36E-06 |
| IE1A | All subjects | 6 | RS3094736 | 29 662 374 | T | 0.8409 (0.7813-0.9051) | 3,94E-06 |
| IE1A | All subjects | 6 | RS3129066 | 29 668 537 | G | 0.8285 (0.7625-0.9001) | 8,83E-06 |
| IE1A | All subjects | 6 | RS2107203 | 29 669 830 | A | 0.8277 (0.7617-0.8993) | 8,05E-06 |
| IE1A | All subjects | 6 | RS3129055 | 29 670 261 | G | 0.8272 (0.7613-0.8988) | 7,55E-06 |
| IE1A | All subjects | 6 | RS9258122 | 29 671 740 | A | 0.827 (0.7611-0.8987) | 7,40E-06 |
| IE1A | All subjects | 6 | RS1633041 | 29 733 223 | A | 0.8265 (0.7625-0.8958) | 3,57E-06 |

| | | | | | | | |
|---|---|---|---|---|---|---|---|
| IE1A | All subjects | 6 | RS1737041 | 29 736 229 | A | 0.8264 (0.7624-0.8957) | 3,50E-06 |
| IE1A | All subjects | 6 | RS1737031 | 29 737 481 | A | 0.8228 (0.7599-0.8908) | 1,50E-06 |
| IE1A | All subjects | 6 | RS1362068 | 29 742 108 | G | 0.8364 (0.7739-0.9038) | 6,36E-06 |
| IE1A | All subjects | 6 | RS1362070 | 29 742 299 | G | 0.836 (0.7736-0.9034) | 6,01E-06 |
| IE1A | All subjects | 6 | RS1615962 | 29 745 726 | A | 0.8303 (0.7659-0.9) | 6,19E-06 |
| IE1A | All subjects | 6 | RS1633030 | 29 745 794 | G | 0.8224 (0.7595-0.8904) | 1,42E-06 |
| IE1A | All subjects | 6 | RS1633028 | 29 746 367 | C | 0.823 (0.7601-0.8911) | 1,57E-06 |
| IE1A | All subjects | 6 | RS2735042 | 29 747 373 | G | 0.8231 (0.7602-0.8912) | 1,58E-06 |
| IE1A | All subjects | 6 | RS1002046 | 29 754 016 | A | 0.8273 (0.7632-0.8968) | 4,05E-06 |
| IE1A | All subjects | 6 | RS1610641 | 29 759 066 | G | 0.834 (0.7714-0.9016) | 5,00E-06 |
| IE1A | All subjects | 6 | RS1610644 | 29 759 445 | A | 0.8277 (0.7636-0.8971) | 4,24E-06 |
| IE1A | All subjects | 6 | RS1611205 | 29 759 823 | G | 0.8308 (0.7683-0.8984) | 3,39E-06 |
| IE1A | All subjects | 6 | RS1610719 | 29 760 677 | G | 0.8336 (0.7711-0.9012) | 4,72E-06 |
| IE1A | All subjects | 6 | RS1610723 | 29 761 473 | C | 0.8308 (0.7685-0.8982) | 3,17E-06 |
| IE1A | All subjects | 6 | RS1610724 | 29 761 516 | G | 0.8316 (0.7692-0.899) | 3,54E-06 |
| IE1A | All subjects | 6 | RS1611220 | 29 761 835 | G | 0.8376 (0.7751-0.9051) | 7,54E-06 |
| IE1A | All subjects | 6 | RS1633011 | 29 762 693 | A | 0.828 (0.7638-0.8977) | 4,70E-06 |
| IE1A | All subjects | 6 | RS1633005 | 29 764 472 | A | 0.7729 (0.7013-0.8517) | 2,02E-07 |
| IE1A | All subjects | 6 | RS1633003 | 29 764 759 | G | 0.8395 (0.7768-0.9073) | 9,99E-06 |
| IE1A | All subjects | 6 | RS1630969 | 29 765 001 | A | 0.8291 (0.7636-0.9002) | 8,08E-06 |
| IE1A | All subjects | 6 | **RS1610649** | 29 768 917 | G | 0.8037 (0.7437-0.8686) | **3,43E-08** |
| IE1A | All subjects | 6 | RS1610711 | 29 769 497 | C | 0.8266 (0.7626-0.8961) | 3,73E-06 |
| IE1A | All subjects | 6 | RS1610657 | 29 771 066 | G | 0.838 (0.7754-0.9056) | 7,96E-06 |
| IE1A | All subjects | 6 | RS1611192 | 29 771 879 | G | 0.8167 (0.7491-0.8904) | 4,32E-06 |
| IE1A | All subjects | 6 | RS1632988 | 29 772 395 | A | 0.8291 (0.7633-0.9005) | 8,78E-06 |
| IE1A | All subjects | 6 | RS1632987 | 29 772 549 | G | 0.8274 (0.7633-0.8969) | 4,15E-06 |
| IE1A | All subjects | 6 | RS1632983 | 29 773 509 | A | 0.8288 (0.7658-0.897) | 3,25E-06 |
| IE1A | All subjects | 6 | RS1736976 | 29 773 999 | G | 0.8316 (0.7693-0.8989) | 3,46E-06 |
| IE1A | All subjects | 6 | RS1736971 | 29 776 322 | A | 0.8282 (0.7641-0.8978) | 4,60E-06 |
| IE1A | All subjects | 6 | RS1736969 | 29 776 390 | A | 0.8236 (0.7597-0.8928) | 2,47E-06 |
| IE1A | All subjects | 6 | RS1610669 | 29 780 093 | A | 0.8376 (0.7751-0.9052) | 7,62E-06 |
| IE1A | All subjects | 6 | RS1736957 | 29 782 633 | G | 0.833 (0.7692-0.9022) | 7,13E-06 |
| IE1A | All subjects | 6 | RS1077432 | 29 782 868 | G | 0.8267 (0.7625-0.8962) | 3,81E-06 |
| IE1A | All subjects | 6 | RS1620173 | 29 785 149 | C | 0.8281 (0.7639-0.8976) | 4,51E-06 |
| IE1A | All subjects | 6 | RS1619379 | 29 785 235 | A | 0.8165 (0.7574-0.8802) | 1,24E-07 |
| IE1A | All subjects | 6 | **RS1610678** | 29 789 190 | G | 0.7948 (0.7385-0.8555) | **9,49E-10** |
| IE1A | All subjects | 6 | **RS1611149** | 29 789 999 | A | 0.7943 (0.7381-0.8549) | **7,93E-10** |
| IE1A | All subjects | 6 | **RS1736936** | 29 794 317 | A | 0.7882 (0.7321-0.8485) | **2,54E-10** |
| IE1A | All subjects | 6 | **RS915669** | 29 798 425 | A | 0.7938 (0.7376-0.8543) | **7,27E-10** |
| IE1A | All subjects | 6 | RS1611133 | 29 809 382 | A | 0.7953 (0.7286-0.8681) | 2,95E-07 |
| IE1A | All subjects | 6 | RS1611750 | 29 814 778 | C | 0.7921 (0.7256-0.8647) | 1,89E-07 |

| | | | | | | | |
|---|---|---|---|---|---|---|---|
| IE1A | All subjects | 6 | **RS2734985** | 29 818 662 | G | 0.7811 (0.7147-0.8536) | **4,91E-08** |
| IE1A | All subjects | 6 | RS2247504 | 29 819 077 | A | 0.8177 (0.7589-0.8809) | 1,19E-07 |
| IE1A | All subjects | 6 | RS2523759 | 29 819 093 | G | 0.7853 (0.7187-0.8579) | 8,60E-08 |
| IE1A | All subjects | 6 | RS5013093 | 29 820 586 | A | 0.7849 (0.7184-0.8575) | 8,00E-08 |
| IE1A | All subjects | 6 | RS2523756 | 29 820 717 | G | 0.8152 (0.7568-0.8781) | 7,15E-08 |
| IE1A | All subjects | 6 | RS2734981 | 29 821 692 | A | 0.7877 (0.7209-0.8607) | 1,31E-07 |
| IE1A | All subjects | 6 | RS2734980 | 29 821 896 | A | 0.785 (0.7186-0.8577) | 8,25E-08 |
| IE1A | All subjects | 6 | RS2517862 | 29 821 937 | A | 0.7847 (0.7097-0.8676) | 2,22E-06 |
| IE1A | All subjects | 6 | RS2517861 | 29 821 982 | A | 0.785 (0.7185-0.8575) | 8,00E-08 |
| IE1A | All subjects | 6 | RS2517860 | 29 822 521 | G | 0.7841 (0.7178-0.8565) | 6,72E-08 |
| IE1A | All subjects | 6 | RS2428510 | 29 823 027 | A | 0.8154 (0.757-0.8783) | 7,39E-08 |
| IE1A | All subjects | 6 | RS2517855 | 29 823 164 | A | 0.7865 (0.7114-0.8695) | 2,73E-06 |
| IE1A | All subjects | 6 | **RS2517854** | 29 823 296 | G | 0.8102 (0.752-0.8729) | **3,11E-08** |
| IE1A | All subjects | 6 | RS2844826 | 29 823 319 | G | 0.8156 (0.7572-0.8784) | 7,47E-08 |
| IE1A | All subjects | 6 | RS6919513 | 29 823 994 | A | 0.8166 (0.7581-0.8797) | 9,49E-08 |
| IE1A | All subjects | 6 | RS1611684 | 29 825 846 | A | 0.8156 (0.7572-0.8784) | 7,47E-08 |
| IE1A | All subjects | 6 | RS1611717 | 29 829 577 | G | 0.8158 (0.7574-0.8786) | 7,69E-08 |
| IE1A | All subjects | 6 | RS1611732 | 29 831 008 | A | 0.8149 (0.7566-0.8778) | 6,68E-08 |
| IE1A | All subjects | 6 | RS765649 | 29 831 058 | A | 0.8175 (0.7588-0.8807) | 1,13E-07 |
| IE1A | All subjects | 6 | **RS1611737** | 29 831 571 | G | 0.8124 (0.7541-0.8752) | **4,54E-08** |
| IE1A | All subjects | 6 | RS1611738 | 29 831 706 | A | 0.7853 (0.7102-0.8684) | 2,49E-06 |
| IE1A | All subjects | 6 | RS1655900 | 29 916 618 | A | 0.7815 (0.7058-0.8653) | 2,12E-06 |
| IE1A | All subjects | 6 | RS2844806 | 29 933 439 | A | 0.8409 (0.7817-0.9046) | 3,33E-06 |
| IE1A | All subjects | 6 | RS3893464 | 29 935 250 | A | 0.8308 (0.772-0.8942) | 7,60E-07 |
| IE1A | All subjects | 6 | RS3132685 | 29 945 949 | A | 0.7704 (0.6893-0.861) | 4,31E-06 |
| IE1A | All subjects | 6 | RS8321 | 30 032 522 | C | 0.7291 (0.6478-0.8206) | 1,61E-07 |
| IE1A | All subjects | 6 | RS9261290 | 30 038 647 | G | 0.7287 (0.6474-0.8201) | 1,53E-07 |
| IE1A | All subjects | 6 | RS3132682 | 30 044 388 | C | 1.217 (1.131-1.31) | 1,61E-07 |
| IE1A | All subjects | 6 | RS7382061 | 30 047 965 | A | 1.219 (1.133-1.312) | 1,32E-07 |
| IE1A | All subjects | 6 | RS2057728 | 30 054 757 | A | 1.222 (1.135-1.316) | 9,89E-08 |
| IE1A | All subjects | 6 | RS6904455 | 30 055 377 | G | 1.22 (1.133-1.313) | 1,35E-07 |
| IE1A | All subjects | 6 | RS1264706 | 30 063 652 | G | 0.7309 (0.6495-0.8225) | 1,97E-07 |
| IE1A | All subjects | 6 | **RS9261394** | 30 064 562 | A | 1.229 (1.142-1.323) | **4,05E-08** |
| IE1A | All subjects | 6 | RS1116222 | 30 071 279 | C | 0.8265 (0.7603-0.8985) | 7,75E-06 |
| IE1A | All subjects | 6 | RS2523989 | 30 078 275 | A | 0.7725 (0.701-0.8513) | 1,92E-07 |
| IE1A | All subjects | 6 | RS2239529 | 30 078 330 | A | 0.7791 (0.7069-0.8586) | 4,77E-07 |
| IE1A | All subjects | 6 | RS2523988 | 30 079 129 | G | 0.7697 (0.6963-0.8509) | 3,09E-07 |
| IE1A | All subjects | 6 | RS2249099 | 30 079 307 | A | 0.7703 (0.6993-0.8484) | 1,21E-07 |
| IE1A | All subjects | 6 | RS2523987 | 30 079 993 | C | 0.7705 (0.697-0.8518) | 3,48E-07 |
| IE1A | All subjects | 6 | RS2517598 | 30 080 274 | A | 0.7715 (0.7004-0.8498) | 1,44E-07 |
| IE1A | All subjects | 6 | RS2844793 | 30 080 496 | A | 0.7785 (0.7065-0.8578) | 4,25E-07 |

| | | | | | | | |
|---|---|---|---|---|---|---|---|
| IE1A | All subjects | 6 | RS2517597 | 30 081 189 | A | 0.7713 (0.6977-0.8526) | 3,83E-07 |
| IE1A | All subjects | 6 | RS2523986 | 30 081 246 | A | 0.7716 (0.698-0.853) | 4,03E-07 |
| IE1A | All subjects | 6 | RS2523985 | 30 081 334 | G | 0.7713 (0.7002-0.8495) | 1,39E-07 |
| IE1A | All subjects | 6 | RS2523984 | 30 082 003 | A | 0.7713 (0.7002-0.8495) | 1,39E-07 |
| IE1A | All subjects | 6 | RS2245420 | 30 082 688 | C | 0.7657 (0.695-0.8436) | 6,68E-08 |
| IE1A | All subjects | 6 | RS2523981 | 30 083 182 | A | 0.7732 (0.6996-0.8547) | 4,81E-07 |
| IE1A | All subjects | 6 | RS2523979 | 30 083 515 | A | 0.7899 (0.7157-0.8717) | 2,75E-06 |
| IE1A | All subjects | 6 | RS1362104 | 30 101 656 | A | 0.8261 (0.7666-0.8903) | 5,67E-07 |
| IE1A | All subjects | 6 | **RS2517645** | 30 122 623 | G | 0.7547 (0.6821-0.835) | **4,93E-08** |
| IE1A | All subjects | 6 | RS929157 | 30 137 209 | A | 0.7474 (0.6714-0.832) | 1,02E-07 |
| IE1A | All subjects | 6 | RS2106072 | 30 153 363 | A | 0.7212 (0.6408-0.8117) | 5,99E-08 |
| IE1A | All subjects | 6 | RS2517617 | 30 161 536 | A | 0.7211 (0.6406-0.8116) | 5,94E-08 |
| IE1A | All subjects | 6 | RS2517614 | 30 163 955 | A | 0.7822 (0.7082-0.8641) | 1,31E-06 |
| IE1A | All subjects | 6 | RS2523722 | 30 165 273 | A | 0.7824 (0.7083-0.8643) | 1,35E-06 |
| IE1A | All subjects | 6 | RS2517613 | 30 165 662 | A | 0.7496 (0.6734-0.8346) | 1,42E-07 |
| IE1A | All subjects | 6 | RS2523721 | 30 166 266 | A | 0.7802 (0.7066-0.8615) | 9,13E-07 |
| IE1A | All subjects | 6 | RS2523719 | 30 168 319 | A | 0.7444 (0.6691-0.8281) | 5,79E-08 |
| IE1A | All subjects | 6 | RS2517612 | 30 169 092 | G | 0.7475 (0.6715-0.8321) | 1,04E-07 |
| IE1A | All subjects | 6 | RS2517611 | 30 169 327 | G | 0.7807 (0.7068-0.8624) | 1,07E-06 |
| IE1A | All subjects | 6 | RS2517610 | 30 170 280 | G | 0.7764 (0.7032-0.8572) | 5,39E-07 |
| IE1A | All subjects | 6 | RS1117490 | 30 170 510 | G | 0.777 (0.7038-0.8578) | 5,80E-07 |
| IE1A | All subjects | 6 | RS2844776 | 30 171 827 | G | 0.7761 (0.703-0.8569) | 5,16E-07 |
| IE1A | All subjects | 6 | RS971570 | 30 172 513 | C | 0.7733 (0.7003-0.8539) | 3,75E-07 |
| IE1A | All subjects | 6 | RS2021722 | 30 174 131 | A | 0.7754 (0.7023-0.856) | 4,68E-07 |
| IE1A | All subjects | 6 | RS885912 | 30 174 633 | A | 0.7768 (0.7036-0.8575) | 5,58E-07 |
| IE1A | All subjects | 6 | RS2188100 | 30 181 883 | A | 0.7483 (0.6722-0.833) | 1,17E-07 |
| IE1A | All subjects | 6 | RS885916 | 30 202 571 | A | 0.7469 (0.671-0.8314) | 9,48E-08 |
| IE1A | All subjects | 6 | RS3094078 | 30 224 970 | T | 0.7225 (0.6418-0.8133) | 7,41E-08 |
| IE1A | All subjects | 6 | **RS3094073** | 30 231 224 | A | 0.7442 (0.6718-0.8243) | **1,48E-08** |
| IE1A | All subjects | 6 | **RS3130401** | 30 231 273 | G | 0.7448 (0.6723-0.825) | **1,65E-08** |
| IE1A | All subjects | 6 | **RS3132659** | 30 231 330 | G | 0.7456 (0.673-0.8259) | **1,86E-08** |
| IE1A | All subjects | 6 | **RS3132658** | 30 231 666 | G | 0.7434 (0.6711-0.8236) | **1,36E-08** |
| IE1A | All subjects | 6 | **RS3094071** | 30 231 768 | A | 0.7443 (0.6719-0.8245) | **1,54E-08** |
| IE1A | All subjects | 6 | **RS3130403** | 30 232 009 | C | 0.7433 (0.671-0.8235) | **1,37E-08** |
| IE1A | All subjects | 6 | **RS3094630** | 30 232 436 | A | 0.7448 (0.6724-0.8251) | **1,66E-08** |
| IE1A | All subjects | 6 | **RS3129702** | 30 232 785 | A | 0.7448 (0.6724-0.8251) | **1,66E-08** |
| IE1A | All subjects | 6 | **RS3094629** | 30 232 953 | A | 0.7448 (0.6723-0.8251) | **1,70E-08** |
| IE1A | All subjects | 6 | **RS3129703** | 30 233 558 | A | 0.7443 (0.6719-0.8245) | **1,54E-08** |
| IE1A | All subjects | 6 | **RS3130405** | 30 234 152 | A | 0.7447 (0.6723-0.825) | **1,64E-08** |
| IE1A | All subjects | 6 | **RS3129832** | 30 234 657 | G | 0.739 (0.667-0.8189) | **7,64E-09** |
| IE1A | All subjects | 6 | **RS3129705** | 30 234 721 | A | 0.7446 (0.6722-0.8248) | **1,60E-08** |

| | | | | | | | |
|---|---|---|---|---|---|---|---|
| IE1A | All subjects | 6 | RS3129830 | 30 248 711 | A | 0.7216 (0.6413-0.8119) | 5,90E-08 |
| IE1A | All subjects | 6 | **RS3130380** | 30 279 130 | A | 0.7157 (0.6361-0.8053) | **2,71E-08** |
| IE1A | All subjects | 6 | **RS3094064** | 30 296 253 | A | 0.7188 (0.6391-0.8084) | **3,64E-08** |
| IE1A | All subjects | 6 | RS3094067 | 30 299 245 | C | 0.7225 (0.6421-0.8129) | 6,57E-08 |
| IE1A | All subjects | 6 | **RS3129837** | 30 306 306 | G | 0.7475 (0.6746-0.8282) | **2,71E-08** |
| IE1A | All subjects | 6 | **RS3129840** | 30 307 342 | G | 0.7491 (0.676-0.8301) | **3,49E-08** |
| IE1A | All subjects | 6 | RS3132649 | 30 321 057 | A | 0.7343 (0.6549-0.8234) | 1,24E-07 |
| IE1A | All subjects | 6 | RS3094061 | 30 321 189 | C | 0.7343 (0.6549-0.8234) | 1,23E-07 |
| IE1A | All subjects | 6 | RS3130374 | 30 321 336 | A | 0.7343 (0.6549-0.8234) | 1,25E-07 |
| IE1A | All subjects | 6 | RS3130375 | 30 321 732 | A | 0.7359 (0.6562-0.8252) | 1,55E-07 |
| IE1A | All subjects | 6 | RS3130350 | 30 327 839 | A | 0.7242 (0.6441-0.8144) | 7,02E-08 |
| IE1A | All subjects | 6 | RS3094622 | 30 327 952 | G | 0.7333 (0.6538-0.8226) | 1,21E-07 |
| IE1A | All subjects | 6 | RS3130351 | 30 328 192 | A | 0.724 (0.6438-0.8141) | 6,87E-08 |
| IE1A | All subjects | 6 | **RS3130352** | 30 328 357 | A | 0.7212 (0.6414-0.811) | **4,77E-08** |
| IE1A | All subjects | 6 | RS3094621 | 30 328 753 | G | 0.7347 (0.655-0.8242) | 1,46E-07 |
| IE1A | All subjects | 6 | RS3132647 | 30 330 737 | G | 0.7256 (0.6452-0.8161) | 8,74E-08 |
| IE1A | All subjects | 6 | RS3094054 | 30 333 505 | A | 0.7569 (0.675-0.8487) | 1,86E-06 |
| IE1A | All subjects | 6 | RS3129809 | 30 335 621 | G | 0.7413 (0.6612-0.8312) | 2,96E-07 |
| IE1A | All subjects | 6 | RS3129817 | 30 342 753 | A | 0.745 (0.6655-0.834) | 3,19E-07 |
| IE1A | All subjects | 6 | RS3129820 | 30 343 569 | A | 0.7412 (0.6611-0.831) | 2,87E-07 |
| IE1A | All subjects | 6 | RS3132631 | 30 344 645 | A | 0.7402 (0.6602-0.8299) | 2,56E-07 |
| IE1A | All subjects | 6 | RS3132630 | 30 345 118 | A | 0.7418 (0.6613-0.8322) | 3,50E-07 |
| IE1A | All subjects | 6 | RS3129822 | 30 346 208 | A | 0.7434 (0.6627-0.8339) | 4,19E-07 |
| IE1A | All subjects | 6 | RS3132625 | 30 347 720 | G | 0.7413 (0.6611-0.8311) | 2,92E-07 |
| IE1A | All subjects | 6 | RS3094032 | 30 351 547 | A | 0.7394 (0.6594-0.8291) | 2,34E-07 |
| IE1A | All subjects | 6 | RS3130126 | 30 353 739 | A | 0.7408 (0.6606-0.8306) | 2,79E-07 |
| IE1A | All subjects | 6 | RS3094050 | 30 358 591 | G | 0.7416 (0.6615-0.8314) | 2,95E-07 |
| IE1A | All subjects | 6 | RS3130116 | 30 365 448 | G | 0.7521 (0.6718-0.8421) | 7,72E-07 |
| IE1A | All subjects | 6 | RS3130141 | 30 432 177 | A | 0.7573 (0.6761-0.8482) | 1,53E-06 |
| IE1A | All subjects | 6 | RS3094694 | 30 451 904 | G | 0.7924 (0.7222-0.8695) | 8,94E-07 |
| IE1A | All subjects | 6 | RS3130117 | 30 508 956 | A | 0.7585 (0.6776-0.8491) | 1,57E-06 |
| IE1A | All subjects | 6 | RS3130247 | 30 515 043 | G | 0.7571 (0.6764-0.8474) | 1,31E-06 |
| IE1A | All subjects | 6 | RS3132610 | 30 544 401 | G | 0.7614 (0.6811-0.8511) | 1,62E-06 |
| IE1A | All subjects | 6 | RS9262132 | 30 611 350 | A | 0.7521 (0.6721-0.8416) | 6,88E-07 |
| IE1A | All subjects | 6 | RS9262135 | 30 618 906 | G | 0.7499 (0.6701-0.8392) | 5,36E-07 |
| IE1A | All subjects | 6 | RS9262141 | 30 644 137 | G | 0.7569 (0.6763-0.8471) | 1,24E-06 |
| IE1A | All subjects | 6 | RS9262142 | 30 650 026 | A | 0.7621 (0.6807-0.8531) | 2,36E-06 |
| IE1A | All subjects | 6 | RS9262143 | 30 652 781 | A | 0.7577 (0.6772-0.8478) | 1,28E-06 |
| IE1A | All subjects | 6 | RS3132585 | 30 687 614 | G | 0.7507 (0.6711-0.8398) | 5,40E-07 |
| IE1A | All subjects | 6 | RS3132583 | 30 688 575 | C | 0.75 (0.6704-0.839) | 4,95E-07 |
| IE1A | All subjects | 6 | RS3132582 | 30 689 001 | T | 0.7679 (0.6881-0.857) | 2,39E-06 |

| | | | | | | | |
|---|---|---|---|---|---|---|---|
| IE1A | All subjects | 6 | RS1059612 | 30 708 955 | A | 0.7573 (0.6767-0.8475) | 1,29E-06 |
| IE1A | All subjects | 6 | RS3129973 | 30 721 143 | A | 0.7413 (0.6628-0.829) | 1,54E-07 |
| IE1A | All subjects | 6 | RS3095336 | 30 738 446 | A | 0.7628 (0.6823-0.8527) | 1,92E-06 |
| IE1A | All subjects | 6 | RS3130673 | 30 746 519 | A | 0.7654 (0.6847-0.8557) | 2,59E-06 |
| IE1A | All subjects | 6 | RS3131050 | 30 760 025 | G | 0.7687 (0.6877-0.8593) | 3,73E-06 |
| IE1A | All subjects | 6 | RS3131060 | 30 763 291 | A | 0.7674 (0.6864-0.8578) | 3,22E-06 |
| IE1A | All subjects | 6 | RS3129986 | 30 763 562 | A | 0.7844 (0.7062-0.8712) | 5,83E-06 |
| IE1A | All subjects | 6 | RS3130641 | 30 764 081 | A | 0.7659 (0.6851-0.8562) | 2,73E-06 |
| IE1A | All subjects | 6 | RS1264377 | 30 764 907 | A | 0.7824 (0.7049-0.8684) | 4,02E-06 |
| IE1A | All subjects | 6 | RS1264376 | 30 765 579 | A | 0.7818 (0.7027-0.8697) | 6,01E-06 |
| IE1A | All subjects | 6 | **RS1264341** | 30 802 465 | G | 0.727 (0.6496-0.8136) | **2,83E-08** |
| IE1A | All subjects | 6 | **RS2535340** | 30 838 497 | G | 0.7261 (0.649-0.8125) | **2,37E-08** |
| IE1A | All subjects | 6 | **RS1264322** | 30 857 894 | A | 0.728 (0.6506-0.8146) | **3,10E-08** |
| IE1A | All subjects | 6 | **RS886422** | 30 864 279 | A | 0.7281 (0.6507-0.8147) | **3,14E-08** |
| IE1A | All subjects | 6 | **RS1049633** | 30 867 527 | A | 0.7268 (0.6494-0.8134) | **2,76E-08** |
| IE1A | All subjects | 6 | **RS1264312** | 30 872 982 | A | 0.7238 (0.6464-0.8105) | **2,11E-08** |
| IE1A | All subjects | 6 | **RS1264310** | 30 873 605 | A | 0.7285 (0.6508-0.8156) | **3,77E-08** |
| IE1A | All subjects | 6 | **RS1264308** | 30 879 987 | A | 0.7277 (0.6503-0.8142) | **2,95E-08** |
| IE1A | All subjects | 6 | **RS1264304** | 30 882 415 | A | 0.726 (0.6488-0.8123) | **2,32E-08** |
| IE1A | All subjects | 6 | RS3131921 | 30 907 335 | G | 0.7544 (0.6782-0.8393) | 2,18E-07 |
| IE1A | All subjects | 6 | RS3130782 | 30 914 843 | A | 0.7571 (0.6802-0.8428) | 3,59E-07 |
| IE1A | All subjects | 6 | **RS3094086** | 30 919 391 | A | 0.7371 (0.6605-0.8225) | **4,98E-08** |
| IE1A | All subjects | 6 | **RS3131934** | 30 931 844 | G | 0.7382 (0.6641-0.8207) | **1,94E-08** |
| IE1A | All subjects | 6 | **RS3131783** | 30 932 068 | A | 0.7346 (0.6607-0.8169) | **1,22E-08** |
| IE1A | All subjects | 6 | RS3132579 | 30 940 989 | G | 0.8026 (0.7295-0.8829) | 6,30E-06 |
| IE1A | All subjects | 6 | RS1634714 | 30 951 561 | G | 0.8029 (0.7298-0.8833) | 6,57E-06 |
| IE1A | All subjects | 6 | **RS1634721** | 30 977 680 | A | 0.7114 (0.6361-0.7956) | **2,42E-09** |
| IE1A | All subjects | 6 | **RS3130544** | 31 058 340 | A | 0.6939 (0.6209-0.7756) | **1,23E-10** |
| IE1A | All subjects | 6 | RS2233974 | 31 080 016 | C | 0.7656 (0.6941-0.8446) | 9,64E-08 |
| IE1A | All subjects | 6 | RS2233956 | 31 081 205 | G | 0.7896 (0.7169-0.8696) | 1,60E-06 |
| IE1A | All subjects | 6 | **RS3130557** | 31 094 703 | A | 0.6862 (0.6138-0.7672) | **3,68E-11** |
| IE1A | All subjects | 6 | **RS7750641** | 31 129 310 | A | 0.6882 (0.6157-0.7692) | **4,71E-11** |
| IE1A | All subjects | 6 | **RS3132510** | 31 172 151 | G | 0.6642 (0.5911-0.7463) | **6,01E-12** |
| IE1A | All subjects | 6 | RS9263911 | 31 175 667 | G | 1.181 (1.097-1.27) | 9,14E-06 |
| IE1A | All subjects | 6 | RS3132505 | 31 177 503 | A | 0.8189 (0.7497-0.8945) | 9,16E-06 |
| IE1A | All subjects | 6 | RS2394895 | 31 206 979 | G | 0.817 (0.748-0.8925) | 7,31E-06 |
| IE1A | All subjects | 6 | RS2394944 | 31 220 450 | A | 0.8062 (0.7369-0.882) | 2,61E-06 |
| IE1A | All subjects | 6 | RS1793891 | 31 221 698 | A | 0.8105 (0.7433-0.8838) | 1,96E-06 |
| IE1A | All subjects | 6 | RS1986997 | 31 228 410 | A | 0.847 (0.7868-0.9118) | 9,97E-06 |
| IE1A | All subjects | 6 | RS2245822 | 31 230 800 | A | 0.8087 (0.7407-0.8831) | 2,23E-06 |
| IE1A | All subjects | 6 | RS1049281 | 31 236 567 | A | 0.8232 (0.7636-0.8874) | 3,88E-07 |

| | | | | | | | |
|---|---|---|---|---|---|---|---|
| IE1A | All subjects | 6 | **RS2844613** | 31 243 846 | A | 0.7359 (0.6645-0.815) | **3,93E-09** |
| IE1A | All subjects | 6 | RS3130696 | 31 243 884 | A | 0.8002 (0.7319-0.8748) | 9,67E-07 |
| IE1A | All subjects | 6 | RS2524074 | 31 244 021 | G | 0.8119 (0.7521-0.8763) | 9,03E-08 |
| IE1A | All subjects | 6 | RS2844603 | 31 250 854 | A | 0.8402 (0.7804-0.9047) | 3,95E-06 |
| IE1A | All subjects | 6 | RS2853933 | 31 254 088 | A | 0.8443 (0.7842-0.909) | 7,01E-06 |
| IE1A | All subjects | 6 | RS2524040 | 31 257 625 | A | 0.8391 (0.7793-0.9035) | 3,32E-06 |
| IE1A | All subjects | 6 | RS2524163 | 31 259 579 | G | 0.8438 (0.7837-0.9085) | 6,58E-06 |
| IE1A | All subjects | 6 | RS2524156 | 31 260 397 | A | 0.8441 (0.7839-0.9088) | 6,98E-06 |
| IE1A | All subjects | 6 | RS2243868 | 31 261 276 | A | 0.8431 (0.7831-0.9078) | 5,99E-06 |
| IE1A | All subjects | 6 | **RS2247056** | 31 265 490 | A | 0.7866 (0.7273-0.8508) | **1,95E-09** |
| IE1A | All subjects | 6 | RS2853922 | 31 266 190 | A | 0.8407 (0.7807-0.9052) | 4,28E-06 |
| IE1A | All subjects | 6 | RS2524089 | 31 266 522 | C | 0.8395 (0.7796-0.9039) | 3,52E-06 |
| IE1A | All subjects | 6 | RS2524066 | 31 269 154 | A | 0.8398 (0.78-0.9042) | 3,62E-06 |
| IE1A | All subjects | 6 | **RS2596487** | 31 325 056 | A | 0.7058 (0.6385-0.7801) | **9,07E-12** |
| IE1A | All subjects | 6 | **RS2523573** | 31 328 988 | C | 0.6463 (0.5772-0.7237) | **3,86E-14** |
| IE1A | All subjects | 6 | RS2844575 | 31 334 945 | G | 1.191 (1.107-1.281) | 2,66E-06 |
| IE1A | All subjects | 6 | RS3094600 | 31 347 144 | A | 0.8221 (0.7555-0.8946) | 5,50E-06 |
| IE1A | All subjects | 6 | **RS9266669** | 31 348 077 | A | 0.6732 (0.6056-0.7484) | **2,37E-13** |
| IE1A | All subjects | 6 | RS3094604 | 31 434 111 | G | 0.8006 (0.7267-0.882) | 6,74E-06 |
| IE1A | All subjects | 6 | **RS3094013** | 31 434 366 | A | 0.6485 (0.5793-0.726) | **5,65E-14** |
| IE1A | All subjects | 6 | **RS3131618** | 31 434 621 | G | 0.6468 (0.5773-0.7246) | **5,68E-14** |
| IE1A | All subjects | 6 | RS2518028 | 31 436 047 | A | 0.8337 (0.7733-0.8987) | 2,08E-06 |
| IE1A | All subjects | 6 | RS1055569 | 31 440 082 | A | 1.205 (1.11-1.308) | 7,86E-06 |
| IE1A | All subjects | 6 | RS3828886 | 31 440 552 | C | 1.235 (1.134-1.346) | 1,38E-06 |
| IE1A | All subjects | 6 | **RS3131643** | 31 442 782 | A | 0.6932 (0.6259-0.7677) | **2,02E-12** |
| IE1A | All subjects | 6 | **RS3099844** | 31 448 976 | A | 0.636 (0.5683-0.7118) | **3,38E-15** |
| IE1A | All subjects | 6 | RS2523706 | 31 451 567 | G | 0.8361 (0.7737-0.9037) | 6,29E-06 |
| IE1A | All subjects | 6 | **RS3094011** | 31 451 836 | G | 0.6345 (0.5667-0.7104) | **3,02E-15** |
| IE1A | All subjects | 6 | RS2516504 | 31 452 475 | A | 0.839 (0.7763-0.9067) | 9,37E-06 |
| IE1A | All subjects | 6 | RS2516408 | 31 463 491 | A | 0.8349 (0.7729-0.902) | 4,68E-06 |
| IE1A | All subjects | 6 | **RS2855812** | 31 472 720 | A | 0.7856 (0.7236-0.8531) | **9,28E-09** |
| IE1A | All subjects | 6 | RS2516400 | 31 481 105 | A | 0.8346 (0.7725-0.9016) | 4,44E-06 |
| IE1A | All subjects | 6 | **RS9267445** | 31 483 481 | C | 0.6446 (0.5749-0.7228) | **5,43E-14** |
| IE1A | All subjects | 6 | **RS3093988** | 31 492 453 | A | 0.7487 (0.6781-0.8267) | **1,03E-08** |
| IE1A | All subjects | 6 | RS2734573 | 31 494 738 | A | 0.8419 (0.7817-0.9067) | 5,48E-06 |
| IE1A | All subjects | 6 | RS2516482 | 31 496 569 | G | 0.7624 (0.6907-0.8414) | 7,12E-08 |
| IE1A | All subjects | 6 | **RS2734583** | 31 505 480 | G | 0.6634 (0.5927-0.7427) | **1,02E-12** |
| IE1A | All subjects | 6 | RS3093974 | 31 506 210 | A | 0.8236 (0.7626-0.8895) | 7,80E-07 |
| IE1A | All subjects | 6 | RS2071592 | 31 515 340 | A | 0.8231 (0.7608-0.8906) | 1,28E-06 |
| IE1A | All subjects | 6 | RS909253 | 31 540 313 | G | 0.8398 (0.7777-0.907) | 8,63E-06 |
| IE1A | All subjects | 6 | RS1041981 | 31 540 784 | A | 0.8366 (0.7746-0.9035) | 5,55E-06 |

| | | | | | | | |
|---|---|---|---|---|---|---|---|
| IE1A | All subjects | 6 | RS1800629 | 31 543 031 | A | 0.7667 (0.6948-0.846) | 1,22E-07 |
| IE1A | All subjects | 6 | RS2857595 | 31 568 469 | A | 0.8081 (0.7377-0.8852) | 4,58E-06 |
| IE1A | All subjects | 6 | RS3132451 | 31 582 025 | C | 0.7966 (0.7257-0.8745) | 1,76E-06 |
| IE1A | All subjects | 6 | RS3130070 | 31 591 808 | G | 0.7811 (0.711-0.8581) | 2,63E-07 |
| IE1A | All subjects | 6 | RS3130622 | 31 592 524 | C | 0.7862 (0.7155-0.864) | 5,75E-07 |
| IE1A | All subjects | 6 | RS3130623 | 31 597 700 | A | 0.7987 (0.7285-0.8757) | 1,71E-06 |
| IE1A | All subjects | 6 | RS3130626 | 31 598 489 | G | 0.7818 (0.7115-0.859) | 3,02E-07 |
| IE1A | All subjects | 6 | RS2736157 | 31 600 820 | G | 0.7841 (0.7137-0.8614) | 4,02E-07 |
| IE1A | All subjects | 6 | RS3115663 | 31 601 843 | G | 0.7871 (0.7153-0.866) | 9,17E-07 |
| IE1A | All subjects | 6 | RS9267522 | 31 603 770 | G | 0.7805 (0.7104-0.8575) | 2,45E-07 |
| IE1A | All subjects | 6 | RS10885 | 31 604 591 | A | 0.7828 (0.7124-0.8602) | 3,54E-07 |
| IE1A | All subjects | 6 | RS3130628 | 31 609 272 | G | 0.7821 (0.7118-0.8593) | 3,13E-07 |
| IE1A | All subjects | 6 | RS3130048 | 31 613 739 | G | 0.8186 (0.752-0.8911) | 3,82E-06 |
| IE1A | All subjects | 6 | RS3117583 | 31 619 576 | G | 0.7798 (0.7098-0.8568) | 2,25E-07 |
| IE1A | All subjects | 6 | **RS3117582** | 31 620 520 | C | 0.6623 (0.5902-0.7431) | **2,35E-12** |
| IE1A | All subjects | 6 | **RS805262** | 31 628 733 | A | 0.7764 (0.7213-0.8358) | **1,65E-11** |
| IE1A | All subjects | 6 | RS3130618 | 31 632 134 | A | 0.7807 (0.7105-0.8579) | 2,68E-07 |
| IE1A | All subjects | 6 | **RS9267531** | 31 636 742 | G | 0.6629 (0.5907-0.7439) | **2,76E-12** |
| IE1A | All subjects | 6 | **RS3101018** | 31 705 864 | A | 0.6599 (0.5875-0.7411) | **2,27E-12** |
| IE1A | All subjects | 6 | **RS3132445** | 31 712 196 | A | 0.6645 (0.5922-0.7457) | **3,65E-12** |
| IE1A | All subjects | 6 | **RS3130484** | 31 715 882 | G | 0.6632 (0.591-0.7442) | **2,83E-12** |
| IE1A | All subjects | 6 | **RS3131379** | 31 721 033 | A | 0.6661 (0.5935-0.7476) | **5,21E-12** |
| IE1A | All subjects | 6 | **RS3117574** | 31 725 230 | A | 0.6648 (0.5924-0.746) | **3,83E-12** |
| IE1A | All subjects | 6 | **RS3131378** | 31 725 285 | G | 0.6623 (0.5902-0.7432) | **2,42E-12** |
| IE1A | All subjects | 6 | **RS3117577** | 31 727 474 | G | 0.6625 (0.5904-0.7435) | **2,54E-12** |
| IE1A | All subjects | 6 | **RS3101017** | 31 733 466 | G | 0.6646 (0.5923-0.7457) | **3,55E-12** |
| IE1A | All subjects | 6 | RS2075798 | 31 846 741 | A | 1.378 (1.206-1.575) | 2,52E-06 |
| IE1A | All subjects | 6 | RS555007 | 31 850 332 | G | 1.357 (1.189-1.55) | 6,56E-06 |
| IE1A | All subjects | 6 | **RS652888** | 31 851 234 | G | 0.7644 (0.6975-0.8376) | **8,69E-09** |
| IE1A | All subjects | 6 | **RS558702** | 31 870 326 | A | 0.6602 (0.5882-0.7409) | **1,79E-12** |
| IE1A | All subjects | 6 | **RS519417** | 31 878 433 | A | 0.6561 (0.5844-0.7365) | **9,34E-13** |
| IE1A | All subjects | 6 | RS511294 | 31 888 869 | C | 1.541 (1.302-1.823) | 4,90E-07 |
| IE1A | All subjects | 6 | **RS497309** | 31 892 484 | C | 0.6572 (0.5857-0.7375) | **9,47E-13** |
| IE1A | All subjects | 6 | RS498240 | 31 892 592 | A | 1.54 (1.301-1.823) | 5,29E-07 |
| IE1A | All subjects | 6 | RS2257331 | 31 898 285 | G | 1.539 (1.3-1.821) | 5,25E-07 |
| IE1A | All subjects | 6 | RS9332735 | 31 901 773 | A | 1.764 (1.412-2.203) | 5,72E-07 |
| IE1A | All subjects | 6 | RS635348 | 31 902 289 | C | 1.525 (1.289-1.805) | 9,11E-07 |
| IE1A | All subjects | 6 | RS621701 | 31 903 121 | A | 1.527 (1.29-1.807) | 8,26E-07 |
| IE1A | All subjects | 6 | **RS1042663** | 31 905 130 | A | 1.471 (1.283-1.686) | **2,96E-08** |
| IE1A | All subjects | 6 | **RS2507953** | 31 905 328 | A | 1.472 (1.284-1.687) | **2,81E-08** |
| IE1A | All subjects | 6 | **RS550605** | 31 907 147 | G | 1.466 (1.279-1.681) | **3,89E-08** |

| | | | | | | | |
|---|---|---|---|---|---|---|---|
| IE1A | All subjects | 6 | **RS653414** | 31 907 168 | A | 1.471 (1.283-1.686) | **2,95E-08** |
| IE1A | All subjects | 6 | **RS497239** | 31 908 761 | G | 1.468 (1.281-1.682) | **3,39E-08** |
| IE1A | All subjects | 6 | **RS609061** | 31 910 162 | A | 1.469 (1.281-1.683) | **3,29E-08** |
| IE1A | All subjects | 6 | **RS2242572** | 31 910 929 | A | 1.47 (1.283-1.685) | **3,06E-08** |
| IE1A | All subjects | 6 | **RS547154** | 31 910 938 | A | 1.469 (1.282-1.684) | **3,17E-08** |
| IE1A | All subjects | 6 | **RS541862** | 31 916 951 | G | 1.466 (1.279-1.68) | **3,96E-08** |
| IE1A | All subjects | 6 | **RS1270942** | 31 918 860 | G | 0.6563 (0.5849-0.7366) | **8,23E-13** |
| IE1A | All subjects | 6 | RS522162 | 31 919 917 | G | 1.455 (1.269-1.667) | 6,95E-08 |
| IE1A | All subjects | 6 | **RS760070** | 31 919 956 | G | 1.465 (1.278-1.679) | **4,23E-08** |
| IE1A | All subjects | 6 | **RS550513** | 31 920 687 | A | 1.466 (1.279-1.68) | **3,92E-08** |
| IE1A | All subjects | 6 | **RS403569** | 31 924 880 | A | 1.465 (1.278-1.679) | **4,17E-08** |
| IE1A | All subjects | 6 | **RS438999** | 31 928 306 | G | 1.459 (1.274-1.672) | **5,00E-08** |
| IE1A | All subjects | 6 | RS429608 | 31 930 462 | A | 1.297 (1.164-1.446) | 2,49E-06 |
| IE1A | All subjects | 6 | RS444921 | 31 932 177 | A | 1.314 (1.167-1.479) | 6,33E-06 |
| IE1A | All subjects | 6 | RS406936 | 31 933 161 | A | 1.313 (1.167-1.479) | 6,58E-06 |
| IE1A | All subjects | 6 | RS492899 | 31 933 518 | G | 1.41 (1.218-1.632) | 4,26E-06 |
| IE1A | All subjects | 6 | RS454212 | 31 934 372 | A | 1.315 (1.168-1.48) | 6,02E-06 |
| IE1A | All subjects | 6 | RS453821 | 31 935 311 | A | 1.312 (1.165-1.477) | 7,08E-06 |
| IE1A | All subjects | 6 | RS449643 | 31 936 679 | A | 1.315 (1.168-1.48) | 6,08E-06 |
| IE1A | All subjects | 6 | RS387608 | 31 941 557 | A | 1.312 (1.165-1.477) | 7,05E-06 |
| IE1A | All subjects | 6 | RS6455 | 32 006 896 | G | 1.699 (1.356-2.127) | 3,94E-06 |
| IE1A | All subjects | 6 | RS12333245 | 32 019 769 | A | 1.346 (1.185-1.53) | 5,06E-06 |
| IE1A | All subjects | 6 | RS11969759 | 32 021 130 | A | 1.447 (1.234-1.698) | 5,81E-06 |
| IE1A | All subjects | 6 | RS2269429 | 32 029 183 | A | 1.34 (1.179-1.523) | 7,74E-06 |
| IE1A | All subjects | 6 | **RS1150755** | 32 038 550 | A | 0.7034 (0.6358-0.7782) | **8,87E-12** |
| IE1A | All subjects | 6 | RS7774197 | 32 046 275 | C | 1.449 (1.235-1.699) | 5,31E-06 |
| IE1A | All subjects | 6 | **RS1150754** | 32 050 758 | A | 0.7022 (0.6347-0.7768) | **7,02E-12** |
| IE1A | All subjects | 6 | RS169496 | 32 052 983 | A | 1.341 (1.181-1.524) | 6,45E-06 |
| IE1A | All subjects | 6 | RS204900 | 32 056 580 | C | 1.343 (1.182-1.526) | 5,94E-06 |
| IE1A | All subjects | 6 | RS204899 | 32 057 627 | A | 1.348 (1.185-1.533) | 5,44E-06 |
| IE1A | All subjects | 6 | **RS1150753** | 32 059 867 | G | 0.6601 (0.5882-0.7408) | **1,66E-12** |
| IE1A | All subjects | 6 | **RS1150752** | 32 064 726 | G | 0.6521 (0.5806-0.7323) | **5,19E-13** |
| IE1A | All subjects | 6 | RS439844 | 32 072 940 | A | 1.342 (1.181-1.525) | 6,38E-06 |
| IE1A | All subjects | 6 | RS411337 | 32 077 380 | A | 1.416 (1.238-1.62) | 3,75E-07 |
| IE1A | All subjects | 6 | **RS1269852** | 32 080 191 | C | 0.6684 (0.5957-0.75) | **7,18E-12** |
| IE1A | All subjects | 6 | RS9469084 | 32 080 383 | A | 1.407 (1.23-1.609) | 6,35E-07 |
| IE1A | All subjects | 6 | RS204888 | 32 089 142 | A | 1.407 (1.231-1.61) | 6,01E-07 |
| IE1A | All subjects | 6 | **RS3130288** | 32 096 001 | A | 0.6665 (0.594-0.7478) | **4,96E-12** |
| IE1A | All subjects | 6 | RS169494 | 32 097 876 | A | 1.427 (1.247-1.632) | 2,27E-07 |
| IE1A | All subjects | 6 | **RS3134608** | 32 117 971 | C | 0.7463 (0.6787-0.8207) | **1,55E-09** |
| IE1A | All subjects | 6 | RS1053924 | 32 120 715 | A | 1.225 (1.134-1.323) | 2,27E-07 |

| | | | | | | | |
|---|---|---|---|---|---|---|---|
| IE1A | All subjects | 6 | RS3134950 | 32 127 477 | C | 1.223 (1.135-1.318) | 1,33E-07 |
| IE1A | All subjects | 6 | **RS3096697** | 32 134 510 | A | 0.7444 (0.6769-0.8186) | **1,17E-09** |
| IE1A | All subjects | 6 | **RS3130347** | 32 134 656 | G | 0.7465 (0.6788-0.8208) | **1,60E-09** |
| IE1A | All subjects | 6 | RS1061808 | 32 136 547 | A | 1.222 (1.134-1.317) | 1,43E-07 |
| IE1A | All subjects | 6 | **RS3130284** | 32 140 487 | G | 0.7456 (0.678-0.8199) | **1,41E-09** |
| IE1A | All subjects | 6 | RS408359 | 32 141 883 | A | 1.436 (1.231-1.675) | 4,15E-06 |
| IE1A | All subjects | 6 | **RS3134947** | 32 145 205 | A | 0.7465 (0.6788-0.8209) | **1,63E-09** |
| IE1A | All subjects | 6 | RS2269423 | 32 145 707 | A | 1.216 (1.129-1.31) | 2,65E-07 |
| IE1A | All subjects | 6 | **RS3134945** | 32 146 492 | A | 0.7465 (0.6788-0.8209) | **1,66E-09** |
| IE1A | All subjects | 6 | **RS3130349** | 32 147 696 | A | 0.7183 (0.6505-0.7931) | **5,97E-11** |
| IE1A | All subjects | 6 | **RS1800625** | 32 152 442 | G | 0.7189 (0.6511-0.7937) | **6,51E-11** |
| IE1A | All subjects | 6 | RS204995 | 32 154 285 | G | 0.7736 (0.7038-0.8503) | 1,03E-07 |
| IE1A | All subjects | 6 | **RS204993** | 32 155 581 | G | 0.7503 (0.6862-0.8203) | **2,85E-10** |
| IE1A | All subjects | 6 | **RS204992** | 32 156 908 | A | 0.7637 (0.6944-0.8399) | **2,79E-08** |
| IE1A | All subjects | 6 | **RS176095** | 32 158 319 | G | 0.7641 (0.6946-0.8407) | **3,30E-08** |
| IE1A | All subjects | 6 | RS204991 | 32 161 366 | G | 0.7577 (0.6854-0.8376) | 5,77E-08 |
| IE1A | All subjects | 6 | **RS204990** | 32 161 430 | A | 0.7506 (0.6785-0.8304) | **2,62E-08** |
| IE1A | All subjects | 6 | **RS2071278** | 32 165 444 | G | 0.709 (0.6375-0.7885) | **2,27E-10** |
| IE1A | All subjects | 6 | **RS3134942** | 32 168 771 | A | 0.6734 (0.6029-0.752) | **2,28E-12** |
| IE1A | All subjects | 6 | **RS2071287** | 32 170 433 | A | 0.7813 (0.7254-0.8416) | **7,37E-11** |
| IE1A | All subjects | 6 | **RS3132935** | 32 171 075 | G | 0.758 (0.6885-0.8346) | **1,66E-08** |
| IE1A | All subjects | 6 | **RS2071277** | 32 171 683 | G | 0.7775 (0.722-0.8372) | **2,58E-11** |
| IE1A | All subjects | 6 | **RS3131296** | 32 172 993 | A | 0.6716 (0.601-0.7504) | **2,08E-12** |
| IE1A | All subjects | 6 | **RS206017** | 32 176 211 | G | 1.286 (1.194-1.385) | **3,03E-11** |
| IE1A | All subjects | 6 | **RS206018** | 32 177 880 | C | 1.323 (1.203-1.456) | **8,98E-09** |
| IE1A | All subjects | 6 | **RS3132956** | 32 179 438 | A | 0.6741 (0.6036-0.7528) | **2,59E-12** |
| IE1A | All subjects | 6 | **RS3131290** | 32 183 175 | A | 0.742 (0.6871-0.8013) | **2,89E-14** |
| IE1A | All subjects | 6 | **RS3134799** | 32 184 221 | A | 0.7529 (0.6984-0.8116) | **1,30E-13** |
| IE1A | All subjects | 6 | **RS436388** | 32 186 264 | A | 1.296 (1.203-1.396) | **7,20E-12** |
| IE1A | All subjects | 6 | **RS379464** | 32 186 348 | A | 1.677 (1.404-2.005) | **1,26E-08** |
| IE1A | All subjects | 6 | **RS394657** | 32 187 023 | G | 0.757 (0.7023-0.8159) | **3,49E-13** |
| IE1A | All subjects | 6 | **RS1109771** | 32 187 605 | G | 0.7815 (0.7257-0.8416) | **6,99E-11** |
| IE1A | All subjects | 6 | **RS8192584** | 32 189 192 | A | 1.87 (1.513-2.312) | **6,86E-09** |
| IE1A | All subjects | 6 | RS715299 | 32 189 841 | C | 0.8144 (0.7481-0.8866) | 2,15E-06 |
| IE1A | All subjects | 6 | RS915895 | 32 190 217 | G | 0.8177 (0.7514-0.8898) | 3,07E-06 |
| IE1A | All subjects | 6 | **RS915894** | 32 190 390 | C | 0.7322 (0.6765-0.7924) | **1,13E-14** |
| IE1A | All subjects | 6 | RS443198 | 32 190 406 | G | 0.8129 (0.7488-0.8826) | 7,79E-07 |
| IE1A | All subjects | 6 | **RS3134930** | 32 191 620 | A | 0.7583 (0.6917-0.8313) | **3,64E-09** |
| IE1A | All subjects | 6 | RS377763 | 32 199 144 | A | 0.7955 (0.7253-0.8726) | 1,23E-06 |
| IE1A | All subjects | 6 | **RS6901158** | 32 205 942 | A | 0.675 (0.6069-0.7508) | **4,45E-13** |
| IE1A | All subjects | 6 | **RS424232** | 32 208 324 | A | 0.77 (0.706-0.8398) | **3,52E-09** |

| | | | | | | | |
|---|---|---|---|---|---|---|---|
| IE1A | All subjects | 6 | **RS3132971** | 32 230 256 | C | 0.6381 (0.5682-0.7165) | **3,08E-14** |
| IE1A | All subjects | 6 | **RS7775397** | 32 261 252 | C | 0.6425 (0.5723-0.7213) | **6,56E-14** |
| IE1A | All subjects | 6 | **RS9268219** | 32 284 108 | C | 0.6632 (0.5892-0.7463) | **9,56E-12** |
| IE1A | All subjects | 6 | **RS9268235** | 32 290 208 | A | 0.6438 (0.5734-0.7228) | **9,15E-14** |
| IE1A | All subjects | 6 | RS1265762 | 32 321 115 | C | 1.188 (1.103-1.28) | 5,63E-06 |
| IE1A | All subjects | 6 | RS1265760 | 32 321 872 | A | 1.19 (1.105-1.282) | 4,37E-06 |
| IE1A | All subjects | 6 | RS1265759 | 32 322 393 | G | 1.187 (1.102-1.279) | 6,78E-06 |
| IE1A | All subjects | 6 | RS6904636 | 32 327 781 | G | 1.183 (1.098-1.275) | 9,44E-06 |
| IE1A | All subjects | 6 | RS3129911 | 32 328 403 | G | 1.187 (1.102-1.279) | 6,38E-06 |
| IE1A | All subjects | 6 | RS3129922 | 32 333 098 | G | 1.192 (1.106-1.284) | 3,87E-06 |
| IE1A | All subjects | 6 | **RS2273017** | 32 337 630 | G | 1.234 (1.145-1.331) | **4,36E-08** |
| IE1A | All subjects | 6 | **RS2073045** | 32 339 548 | A | 0.7661 (0.7083-0.8286) | **2,80E-11** |
| IE1A | All subjects | 6 | **RS7758128** | 32 345 283 | A | 1.813 (1.476-2.226) | **1,35E-08** |
| IE1A | All subjects | 6 | **RS2894254** | 32 345 689 | C | 0.6436 (0.5728-0.7231) | **1,21E-13** |
| IE1A | All subjects | 6 | RS17423649 | 32 357 133 | A | 1.299 (1.17-1.442) | 9,23E-07 |
| IE1A | All subjects | 6 | RS17202259 | 32 357 489 | C | 1.298 (1.169-1.442) | 9,88E-07 |
| IE1A | All subjects | 6 | RS3117099 | 32 358 270 | A | 0.8188 (0.7495-0.8944) | 9,28E-06 |
| IE1A | All subjects | 6 | RS16870123 | 32 359 460 | A | 1.293 (1.165-1.435) | 1,35E-06 |
| IE1A | All subjects | 6 | RS3817969 | 32 361 388 | A | 1.281 (1.155-1.421) | 2,86E-06 |
| IE1A | All subjects | 6 | RS1980493 | 32 363 215 | G | 0.794 (0.7194-0.8763) | 4,59E-06 |
| IE1A | All subjects | 6 | RS2227138 | 32 384 500 | A | 0.7787 (0.7042-0.8612) | 1,11E-06 |
| IE1A | All subjects | 6 | RS3135353 | 32 392 877 | A | 0.7766 (0.7022-0.8589) | 8,58E-07 |
| IE1A | All subjects | 6 | **RS3129843** | 32 395 726 | G | 0.6484 (0.5772-0.7285) | **3,01E-13** |
| IE1A | All subjects | 6 | **RS3763326** | 32 413 557 | G | 1.814 (1.483-2.218) | **6,62E-09** |
| IE1A | All subjects | 6 | **RS9268877** | 32 431 147 | A | 1.234 (1.145-1.331) | **4,48E-08** |
| IE1A | All subjects | 6 | RS9268979 | 32 435 044 | A | 1.23 (1.141-1.328) | 8,55E-08 |
| IE1A | All subjects | 6 | **RS9270665** | 32 566 232 | A | 0.734 (0.6679-0.8067) | **1,34E-10** |
| IE1A | All subjects | 6 | **RS642093** | 32 582 075 | A | 0.7452 (0.6798-0.817) | **3,71E-10** |
| IE1A | All subjects | 6 | **RS3129763** | 32 590 925 | A | 0.7349 (0.6689-0.8075) | **1,49E-10** |
| IE1A | All subjects | 6 | **RS9272219** | 32 602 269 | A | 0.7481 (0.6871-0.8146) | **2,39E-11** |
| IE1A | All subjects | 6 | RS3104369 | 32 602 482 | A | 1.205 (1.116-1.301) | 2,06E-06 |
| IE1A | All subjects | 6 | **RS2187668** | 32 605 884 | A | 0.7017 (0.6285-0.7836) | **3,08E-10** |
| IE1A | All subjects | 6 | **RS9273012** | 32 611 641 | G | 0.7461 (0.6853-0.8123) | **1,48E-11** |
| IE1A | All subjects | 6 | **RS17843604** | 32 620 283 | A | 0.7302 (0.6761-0.7885) | **1,05E-15** |
| IE1A | All subjects | 6 | **RS9273327** | 32 623 223 | C | 0.7015 (0.6278-0.7839) | **3,94E-10** |
| IE1A | All subjects | 6 | **RS9273349** | 32 625 869 | G | 0.7042 (0.652-0.7606) | **4,70E-19** |
| IE1A | All subjects | 6 | RS6928482 | 32 626 249 | G | 0.833 (0.7697-0.9016) | 6,04E-06 |
| IE1A | All subjects | 6 | **RS9273363** | 32 626 272 | A | 0.7238 (0.667-0.7855) | **9,43E-15** |
| IE1A | All subjects | 6 | **RS6906021** | 32 626 311 | G | 0.7653 (0.7074-0.828) | **2,75E-11** |
| IE1A | All subjects | 6 | **RS1063355** | 32 627 714 | C | 0.7038 (0.6516-0.7602) | **4,40E-19** |
| IE1A | All subjects | 6 | **RS9273448** | 32 627 747 | A | 1.327 (1.227-1.435) | **1,66E-12** |

| | | | | | | | |
|---|---|---|---|---|---|---|---|
| IE1A | All subjects | 6 | **RS2854275** | 32 628 428 | A | 0.6983 (0.6255-0.7797) | **1,69E-10** |
| IE1A | All subjects | 6 | RS2647025 | 32 635 949 | A | 0.8251 (0.758-0.8981) | 8,93E-06 |
| IE1A | All subjects | 6 | RS9275141 | 32 651 117 | C | 0.8158 (0.7544-0.8823) | 3,46E-07 |
| IE1A | All subjects | 6 | RS2856695 | 32 651 894 | G | 0.8165 (0.755-0.883) | 3,84E-07 |
| IE1A | All subjects | 6 | RS3021058 | 32 652 359 | C | 0.8145 (0.7532-0.8808) | 2,78E-07 |
| IE1A | All subjects | 6 | RS4947342 | 32 653 070 | A | 0.8075 (0.7389-0.8825) | 2,36E-06 |
| IE1A | All subjects | 6 | RS2856692 | 32 653 385 | C | 0.8044 (0.736-0.8792) | 1,60E-06 |
| IE1A | All subjects | 6 | RS4642516 | 32 657 543 | A | 0.8128 (0.7516-0.879) | 2,13E-07 |
| IE1A | All subjects | 6 | RS5000634 | 32 663 564 | G | 0.8188 (0.7557-0.8872) | 1,03E-06 |
| IE1A | All subjects | 6 | RS6457622 | 32 664 163 | C | 0.8163 (0.7535-0.8844) | 6,83E-07 |
| IE1A | All subjects | 6 | RS9275312 | 32 665 728 | G | 0.8021 (0.7282-0.8835) | 7,78E-06 |
| IE1A | All subjects | 6 | **RS1794282** | 32 666 526 | A | 0.6492 (0.5777-0.7295) | **3,85E-13** |
| IE1A | All subjects | 6 | RS9275328 | 32 666 822 | A | 0.8035 (0.7294-0.8851) | 9,21E-06 |
| IE1A | All subjects | 6 | RS9275330 | 32 666 875 | G | 0.8037 (0.7296-0.8853) | 9,51E-06 |
| IE1A | All subjects | 6 | RS9275333 | 32 666 968 | G | 0.8018 (0.7279-0.8831) | 7,46E-06 |
| IE1A | All subjects | 6 | **RS3135006** | 32 667 119 | A | 1.33 (1.229-1.439) | **1,40E-12** |
| IE1A | All subjects | 6 | **RS4947344** | 32 677 846 | A | 1.283 (1.189-1.384) | **1,23E-10** |
| IE1A | All subjects | 6 | RS9275601 | 32 682 664 | A | 0.8145 (0.755-0.8786) | 1,13E-07 |
| IE1A | All subjects | 6 | RS3873444 | 32 682 724 | A | 1.367 (1.202-1.555) | 2,04E-06 |
| IE1A | All subjects | 6 | RS2857193 | 32 747 528 | G | 1.188 (1.102-1.281) | 6,69E-06 |
| IE1A | All subjects | 6 | RS28986337 | 32 748 398 | A | 1.381 (1.206-1.582) | 3,02E-06 |
| IE1A | All subjects | 6 | RS28986338 | 32 748 810 | A | 1.386 (1.211-1.587) | 2,33E-06 |
| IE1A | All subjects | 6 | RS28986359 | 32 751 349 | C | 1.379 (1.204-1.58) | 3,64E-06 |
| IE1A | All subjects | 6 | RS28986364 | 32 751 691 | C | 1.383 (1.207-1.585) | 2,97E-06 |
| IE1A | All subjects | 6 | **RS9276689** | 32 751 962 | A | 0.6444 (0.57-0.7285) | **2,19E-12** |
| IE1A | All subjects | 6 | RS28986373 | 32 752 150 | A | 1.383 (1.208-1.584) | 2,71E-06 |
| IE1A | All subjects | 6 | RS13204139 | 32 752 685 | A | 1.374 (1.2-1.573) | 4,33E-06 |
| IE1A | All subjects | 6 | RS2857177 | 32 752 840 | A | 1.187 (1.102-1.28) | 7,31E-06 |
| IE1A | All subjects | 6 | RS28986396 | 32 754 430 | A | 1.379 (1.205-1.58) | 3,25E-06 |
| IE1A | All subjects | 6 | **RS7762279** | 32 755 290 | G | 0.6429 (0.5689-0.7265) | **1,41E-12** |
| IE1A | All subjects | 6 | RS2857165 | 32 757 737 | G | 1.19 (1.104-1.283) | 5,46E-06 |
| IE1A | All subjects | 6 | RS2857164 | 32 758 219 | C | 1.191 (1.105-1.284) | 4,64E-06 |
| IE1A | All subjects | 6 | RS1158785 | 32 758 571 | A | 1.193 (1.107-1.286) | 4,09E-06 |
| IE1A | All subjects | 6 | RS2621384 | 32 759 273 | G | 1.192 (1.106-1.284) | 4,61E-06 |
| IE1A | All subjects | 6 | RS2857161 | 32 759 297 | G | 1.194 (1.108-1.287) | 3,53E-06 |
| IE1A | All subjects | 6 | RS9276722 | 32 762 391 | G | 1.381 (1.207-1.581) | 2,86E-06 |
| IE1A | All subjects | 6 | RS9276726 | 32 763 888 | G | 0.8219 (0.7602-0.8887) | 8,53E-07 |
| IE1A | All subjects | 6 | RS1383261 | 32 765 451 | A | 0.8353 (0.7728-0.9029) | 5,78E-06 |
| IE1A | All subjects | 6 | RS6932969 | 32 765 671 | G | 0.833 (0.7708-0.9003) | 3,99E-06 |
| IE1A | All subjects | 6 | RS7383606 | 32 766 593 | A | 0.8341 (0.7718-0.9014) | 4,61E-06 |
| IE1A | All subjects | 6 | RS9276734 | 32 766 854 | A | 0.8336 (0.7714-0.9009) | 4,36E-06 |

| | | | | | | | |
|---|---|---|---|---|---|---|---|
| IE1A | All subjects | 6 | RS7382714 | 32 767 496 | G | 0.8339 (0.7716-0.9012) | 4,51E-06 |
| IE1A | All subjects | 6 | **RS4947350** | 32 767 620 | G | 0.7622 (0.6978-0.8325) | **1,61E-09** |
| IE1A | All subjects | 6 | RS7381376 | 32 767 673 | G | 0.8346 (0.7721-0.9021) | 5,26E-06 |
| IE1A | All subjects | 6 | RS6899857 | 32 770 482 | G | 0.8345 (0.7723-0.9018) | 4,82E-06 |
| IE1A | All subjects | 6 | RS6912414 | 32 774 465 | A | 0.8218 (0.7602-0.8885) | 8,14E-07 |
| IE1A | All subjects | 6 | RS7382619 | 32 775 809 | A | 0.8348 (0.7725-0.9022) | 5,16E-06 |
| IE1A | All subjects | 6 | RS7382649 | 32 776 087 | A | 0.8343 (0.772-0.9016) | 4,76E-06 |
| IE1A | All subjects | 6 | **RS11244** | 32 780 724 | A | 0.782 (0.7192-0.8503) | **8,77E-09** |
| IE1A | All subjects | 6 | RS13209654 | 32 792 659 | G | 1.389 (1.213-1.591) | 2,03E-06 |
| IE1A | All subjects | 6 | RS1015166 | 32 798 731 | A | 0.8312 (0.7696-0.8977) | 2,52E-06 |
| IE1A | All subjects | 6 | RS3819714 | 32 804 217 | A | 1.238 (1.144-1.34) | 1,20E-07 |
| IE1A | All subjects | 6 | RS3819715 | 32 804 219 | A | 1.235 (1.141-1.336) | 1,62E-07 |
| IE1A | All subjects | 6 | RS241427 | 32 804 414 | A | 1.228 (1.14-1.323) | 7,26E-08 |
| IE1A | All subjects | 6 | RS241426 | 32 804 553 | A | 1.226 (1.139-1.321) | 7,09E-08 |
| IE1A | All subjects | 6 | **RS3819720** | 32 804 570 | A | 0.748 (0.6912-0.8096) | **6,23E-13** |
| IE1A | All subjects | 6 | **RS241425** | 32 804 909 | A | 1.273 (1.183-1.371) | **1,43E-10** |
| IE1A | All subjects | 6 | **RS241424** | 32 804 934 | A | 0.8129 (0.7549-0.8753) | **4,06E-08** |
| IE1A | All subjects | 6 | **RS2239701** | 32 805 049 | G | 0.7802 (0.7239-0.8409) | **8,37E-11** |
| IE1A | All subjects | 6 | RS2071465 | 32 805 470 | C | 0.8217 (0.7617-0.8864) | 3,88E-07 |
| IE1A | All subjects | 6 | RS4148870 | 32 806 391 | A | 0.839 (0.7785-0.9043) | 4,40E-06 |
| IE1A | All subjects | 6 | RS2071552 | 32 806 461 | G | 0.8352 (0.775-0.9001) | 2,42E-06 |
| IE1A | All subjects | 6 | RS4148868 | 32 806 584 | A | 0.8345 (0.7742-0.8995) | 2,28E-06 |
| IE1A | All subjects | 6 | RS4713598 | 32 806 786 | C | 0.8351 (0.7744-0.9006) | 2,86E-06 |
| IE1A | All subjects | 6 | RS3763366 | 32 807 446 | C | 0.8316 (0.7718-0.896) | 1,26E-06 |
| IE1A | All subjects | 6 | RS3763349 | 32 808 232 | G | 0.8335 (0.7736-0.898) | 1,70E-06 |
| IE1A | All subjects | 6 | RS9276810 | 32 810 443 | A | 0.83 (0.7702-0.8944) | 1,02E-06 |
| IE1A | All subjects | 6 | RS6924102 | 32 811 383 | G | 0.8378 (0.777-0.9034) | 4,15E-06 |
| IE1A | All subjects | 6 | **RS1480380** | 32 913 246 | A | 0.6676 (0.5841-0.763) | **3,02E-09** |
| IE1A | All subjects | 6 | **RS9276933** | 32 930 795 | G | 0.6931 (0.6095-0.7882) | **2,31E-08** |
| IE1A | All subjects | 6 | RS423196 | 33 003 562 | G | 0.7726 (0.6948-0.8592) | 1,95E-06 |
| IE1A | All subjects | 6 | RS406477 | 33 005 644 | G | 0.7699 (0.6928-0.8556) | 1,21E-06 |
| IE1A | All subjects | 6 | RS987870 | 33 042 880 | G | 0.7473 (0.6606-0.8453) | 3,61E-06 |
| IE1A | All subjects | 6 | RS2071354 | 33 044 388 | G | 0.7475 (0.6608-0.8455) | 3,66E-06 |
| IE1A | All subjects | 6 | **RS6914616** | 33 063 592 | A | 0.582 (0.4919-0.6886) | **2,82E-10** |
| IE1A | All subjects | 6 | **RS2179915** | 33 065 734 | A | 0.5821 (0.492-0.6887) | **2,86E-10** |
| IE1A | All subjects | 6 | RS3128930 | 33 075 666 | A | 0.8228 (0.7559-0.8956) | 6,58E-06 |
| IE1A | All subjects | 6 | **RS6924545** | 33 077 607 | A | 0.5928 (0.5011-0.7013) | **1,07E-09** |
| IE1A | All subjects | 6 | RS12529876 | 167 461 501 | A | 1.185 (1.099-1.277) | 8,94E-06 |
| IE1A | All subjects | 6 | RS4710181 | 167 513 998 | G | 1.183 (1.099-1.273) | 7,74E-06 |
| IE1A | All subjects | 6 | RS6907666 | 167 523 395 | A | 1.192 (1.106-1.285) | 4,72E-06 |
| IE1A | All subjects | 10 | RS4132050 | 12 797 758 | A | 0.8261 (0.7647-0.8924) | 1,22E-06 |

| | | | | | | | |
|---|---|---|---|---|---|---|---|
| IE1A | All subjects | 10 | RS2147268 | 105 793 459 | G | 0.8196 (0.7504-0.8952) | 9,99E-06 |
| IE1A | All subjects | 13 | RS2991478 | 79 754 928 | A | 0.7325 (0.6439-0.8333) | 2,23E-06 |
| IE1A | All subjects | 13 | RS2988030 | 79 762 645 | A | 0.7292 (0.6411-0.8293) | 1,52E-06 |
| IE1A | All subjects | 14 | RS7157970 | 105 463 936 | A | 1.196 (1.105-1.295) | 9,85E-06 |
| IE1A | All subjects | 18 | RS11872992 | 58 040 587 | A | 0.7858 (0.7094-0.8704) | 3,82E-06 |
| IE1A | All subjects | 22 | RS5753103 | 30 768 777 | A | 0.8411 (0.7812-0.9055) | 4,28E-06 |
| IE1A | All subjects | 22 | RS4820845 | 30 800 338 | C | 0.8291 (0.7699-0.8929) | 7,29E-07 |
| IE1A | All subjects | 22 | RS5997641 | 30 809 342 | A | 0.8251 (0.7602-0.8956) | 4,33E-06 |
| IE1A | All subjects | 22 | RS1061660 | 30 819 628 | G | 0.827 (0.7623-0.8973) | 5,05E-06 |
| IE1B | MS cases | 3 | RS7628773 | 24723928 | A | 1.285 (1.15-1.436) | 9,12E-06 |
| IE1B | MS cases | 3 | RS7629359 | 24727377 | G | 1.293 (1.157-1.445) | 5,84E-06 |
| IE1B | MS cases | 10 | RS1466196 | 130749975 | A | 1.267 (1.142-1.405) | 7,50E-06 |
| IE1B | Controls | 1 | RS6424074 | 3 317 780 | G | 0.7713 (0.6887-0.8639) | 7,08E-06 |
| IE1B | Controls | 14 | RS10149279 | 80 579 214 | C | 1.332 (1.179-1.504) | 3,77E-06 |
| IE1B | Controls | 15 | RS17114996 | 25 245 502 | A | 2.219 (1.559-3.158) | 9,52E-06 |
| IE1B | Controls | 15 | RS8038349 | 25 270 440 | A | 2.226 (1.561-3.173) | 9,81E-06 |
| IE1B | Controls | 15 | RS17115045 | 25 272 862 | G | 2.221 (1.561-3.161) | 9,28E-06 |
| IE1B | Controls | 15 | RS8035400 | 25 273 478 | A | 2.221 (1.56-3.16) | 9,34E-06 |
| IE1B | Controls | 15 | RS17115126 | 25 303 191 | G | 2.221 (1.561-3.161) | 9,28E-06 |
| IE1B | Controls | 15 | RS7171937 | 25 329 547 | A | 2.218 (1.559-3.157) | 9,58E-06 |
| IE1B | Controls | 19 | RS4926207 | 14 473 036 | G | 0.7728 (0.6905-0.865) | 7,44E-06 |
| IE1B | Controls | 20 | RS6089729 | 60 890 564 | A | 0.7377 (0.6447-0.8443) | 9,85E-06 |
| IE1B | All subjects | 6 | RS2516454 | 31420863 | T | 1.537 (1.271-1.86) | 9,74E-06 |
| IE1B | All subjects | 6 | RS2534678 | 31463963 | A | 1.399 (1.227-1.594) | 5,00E-07 |
| IE1B | All subjects | 6 | RS2534667 | 31468099 | A | 1.427 (1.239-1.643) | 8,16E-07 |
| IE1B | All subjects | 6 | RS2534657 | 31472459 | A | 1.401 (1.228-1.598) | 5,12E-07 |
| IE1B | All subjects | 6 | RS2844496 | 31480398 | A | 1.466 (1.259-1.707) | 8,07E-07 |
| IE1B | All subjects | 6 | RS2246986 | 31482203 | G | 1.47 (1.263-1.711) | 6,47E-07 |
| IE1B | All subjects | 6 | RS2844509 | 31510924 | G | 1.297 (1.168-1.439) | 9,79E-07 |
| IE1B | All subjects | 6 | RS2239705 | 31513402 | A | 1.313 (1.168-1.476) | 5,17E-06 |
| IE1B | All subjects | 6 | RS511027 | 32206687 | A | 1.432 (1.244-1.649) | 6,01E-07 |
| IE1B | All subjects | 6 | RS9268557 | 32389305 | G | 0.84 (0.778-0.9069) | 8,24E-06 |
| IE1B | All subjects | 15 | RS2397382 | 95523004 | A | 0.8132 (0.746-0.8864) | 2,60E-06 |
| IE1B | All subjects | 22 | RS442656 | 21416442 | A | 0.7812 (0.7016-0.8697) | 6,60E-06 |
| 101K | MS cases | 5 | RS1347135 | 167484417 | G | 1.302 (1.16-1.461) | 7,47E-06 |
| 101K | Controls | 5 | RS966297 | 23878207 | A | 0.7551 (0.6671-0.8547) | 8,87E-06 |

| | | | | | | | |
|---|---|---|---|---|---|---|---|
| 101K | Controls | 6 | RS2844571 | 31335647 | G | 0.7567 (0.6739-0.8496) | 2,39E-06 |
| 101K | Controls | 6 | RS2253908 | 31336891 | A | 0.7571 (0.674-0.8503) | 2,65E-06 |
| 101K | Controls | 6 | RS2844545 | 31345229 | A | 0.7659 (0.6832-0.8586) | 4,80E-06 |
| 101K | Controls | 6 | RS2269475 | 31583931 | A | 1.429 (1.229-1.662) | 3,66E-06 |
| 101K | Controls | 6 | RS3763295 | 31587938 | G | 1.456 (1.249-1.696) | 1,46E-06 |
| 101K | Controls | 6 | RS2242657 | 31602489 | G | 1.409 (1.212-1.638) | 8,17E-06 |
| 101K | Controls | 6 | RS2295665 | 31632686 | A | 1.427 (1.227-1.66) | 3,95E-06 |
| 101K | Controls | 6 | RS2295664 | 31633165 | A | 1.417 (1.219-1.647) | 5,84E-06 |
| 101K | Controls | 6 | RS14365 | 31635710 | G | 1.424 (1.235-1.643) | 1,25E-06 |
| 101K | Controls | 6 | RS2280800 | 31646398 | A | 1.41 (1.213-1.64) | 8,00E-06 |
| 101K | Controls | 6 | RS2242653 | 31675765 | A | 1.407 (1.215-1.631) | 5,36E-06 |
| 101K | Controls | 6 | RS2293861 | 31711124 | A | 1.441 (1.235-1.681) | 3,51E-06 |
| 101K | Controls | 6 | RS2075788 | 31712181 | C | 1.439 (1.233-1.679) | 3,81E-06 |
| 101K | Controls | 6 | RS3828922 | 31713454 | A | 1.44 (1.234-1.68) | 3,72E-06 |
| 101K | Controls | 6 | RS6905572 | 31731881 | A | 1.436 (1.224-1.684) | 8,83E-06 |
| 101K | Controls | 6 | RS2294878 | 32367795 | A | 1.316 (1.178-1.471) | 1,25E-06 |
| 101K | Controls | 6 | RS7774434 | 32657578 | G | 1.299 (1.159-1.456) | 7,09E-06 |
| 101K | Controls | 6 | RS9275428 | 32670978 | G | 1.317 (1.166-1.487) | 9,19E-06 |
| 101K | Controls | 15 | RS11637194 | 80494256 | G | 0.7656 (0.6822-0.8592) | 5,62E-06 |
| 101K | Controls | 15 | RS2028119 | 80514164 | C | 0.7787 (0.6973-0.8696) | 9,05E-06 |
| 101K | Controls | 20 | RS775121 | 16046227 | A | 0.7348 (0.6412-0.842) | 9,26E-06 |
| 101K | Controls | 20 | RS1467704 | 55441072 | G | 0.742 (0.6508-0.8459) | 8,17E-06 |
| 101K | All subjects | 6 | RS3116798 | 29671991 | G | 1.227 (1.122-1.343) | 7,36E-06 |
| 101K | All subjects | 6 | RS1611364 | 29679602 | A | 1.241 (1.135-1.357) | 2,19E-06 |
| 101K | All subjects | 6 | RS1362126 | 29691019 | A | 0.8418 (0.7804-0.908) | 8,38E-06 |
| 101K | All subjects | 6 | RS2523402 | 29699156 | G | 0.8395 (0.7784-0.9054) | 5,71E-06 |
| 101K | All subjects | 6 | RS2394159 | 29702679 | G | 0.8374 (0.7762-0.9034) | 4,61E-06 |
| 101K | All subjects | 6 | RS2735051 | 29705688 | A | 0.8394 (0.7782-0.9053) | 5,66E-06 |
| 101K | All subjects | 6 | RS2735048 | 29733701 | A | 0.8357 (0.7724-0.9043) | 8,14E-06 |
| 101K | All subjects | 6 | RS2735046 | 29734098 | A | 0.8379 (0.7752-0.9055) | 8,04E-06 |
| 101K | All subjects | 6 | RS2523409 | 29775662 | G | 0.8316 (0.769-0.8993) | 3,87E-06 |
| 101K | All subjects | 6 | RS3128912 | 29815637 | A | 1.254 (1.137-1.383) | 5,88E-06 |
| 101K | All subjects | 6 | RS2523822 | 29828660 | G | 0.8293 (0.7648-0.8993) | 6,04E-06 |
| 101K | All subjects | 6 | RS3094170 | 29829265 | A | 1.252 (1.136-1.381) | 6,64E-06 |
| 101K | All subjects | 6 | RS3094165 | 29833541 | A | 1.214 (1.12-1.315) | 2,34E-06 |
| 101K | All subjects | 6 | RS2860580 | 29906691 | A | 1.185 (1.1-1.276) | 8,09E-06 |
| 101K | All subjects | 6 | RS1616549 | 29914270 | A | 1.255 (1.138-1.384) | 5,19E-06 |
| 101K | All subjects | 6 | RS3823355 | 29942083 | A | 0.8322 (0.7676-0.9023) | 8,38E-06 |
| 101K | All subjects | 6 | RS3823358 | 29942205 | A | 0.829 (0.7643-0.8991) | 6,01E-06 |
| 101K | All subjects | 6 | RS6904029 | 29943067 | A | 0.8314 (0.7668-0.9013) | 7,49E-06 |

| | | | | | | | |
|---|---|---|---|---|---|---|---|
| 101K | All subjects | 6 | RS4713281 | 29978352 | A | 0.8301 (0.7653-0.9004) | 7,16E-06 |
| 101K | All subjects | 6 | RS9357092 | 29984252 | A | 0.8264 (0.7619-0.8963) | 4,21E-06 |
| 101K | All subjects | 6 | RS3910312 | 30008746 | C | 0.8299 (0.7651-0.9001) | 6,87E-06 |
| 101K | All subjects | 6 | RS9380150 | 30010492 | G | 0.8317 (0.7668-0.902) | 8,64E-06 |
| 101K | All subjects | 6 | RS9295829 | 30028800 | G | 0.8308 (0.766-0.9011) | 7,78E-06 |
| 101K | All subjects | 6 | RS3807031 | 30033884 | A | 0.8121 (0.7434-0.887) | 3,82E-06 |
| 101K | All subjects | 6 | RS9393989 | 30040084 | A | 0.83 (0.7652-0.9003) | 7,09E-06 |
| 101K | All subjects | 6 | RS4959041 | 30077967 | G | 0.8227 (0.7586-0.8923) | 2,43E-06 |
| 101K | All subjects | 6 | RS9261438 | 30089280 | A | 0.8258 (0.7617-0.8952) | 3,39E-06 |
| 101K | All subjects | 6 | RS9261639 | 30237859 | A | 0.8275 (0.761-0.8997) | 9,34E-06 |
| 101K | All subjects | 6 | RS2844571 | 31335647 | G | 0.8282 (0.7648-0.8969) | 3,53E-06 |
| 101K | All subjects | 6 | RS2253908 | 31336891 | A | 0.8289 (0.7653-0.8978) | 4,05E-06 |
| 101K | All subjects | 6 | RS9266629 | 31346822 | G | 0.7942 (0.7246-0.8705) | 8,63E-07 |
| 101K | All subjects | 6 | RS7741091 | 31352631 | G | 1.212 (1.115-1.318) | 6,07E-06 |
| 101K | All subjects | 6 | RS2516448 | 31390410 | G | 1.181 (1.097-1.272) | 9,34E-06 |
| 101K | All subjects | 6 | RS3763295 | 31587938 | G | 1.266 (1.141-1.405) | 8,36E-06 |
| 101K | All subjects | 6 | RS2295665 | 31632686 | A | 1.263 (1.14-1.401) | 8,69E-06 |
| 101K | All subjects | 6 | RS2295664 | 31633165 | A | 1.261 (1.138-1.398) | 9,98E-06 |
| 101K | All subjects | 6 | RS2293861 | 31711124 | A | 1.277 (1.15-1.419) | 5,05E-06 |
| 101K | All subjects | 6 | RS2075788 | 31712181 | C | 1.275 (1.148-1.416) | 5,76E-06 |
| 101K | All subjects | 6 | RS3828922 | 31713454 | A | 1.276 (1.149-1.418) | 5,29E-06 |
| 101K | All subjects | 6 | RS481825 | 31780594 | A | 0.6587 (0.5533-0.7842) | 2,71E-06 |
| 101K | All subjects | 6 | RS35294112 | 31794443 | G | 0.5835 (0.471-0.7229) | 8,25E-07 |
| 101K | All subjects | 6 | RS17201248 | 31803130 | A | 0.5826 (0.4706-0.7213) | 7,06E-07 |
| 101K | All subjects | 6 | RS391755 | 32192436 | G | 0.6883 (0.588-0.8057) | 3,32E-06 |
| 101K | All subjects | 6 | RS397081 | 32192617 | G | 0.6277 (0.5258-0.7493) | 2,56E-07 |
| 101K | All subjects | 6 | RS2294878 | 32367795 | A | 1.203 (1.115-1.299) | 2,06E-06 |
| 101K | All subjects | 6 | RS660895 | 32577380 | G | 1.245 (1.136-1.364) | 2,98E-06 |
| 101K | All subjects | 6 | RS7774434 | 32657578 | G | 1.207 (1.116-1.306) | 2,90E-06 |
| 101K | All subjects | 6 | RS9275245 | 32660943 | G | 1.192 (1.104-1.287) | 6,40E-06 |
| 101K | All subjects | 6 | RS6457617 | 32663851 | A | 1.207 (1.119-1.302) | 1,15E-06 |
| 101K | All subjects | 6 | RS9275312 | 32665728 | G | 1.264 (1.145-1.394) | 3,15E-06 |
| 101K | All subjects | 6 | RS9275328 | 32666822 | A | 1.267 (1.148-1.398) | 2,51E-06 |
| 101K | All subjects | 6 | RS9275330 | 32666875 | G | 1.266 (1.147-1.396) | 2,77E-06 |
| 101K | All subjects | 6 | RS9275333 | 32666968 | G | 1.262 (1.143-1.392) | 3,73E-06 |
| 101K | All subjects | 6 | RS9275371 | 32668296 | G | 1.254 (1.153-1.364) | 1,26E-07 |
| 101K | All subjects | 6 | RS9275374 | 32668526 | A | 1.256 (1.155-1.366) | 9,54E-08 |
| 101K | All subjects | 6 | RS9275388 | 32669084 | G | 1.255 (1.154-1.365) | 1,06E-07 |
| 101K | All subjects | 6 | RS9275390 | 32669156 | G | 1.258 (1.157-1.369) | 8,23E-08 |
| 101K | All subjects | 6 | RS9275393 | 32669439 | A | 1.255 (1.153-1.365) | 1,28E-07 |
| 101K | All subjects | 6 | RS9275406 | 32669955 | A | 1.255 (1.154-1.364) | 1,18E-07 |

| | | | | | | | |
|---|---|---|---|---|---|---|---|
| 101K | All subjects | 6 | RS9275407 | 32670037 | A | 1.255 (1.154-1.365) | 1,08E-07 |
| 101K | All subjects | 6 | RS9275418 | 32670244 | G | 1.255 (1.154-1.365) | 1,07E-07 |
| 101K | All subjects | 6 | RS9275424 | 32670576 | G | 1.256 (1.155-1.366) | 1,01E-07 |
| 101K | All subjects | 6 | RS9275428 | 32670978 | G | 1.256 (1.155-1.366) | 1,08E-07 |
| 101K | All subjects | 6 | RS9275439 | 32671521 | G | 1.255 (1.154-1.366) | 1,16E-07 |
| 101K | All subjects | 8 | RS900896 | 68528197 | G | 1.21 (1.112-1.317) | 9,59E-06 |
| 101K | All subjects | 12 | RS11170018 | 52550467 | A | 1.236 (1.126-1.356) | 8,05E-06 |
| 101K | All subjects | 20 | RS6097106 | 36589678 | A | 0.6993 (0.6004-0.8143) | 4,18E-06 |
